# Chromosome breakage-replication/fusion enables rapid DNA amplification

**DOI:** 10.1101/2024.08.17.608415

**Authors:** Cheng-Zhong Zhang, David Pellman

**Affiliations:** Department of Data Sciences, Dana-Farber Cancer Institute, Boston, MA, USA; Department of Pathology, Brigham and Women’s Hospital, Boston, MA, USA; Cancer Program, Broad Institute of MIT and Harvard, Cambridge, MA, USA; Department of Pediatric Oncology, Dana-Farber Cancer Institute, Boston, MA, USA; Department of Cell Biology, Blavatnik Institute, Harvard Medical School, Boston, MA, USA; Howard Hughes Medical Institute, Chevy Chase, MD, USA

## Abstract

DNA rearrangements are thought to arise from two classes of processes. The first class involves DNA breakage and fusion (“cut-and-paste”) without net DNA gain or loss. The second class involves aberrant DNA replication (“copy-and-paste”) and can produce either net DNA gain or loss. We previously demonstrated that the partitioning of chromosomes into aberrant structures of the nucleus, micronuclei or chromosome bridges, can generate cut-and-paste rearrangements by chromosome fragmentation and ligation. Surprisingly, in the progeny clones of single cells that have undergone chromosome bridge breakage, we identified large segmental duplications and short sequence insertions that are commonly attributed to copy-and-paste processes. Here, we demonstrate that both large duplications and short insertions are inherent outcomes of the replication and fusion of unligated DNA ends, a process we term breakage-replication/fusion (B-R/F). We propose that B-R/F provides a unifying explanation for complex rearrangement patterns including chromothripsis and chromoanasynthesis and enables rapid DNA amplification after chromosome fragmentation.

---

Rearrangement of DNA sequence is a fundamental process driving speciation^1^, adaptation^2^, and tumorigenesis^3^. Rearrangements can shuffle the order of DNA sequences without changing their copy number; such alterations can arise from chromosome breakage and fusion, or “cut-and-paste” processes^3^. Rearrangements can also be accompanied with changes in the copy number^4–6^ of repeats^7^ or large chromosome segments^8,9^. Copy-number alterations are associated with congenital disorders^10,11^ and are a prominent feature of cancer genomes^3,4,12^. The underlying mechanisms of DNA rearrangements with copy-number alterations, especially DNA duplication and amplification, remain poorly understood.

Early investigations of DNA amplification suggested that duplicated or amplified DNA^13,14^ is produced by two classes of processes. The first involves uneven distribution of replicated DNA between daughter cells. Such processes produce segmental DNA gain in one daughter cell and reciprocal loss in the other^15^, which can be regarded as a “cut-and-paste” outcome. One well-known example of this process is the breakage-fusion-bridge (BFB) cycles^16,17^, a mechanism proposed for intra-chromosomal DNA amplification^18^. The second process that can lead to DNA gain involves DNA re-replication. Re-replication was initially suggested to occur through repeated firing of replication origins within one cell cycle^14^ as occurs in the amplification of *Drosophila* amplicons in follicle cells^19^. However, deregulation of replication origin firing is thought to be uncommon in eukaryotic cells^20,21^. The generation of net DNA gain is now commonly attributed to a form of conservative DNA synthesis known as break-induced replication (BIR)^22,23^ or Microhomology-Mediated BIR (MMBIR)^24^. MMBIR was first proposed as the mechanism that generates interspersed segmental gains (known as chromoanasynthesis) in congenital diseases^11^. It was later suggested to produce “copy-and-paste” rearrangements in cancer genomes^3^.

Determining the etiology of DNA duplications is a central to dissecting cancer genome complexity. The key difference between “cut-and-paste” and “copy-and-paste” processes is whether the primary event is DNA breakage (‘cut’) or DNA synthesis (‘copy’). In the context of DNA rearrangements, this difference leads to different interpretations of the origin of rearrangement junctions: Do they arise from the ligation of ancestral DNA break ends (“cut-and-paste”) or from the invasion of a break end into an intact donor template (“copy-and-paste”)? For complex genomes with many rearrangement junctions, the ambiguous provenance of individual breakpoints or junctions is compounded: Does the complexity of a genome indicate one insult of massive DNA damage, multiple rounds of DNA breakage, repeated strand invasions from a single break end, or some combination thereof?

We previously demonstrated that the distribution of broken DNA fragments from either micronuclei^15^ or chromosome bridges^25^ into a pair of daughter cells produce single-copy DNA gains and losses within one generation. By contrast, clones expanded from cells with broken chromatids from either micronuclei^26^ or bridges^25,27^ frequently acquire segmental DNA duplications or amplifications. The contrasting short- and long-term copy-number outcomes provide an opportunity to determine how the duplications arise during the downstream evolution of broken chromosomes—whether from further chromosome breakage (cut-and-paste) or involving aberrant DNA synthesis (copy-and-paste).

Here, we present a detailed analysis of segmental copy-number gains arising spontaneously (i.e., not under selection) in clones expanded from single cells that inherited a broken bridge chromosome. The copy-number gains occurred almost exclusively on chromosome 4 that was engineered by CRISPR/Cas9 to form a dicentric chromosome bridge^25^. By resolving the structure of rearranged chromosomes in many different subclones, we show that nearly all segmental copy-number gains are generated from DNA fragments of a single ancestral chromatid (‘breakage’). In contrast to the conventional cut-and-paste model where the fragments are ligated before replication, we demonstrate that many ends of these fragments are duplicated by conventional replication (‘replication’) prior to ligation (‘fusion’). The replication-fusion of DNA ends produces both DNA gains and losses by uneven sister DNA segregation without the need for aberrant DNA synthesis in BIR or MMBIR. We term this process breakage-replication/fusion (B-R/F) to indicate both replication-fusion and fusion-replication of DNA ends after a single episode of DNA damage. A single round of B-R/F inherently produces three copy-number states on a single chromosome, therefore providing a simple explanation for three-state copy-number oscillations commonly seen in both chromothripsis^28^ or chromoanasynthsis^11^. The rearrangement outcomes of B-R/F include both long-range and foldback junctions between duplicated DNA that often contain chains of short insertions. We provide evidence that the insertions are derived from short DNA fragments originating near the ends of large segments. Finally, we identify footprints of B-R/F in cancer genomes and explain how B-R/F can explain numerous features of focal amplifications.

Together, the B-R/F process provides a unifying mechanistic explanation for seemingly contradictory features of “cut-and-paste” and “copy-and-paste” rearrangements and clarifies the ambiguity in the interpretation of complex rearrangement patterns in both cancer and congenital diseases. We propose that the replication of unrepaired DNA ends is an engine of rearrangement complexity and copy-number amplification during cancer genome evolution and other biological contexts.

## RESULTS

### Determining the segmental structure of complex copy-number gains from copy-number variation

We focused on one clone derived from a single cell that inherited a broken chromosome 4 (primary clone 1a^25^, hereafter clone **a**): This clone showed many subclonal segmental copy-number gains and long-range rearrangements. Such patterns are frequently seen in cancer genomes but their origins are obscure: They have been described as “complex, unclassified”^3^ or chromothripsis-like^29^. Single-cell copy-number profiling of this clone revealed variable segmental copy-number gains in different subclones (**Extended Data Figure 1**). To precisely determine the copy number and breakpoints of duplicated or amplified DNA segments, we derived 20 subclones from single cells and performed deep sequencing (10-30x) on 8 representative subclones (**a1**, **a2**, **a3**, **a4-1**, **a4-2**, **a4-3**, **a5**, **a6**). To precisely resolve the junction sequences between rearranged DNA segments, we also generated PacBio circular-consensus sequencing (∼10x) on the parental clone. With the newly generated data, we were able to determine the copy number (**Figure 1A**), breakpoints, and junctions of all rearranged segments in each subclone (**Supplementary Table 1**).

**Figure 1.**
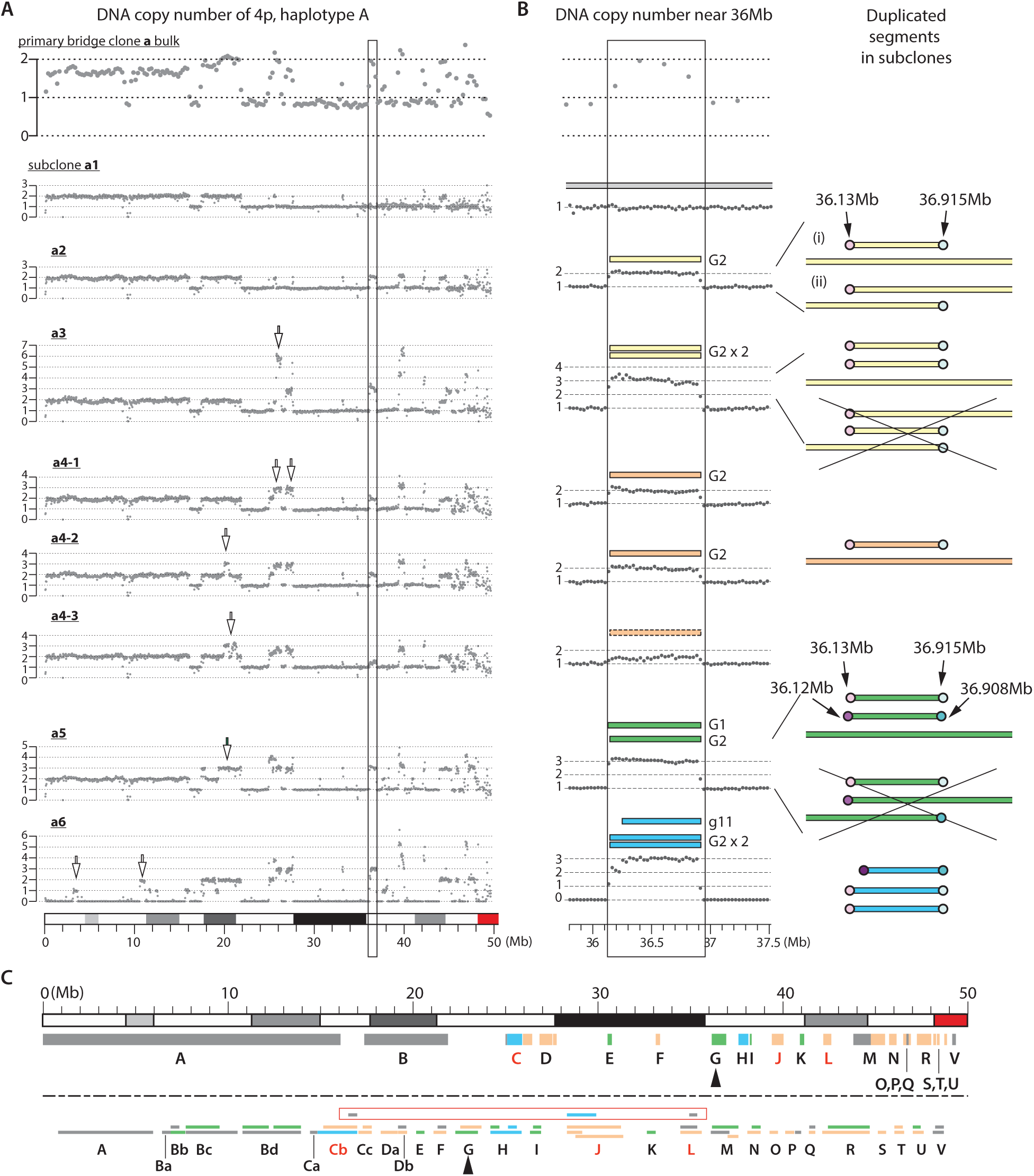
Segmental structure of duplicated DNA in a bridge clone determined from subclonal copy-number variation. **A.** Segmental copy-number gains in subclones derived from a single cell with a broken chromosome 4. Shown are the DNA copy number of a single haplotype (haplotype **A**) of the both the parental clone (primary bridge clone **a**, top, 250kb bins) and of different subclones (**a1**-**a6**, 25kb bins). Arrows point to duplications/amplifications that vary between different subclones. **B.** Copy number variation near 36.5Mb (outlined region in **A**) and the segments of duplicated DNA. The single-copy gain in subclone **a2** and **a4** can reflect either *cis* (i) or *trans* (ii) breakpoints. The *trans* configuration is ruled out by the presence of a two-copy gain of the same segment in subclone **a3** and **a6**. The triplication in subclone **a5** is decomposed into two sister segments **G1** and **G2**. The alternative configuration with two *trans* breakpoints is excluded based on the close proximity between breakpoints (light pink and purple on the left; light green and cyan on the right) implying an origin from staggered ssDNA ends of an ancestral dsDNA end. The triplication in subclone **a6** consists of two copies of **G2** plus a shorter segment **g11** that is a descendent of **G1** based on the shared breakpoint at 36.908Mb. **C.** *Top*: Locations and relative sizes of rearranged segments that produce the copy-number variation in subclone **a1-a6**. *Bottom*: Relative spacing between rearranged segments and breakpoints. See **Extended Data Figure 2** for more details.

We then determined the segmental structure of gained DNA by an integrative analysis of copy-number variation and breakpoints (**Methods**; **Supplementary Information, Sec. 3-5**). A central problem is to resolve the ambiguity of linkage between the breakpoints flanking duplicated segments. To illustrate this ambiguity as well as the approach to resolve it, we consider duplications near 36Mb on chr4 (**Figure 1B**). No duplication was present in subclone **a1**. A single-copy gain was detected in subclone **a2** and **a4** (the gain in **a4-3** was subclonal, likely reflecting loss during single-cell subcloning) and was flanked by unique breakpoints (left: 36.13Mb; right: 36.915Mb). The ambiguity is that the duplicated DNA could be contained either in a single segment (i) with two breakpoints in *cis*, or in two overlapping segments (ii) with breakpoints in *trans*. (See **Supplementary Information, Sec. 1** for definitions of breakpoints, segments, and junctions of rearranged DNA.) We were able to resolve this ambiguity by analyzing the copy-number variation between different subclones: The presence of a single segment with two *cis* breakpoints was established in subclone **a6** where the flanking sequences were completely deleted; the overlapping configuration was also excluded because it was inconsistent with the presence of two copy gains with the same breakpoints in subclone **a3** (**Supplementary Information, Sec. 2G**). By the same strategy, we determined that the subclonal copy-number variation at this locus was due to the variable copy number of three segments (**G1**, **G2**, and **g11**).

In total, we identified 86 rearranged segments with sizes above 10kb that accounted for the copy-number gains in all subclones; these segments were defined by 129 unique breakpoints, including 42 breakpoints shared by multiple segments (**Figure 1C** and **Extended Data Figure 2; Supplementary Table 2**). We next analyzed the evolutionary timing of the rearrangement breakpoints and assessed whether they were consistent with an origin from DNA breakage or aberrant DNA replication.

### Adjacent breakpoints and their mechanistic implications

The evolutionary timing of breakpoints can be inferred both from the distance between breakpoints and from the overlapping pattern of rearranged segments (**Supplementary Information, Sec. 2F** and **2G**). Here we focus on the inference of breakpoints that are not identical but much closer than expected by chance, which we refer to as **adjacent breakpoints**.

Consider a chromosome of size *L* with *K* breakpoints that are distributed randomly. It can be shown that the distance between breakpoints (*d*) approximately follows an exponential distribution: *P* (*d* ≥ *s*) = exp(-*s*/*D*), where *D = L/K* is the average spacing between breakpoints. In the bridge clone **a**, we identified 129 unique breakpoints on chr4p that is approximately 50Mb. Thus, the average spacing between neighbor breakpoints is 50Mb/129 ≈ 387kb. If these breakpoints are generated randomly, the probability of two breakpoints within 10kb is approximately 1 – exp(-10kb/387kb) ≈ 0.025, or 3.3 breakpoint pairs out of 129 breakpoints. By contrast, we observed 20 pairs of breakpoints within 10kb; this observation suggests many adjacent breakpoints are not generated by chance but by a concerted mechanism.

To relate the genomic feature of adjacent breakpoints to molecular mechanisms, we analyzed four types of adjacencies (**Figure 2**). The first type of adjacency is a small gap (i). The most plausible explanation of adjacent gapped breakpoints is that they originate from opposite DNA ends generated by a single double-strand break (DSB). This mechanism has been experimentally demonstrated in several contexts^30,31^. Consistent with this mechanism, we identified four groups of segments (“*cis* segments”) that were linked by gaps less than 1kb (**Extended Data Fig. 3A** and **B**). We also identified potential *cis* segments with larger gaps (10-30kb) that could be due to hyperresection of DSBs^32–34^ (**Extended Data Fig. 3C**).

**Figure 2.**
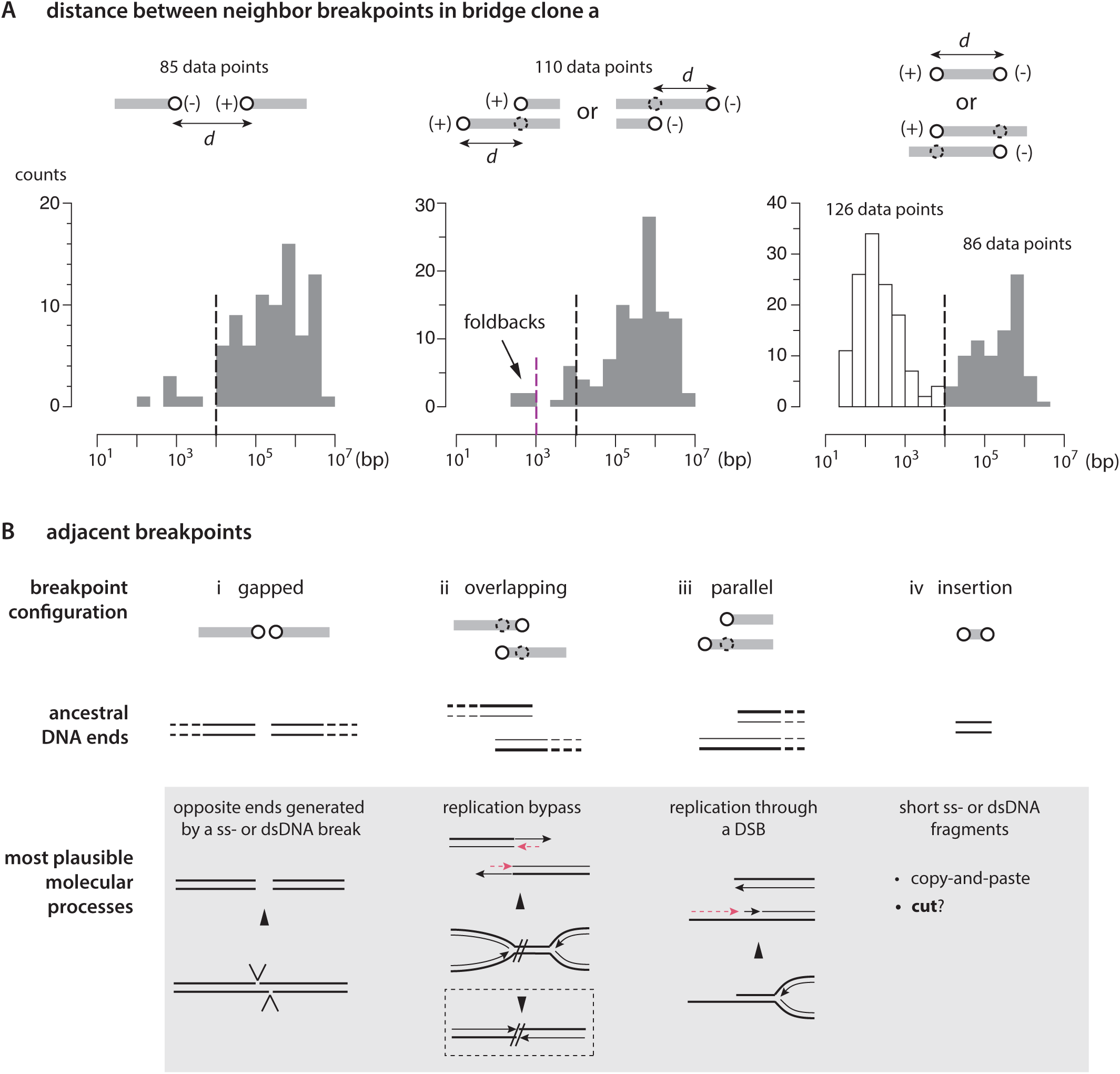
Adjacent rearrangement breakpoints and their mechanistic implications. **A.** Distance distribution of breakpoints in three configurations. Gapped: (-)(+); parallel:(+)(+) or (-)(-); insertion/overlapping: (+)(-). Adjacent gapped breakpoints are typically within 1kb (left); adjacent parallel breakpoints are typically within 10kb (middle); most short insertions are also within 1kb (right). **B.** *Top*: Four configurations of adjacent breakpoints. *Middle*: Inferred structures of ancestral DNA ends that become adjacent breakpoints. In (ii) and (iii), the two dsDNA molecules represent replicated DNA; the thick and thin lines represent the template and newly synthesized DNA strands. *Bottom*: Experimentally demonstrated mechanisms that can produce adjacent breakpoints. (i) Opposite DNA ends are generated by a double-strand break followed by end processing. (ii) Bidirectional replication fork bypass results in a small duplication (≲10kb) on one chromatid but no copy-number change to the sister chromatid (dashed box). (iii) Replication through a dsDNA end creates two dsDNA ends on sister DNA. In (ii) and (iii), red dashed lines indicate second-strand DNA synthesis. (iv) Short insertions are thought to arise from template-switching during break-induced replication but can also originate from short ss- or dsDNA fragments (“cut”) that are incorporated during end-joining repair.

The second type of adjacency is a small overlap (ii). Segments with small overlaps can be explained by the replication-bypass mechanism for collapsed replication forks in BRCA1 deficient cells^35,36^. This mechanism was originally proposed to explain small tandem duplications (∼10kb)^35^ but was recently shown to also produce non-tandem *trans* duplications^37^. We identified two pairs of segments with minor overlaps (**Extended Data Fig. 3D**; **J1a**/**J1b**: 23kb; **M1a**/**M1b**: 3kb) that are consistent with the replication-bypass mechanism. A potential mechanism for these segments is shown in **Extended Data Fig. 3E**.

The third type of adjacency is *parallel* adjacency (iii), which to our knowledge has not been described. Adjacent parallel breakpoints have a unique mechanistic implication: These breakpoints originate from different DNA molecules (in *trans*). If the ancestors of these breakpoints are generated by a single event as suggested by their proximity, then the simplest explanation is that their ancestors are staggered 5’ and 3’ ends of a dsDNA end. To create adjacent breakpoints, the staggered ssDNA ends would have to have been converted to dsDNA ends by replication before being ligated to other DNA ends. (By contrast, if the dsDNA end is instead ligated to another DNA end before replication, then the junction sequence and breakpoints will be duplicated, creating what we refer to as ‘flush’ breakpoints.) Therefore, adjacent parallel breakpoints are a specific signature of an ancestral DSB end that undergoes replication before ligation. In total, we identified 27 pairs of adjacent parallel breakpoints. Further evidence supporting their origin from unrepaired dsDNA ends will be presented in the next section.

The last type of adjacency (iv) reflects short DNA sequences that are often referred to as ‘insertions.’ In human genomic rearrangements, short insertions were first reported in amplified DNA in cancer genomes^38^ and subsequently recognized as a prevalent feature in both congenital disorders^11,39^ and cancer^28,29,40^. The two breakpoints of a short insertion are almost certainly generated simultaneously, but what mechanisms produce these insertions are unresolved^41^. Short insertions are commonly interpreted as the outcome of aberrant DNA synthesis in MMBIR-type processes^11^; however, this interpretation lacks definitive experimental support^41^. In total, we identified 126 short insertions (median:184bp, mean:545bp; **Table S2**). These insertions were clearly distinct from large segments based on the size distribution (**Figure 2A**, right).

In summary, we identified four classes of adjacent breakpoints associated with distinct molecular mechanisms. We next present evidence demonstrating that adjacent parallel breakpoints and short insertions are generated from replication of dsDNA ends.

### Adjacent parallel breakpoints are generated by replication of staggered DNA ends

We identified 27 pairs of adjacent parallel breakpoints (**Supplementary Table 2**). In 21 pairs, the ratio of breakpoint distance (min:223bp; max:17284bp) to segmental size was much smaller (<0.05) than expected if they were generated independently (**Figure 3A**). In the remaining six pairs, the breakpoint distance was in a similar range and likely had the same origin (e.g., DSB hyperresection), even though the ratio of breakpoint distance to segment size does not reach the 0.05 threshold because the segments were shorter.

**Figure 3.**
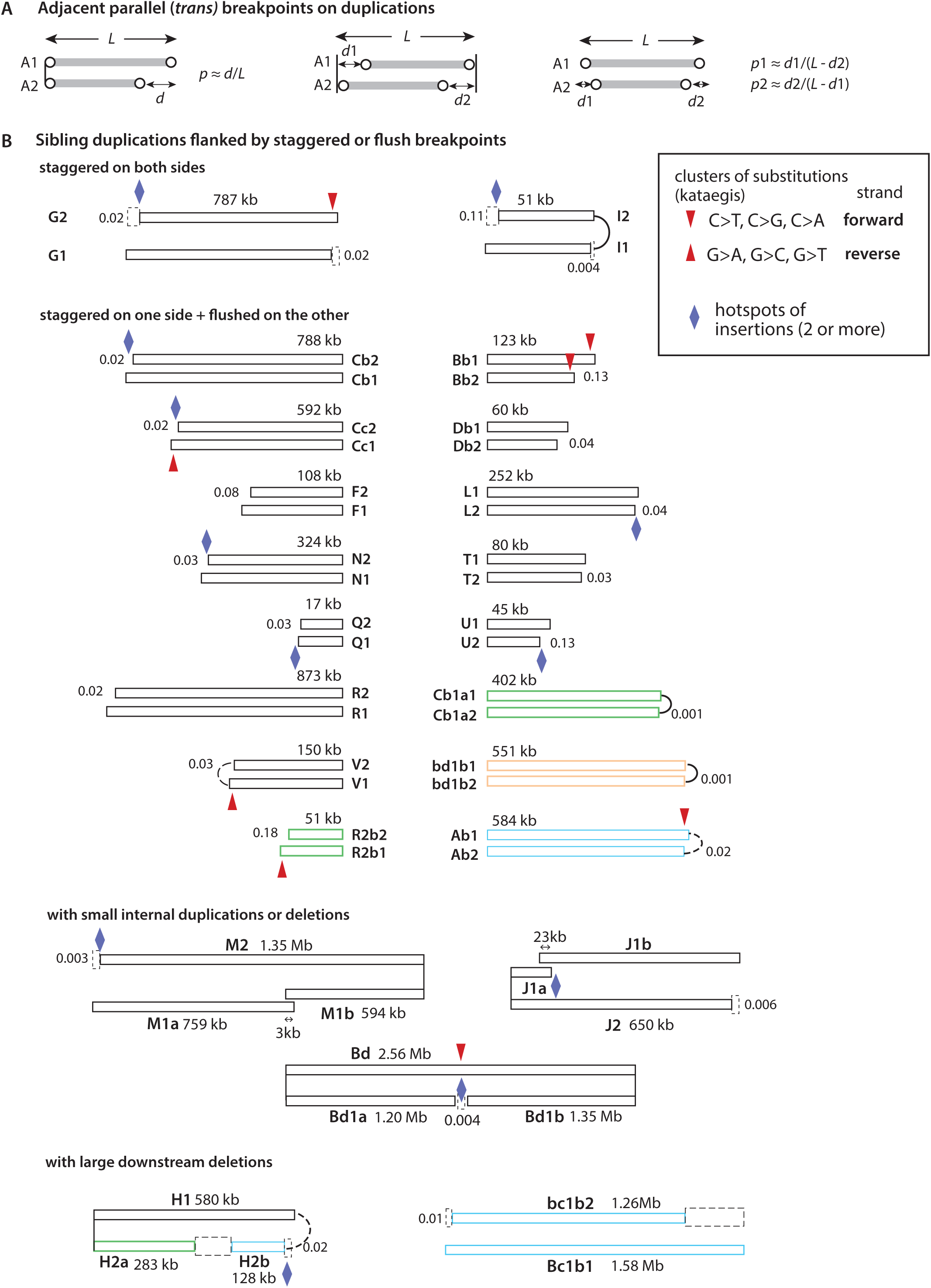
Sister duplications generated from an ancestral dsDNA fragment by replication. **A.** Assessment of the independence of *trans* breakpoints on duplicated segments based on breakpoint distance and segment size. **B.** Examples of duplications inferred to have been generated by replication of ancestral DNA fragments. Ancestral duplications are colored in black; secondary duplications that are private to an individual subclone are colored according the subclonal origin (orange for subclone **a4**, green for subclone **a5**, blue for subclone **a6**). Segment sizes are labelled but not shown true-to-scale. *Top*: Two pairs of duplications with staggered breakpoints on both sides. *Middle*: Sixteen pairs of duplications with flushed breakpoints on one side and staggered breakpoints on the other side. *Middle bottom*: Three sets of segments consisting of two *cis* segments paired with a sister segment. *Bottom*: Two pairs of duplications where one segment acquires large secondary deletions. The number near each pair of breakpoints is the ratio of breakpoint distance relative to the segment size. Smaller values indicate that breakpoints are less likely to have been generated independently. Arcs connecting adjacent breakpoints represent foldback junctions: solid arcs for direct foldbacks; dashed lines for foldbacks with insertions. Clustered substitutions (kataegis) reflecting deamination by the APOBEC family enzymes are highlighted with arrows: downward arrows indicate deamination of cytosines on the forward strand (Tp<u>C</u>>Tp<u>T</u>, Tp<u>G</u>, or Tp<u>A</u>); upward arrows indicate deamination of cytosines on the reverse strand (<u>G</u>pA><u>A</u>pA etc.). Note that when deamination occurs on the right end of a segment, it is always on the forward strand DNA (3’-end); when deamination occurs on the left end of a segment, it is always on the reverse strand DNA (also the 3’-end). The sister segments are drawn such that the top segment corresponds to the forward-strand DNA (5’ to 3’ from left to right) and the bottom segment corresponds to the reverse-strand DNA (3’ to 5’ from left to right). Purple diamonds indicate regions where multiple insertions are mapped. See Figure 5.

Four pairs of parallel breakpoints were present on segments (**Bb1**, **Cb1**, **Cc1, Cb1a**) that were inferred to have been generated by fragmentation (**Extended Data Fig. 3**). The presence of adjacent *cis* breakpoints provides strong evidence that the adjacent parallel breakpoints were generated from staggered 5’ and 3’ ends of an ancestral DSB by replication. For other adjacent parallel breakpoints for which the reciprocal *cis* breakpoints cannot be identified, their origin from staggered DSBs is established by the following observations.

First, almost all adjacent parallel breakpoints were present as pairs. (The only exception is explained in **Supplementary Information, pg. 26**.) The formation of paired breakpoints is expected if they arise from replication of two staggered ssDNA ends. By contrast, if these breakpoints are generated independently, either by strand invasion or from DNA breakage, the frequency of recurrence should follow a Poisson distribution and there should be more instances of ‘trio’ adjacency.

Second, the origin of adjacent parallel breakpoints from staggered DSB ends is supported by the presence of mutation clusters (kataegis) in the ‘overhang’ region between the breakpoints. All kataegis clusters showed the signature of deamination by APOBEC enzymes and nearly all were restricted to the overhang region. (The only exception on the shorter **Bb2** segment is discussed in **Supplementary Information, Sec. 8**). Moreover, the signature of deamination was in exact agreement with the prediction if the shorter breakpoint was derived from the resected 5’-end and the longer breakpoint was the 3’ end of the ancestral DSB: When the longer breakpoint is on the right side (q-ter), the overhang consists of DNA sequence on the forward strand and deamination of cytosines will lead to C>X substitutions (downward arrows); when the longer breakpoint is on the left side (p-ter), deamination of cytosines on the reverse strand will lead to G>X substitutions. The perfect agreement between the strand of deamination and the strand of DNA ends provides strong support for the origin of adjacent parallel breakpoints from staggered DNA ends.

Third, we observed strand coordination between breakpoints on opposite ends of ‘sister’ duplications. The simplest example is the **G1**/**G2** segments where the shorter end on one side is in *cis* with the longer end on the opposite side. This coordination is a direct prediction of the resection of 5’ but not 3’ ends of DSBs. Strand coordination was also observed for breakpoints flanking compound sister duplications consisting of multiple rearranged segments and between breakpoints and deamination (**Extended Data Figure 4**). For these compound segments, we inferred that the intermediate ‘flush’ breakpoints were ligated prior to replication, whereas the flanking adjacent breakpoints were generated by ligation after the replication of staggered DSB ends. Two exceptions to the long-range strand coordination of breakpoints were observed when one of two sister segments was interrupted with a partial duplication. This can be explained by sister chromatid exchange that occurs during the repair of overlapping DNA ends generated by replication bypass (**Extended Data Fig. 3E**).

Taken together, the above observations provide orthogonal evidence that strongly support the origin of adjacent parallel breakpoints from staggered ssDNA ends. Based on this feature, we further deduced that 50 out of 82 duplicated segments longer than 10kb were generated by the replication of ancestral dsDNA fragments, including segments that underwent secondary deletions. In addition, we also identified adjacent parallel breakpoints flanking duplicated segments in another chromosome 4 bridge clone (PC5a^25^), a post-crisis clone (X-29) from a study from a another group^27^, and in cancer genomes (see below). Therefore, adjacent parallel breakpoints are a specific signature of DNA ends that are replicated before ligation.

### Most breakpoints result from a single episode of DNA fragmentation

We inferred that all breakpoints that were not private to a single subclone (“ancestral breakpoints”) were generated concomitantly based on three constraints on the evolutionary timing of breakpoints: (1) Adjacent *cis* breakpoints arise simultaneously from a single dsDNA break; (2) adjacent parallel breakpoints arise simultaneously when a replication fork passes through a staggered dsDNA end; (3) breakpoints that are fused together at rearrangement junctions are also generated contemporaneously; when they originate from DNA ends, these DNA ends are also generated contemporaneously (i.e., within one or two cell cycles).

Based on the observation that the ancestral segments of the duplications (e.g., the parent segment of sister duplications) showed no overlap, we inferred that all the duplications were derived from fragments from a single chromatid. If the duplicated segments were derived independently, either from two sister chromatids (e.g., sister-chromatid chromothripsis^42^), from different copies of the same homologous chromosome, or from repeated invasions of the same donor template in MMBIR, we expect to see segments that have significant partial overlapping (i.e., ‘intersecting’ segments; **Supplementary Information, Sec. 2G**).

In summary, we infer that nearly all the rearranged segments detected in the subclones of primary clone **a,** including duplications, can be traced to ancestral DNA fragments derived from a single chromatid after a single episode of chromosome fragmentation.

### The Breakage-Replication/Fusion process generates three copy-number states in one cycle

The above observations suggest a breakage-replication/fusion (B-R/F) process that creates complex segmental duplications in three steps (**Figure 5**). The first event is extensive DNA damage that creates ssDNA breaks and staggered dsDNA ends. Such lesions are expected for DNA damage in micronuclei^43^ or chromosome bridges^44^, some of which can derive from nicks after incomplete base-excision repair^43^. Such nicks can become staggered DSBs or ssDNA gaps if they are processed by exonucleases (e.g., via TREX1^27^). Some dsDNA ends (e.g., near blunt ends) may be ligated in G1 by non-homologous end-joining (NHEJ). However, ssDNA ends, including those in staggered dsDNA ends, are often ineligible for NHEJ^45^ and persist into S/G2. Next, as the cell enters S phase, the unrepaired DNA ends and the associated DNA segments undergo semiconservative DNA replication. This step converts ssDNA ends into dsDNA ends^46^ and staggered DSBs into staggered dsDNA ends. The duplicated segments (sister segments) will have slightly different lengths both due to the difference between the ancestral 5’ and 3’ ends and due to the end-replication problem^47,48^. Finally, the sister dsDNA fragments undergo ligation in S/G2, generating rearranged chromatids with both deletions and duplications. Therefore, the replication and the fusion of DNA ends can occur in either order. We term this process the breakage-replication/fusion cycle.

Because a single cycle of B-R/F will generate rearranged chromosomes consisting of sister DNA segments from a single chromatid, the rearranged chromosomes will show three copy-number states reflecting deletion, retention of one sister segment, or retention of both sister segments. This mechanism thus provides a parsimonious explanation of chromothripsis with three copy-number states^28^ and chromoanasynthesis that commonly has three copy-number states^11^. A unique feature of B-R/F is the creation of adjacent parallel breakpoints by normal (S-phase) replication of unrepaired dsDNA ends. Moreover, the ligation of DNA ends both before or after replication will create rearrangement junctions with different sequence features: Junctions formed by DNA ligation in G1 (these give rise to flush breakpoints on duplicated segments) should reflect blunt end joining as they primarily occur by classical NHEJ. By contrast, junctions formed by ligation of sister DNA ends generated in S/G2 (these give rise to adjacent/staggered breakpoints) should display more microhomology as the sister ends are prone to DSB resection. Consistent with these predictions, we found that junctions between flush breakpoints were simple (direct joining or having one short insertion) and having little or no microhomology; by contrast, junctions between staggered breakpoints displayed more microhomology or contained more insertions (**Extended Data Figure 4**). Remarkably, the S/G2 junctions between sister DNA ends (adjacent parallel breakpoints) contained most of the inserted sequences (**Figure 2A**, right). We next discuss the unique features of the insertions that suggest they are also an intrinsic feature of B-R/F.

### Short insertions originate as ssDNA fragments mapped to regions adjacent to large segments

We identified 126 short insertions (**Supplementary Table 2**) that are shorter than 10kb (median:184bp, mean:545bp). Almost all the insertion junctions are directly resolved by long reads (Hi-Fi and uncorrected). As the mean read length is ∼9kb, the small size of these insertions (<1kb) is a feature of the underlying mechanism, not a technical limitation for detecting detect longer insertions.

We first analyzed the origin sites of the insertions in the reference. Three features of insertions at the mapped origins suggested a connection to DNA fragmentation. First, nearly all insertions (113/126) were adjacent to (<10kb) breakpoints inferred to be sites of ancestral DNA ends. Second, insertions were often adjacent to each other but did not have significant overlap (**Figure 5A**). 94 insertions were adjacent to another insertion (<100bp), and 76 formed 9 arrays of four or more insertions (**Figure 5B** and **Extended Data Fig. 5A**). In the most extraordinary example, we identified 18 insertions (median:171bp) tiling a 9.7kb region (46,166,808-46,176,544) with little gap (median 8bp; 2-42bp) or overlap (median 8bp; 4-15bp). This tiling pattern indicated that the insertions originate as non-overlapping fragments. Finally, six hotspots of insertions were located within 50bp from breakpoints inferred to be the 5’-end of ancestral staggered DSBs (**Figure 4B** and **5B**). The adjacency between these insertions and breakpoints derived from ssDNA ends indicates that the insertions originate as ssDNA fragments. Both the tiling pattern of insertions and their adjacency to ssDNA ends were also observed in another chromosome 4 bridge clone (**Extended Data Fig. 5B**) and a post-crisis clone from a previous study (X-29, **Supplementary Information, Sec. 7**), and in single cells with broken bridge chromosomes, including an example where we identified insertion hotspots on opposite ends of broken bridge chromosomes (**Extended Data Fig. 5C**). Together, these observations indicate that the short insertions originate as ssDNA fragments near staggered ssDNA ends; the tiling pattern of insertions strongly disfavors copy-and-paste mechanisms as repeated rounds of random strand invasion and copying from the same origin region would generate both gaps and overlaps (**Supplementary Information, Sec. 9**).

**Figure 4.**
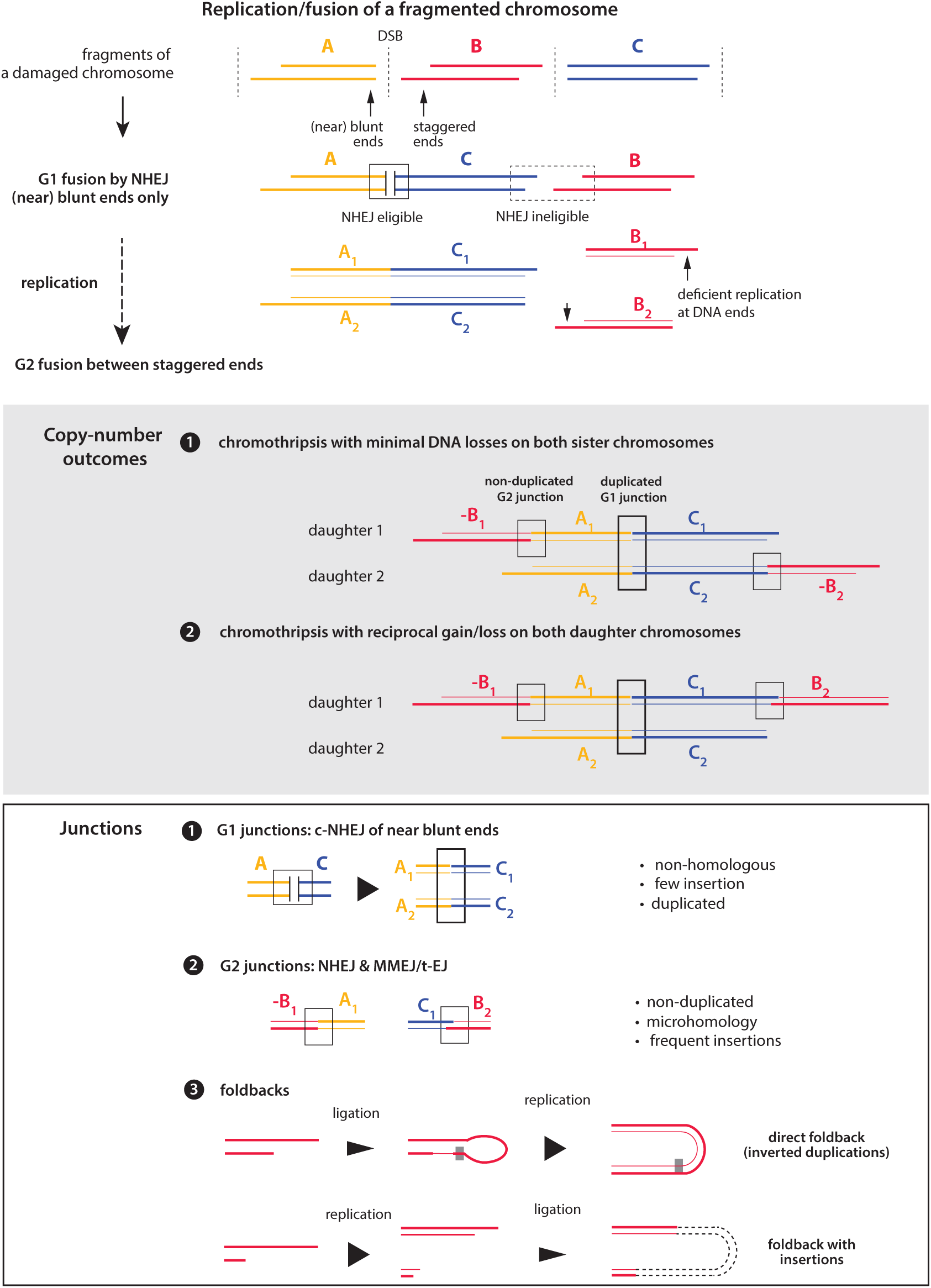
Rearrangement and copy-number outcomes of replication/fusion of a fragmented chromosome. (*Top*) Three dsDNA fragments (A, B, C) are generated from a fragmented chromatid. Near blunt DSB ends on segment A and segment C are joined by c-NHEJ in G1; segment B has staggered ends that remain unligated. DNA replication creates two sister DNA segments from the compound A:C segment and the B segment. The newly created dsDNA ends are then joined by NHEJ or MMEJ. (*Middle*) Copy-number outcomes: End-joining between sister DNA segments can create two different rearranged chromatids with minimal copy-number imbalance (1), or with reciprocal duplications and deletions (2). (*Bottom*) Junctions: The rearranged chromatids may contain three types of junctions: (1) junctions between near-blunt ends that are formed before replication; these junctions are duplicated in both sister segments (flush breakpoints); (2) junctions between unrelated sister DNA ends (staggered/adjacent breakpoints); (3) foldback junctions between sister DNA ends that can occur by ssDNA ligation (forming a hairpin) before replication or by end-joining between sister dsDNA ends after replication. See **Extended Data Figure S7** for examples.

Further evidence against a copy-and-paste origin of short insertions was provided by the analysis of insertions at their destination sites in the rearranged chromosomes. We assembled 17 junctions (c1-c17) with two or more tandem insertions (111 insertions total). Thirteen of these junctions (c1-c13) are shown in **Figure 5C**. All but one (c13) were directly resolved or validated by long reads (**Supplementary Table 1**). Given the *cis* relationship between insertions and adjacent segments at the origin sites, we could infer the original strands of the insertions (represented by arrows in **Figure 5B)** based on their adjacency to the 5’ or 3’ ends of ancestral DNA ends. For example, insertions originating from regions near the left end of the **G2** segment (5’-end on the ancestral DNA) must have originated from ssDNA fragments from the forward strand (**Figure 5A**). If the tandem insertions (**Figure 5C**) were generated by sequential episodes of conservative DNA synthesis as in MMBIR, all the sequences would be first copied to the 3’-end of one DNA strand and then undergo second-strand synthesis. Therefore, all the copied sequences would have to be on the same strand (the arrows should point to the same direction; **Supplementary Information, Sec. 9**). However, such “strand coordination” of insertions was clearly violated for insertions originating from the same hotspot in c2 (red), c3 (magenta and black), c5 (magenta), c6 (black), c7 (light blue), and c8 (light blue). Moreover, when considering all insertions whose original strands could be identified based on adjacent breakpoints, we found 36 instances where insertions next to each other show strand coordination but 43 instances when they show strand alternation (i.e., arrows pointing to opposite directions). These findings exclude the origin of insertion junctions from MMBIR. Instead, the insertions must have originated as ssDNA fragments, and the insertion junctions arise from the ligation of these fragments. The ligation could occur either by microhomology-mediated annealing of ssDNA fragments (which requires strand alternation) or dsDNA ligation after the ssDNA fragments are converted into short dsDNA fragments.

**Figure 5.**
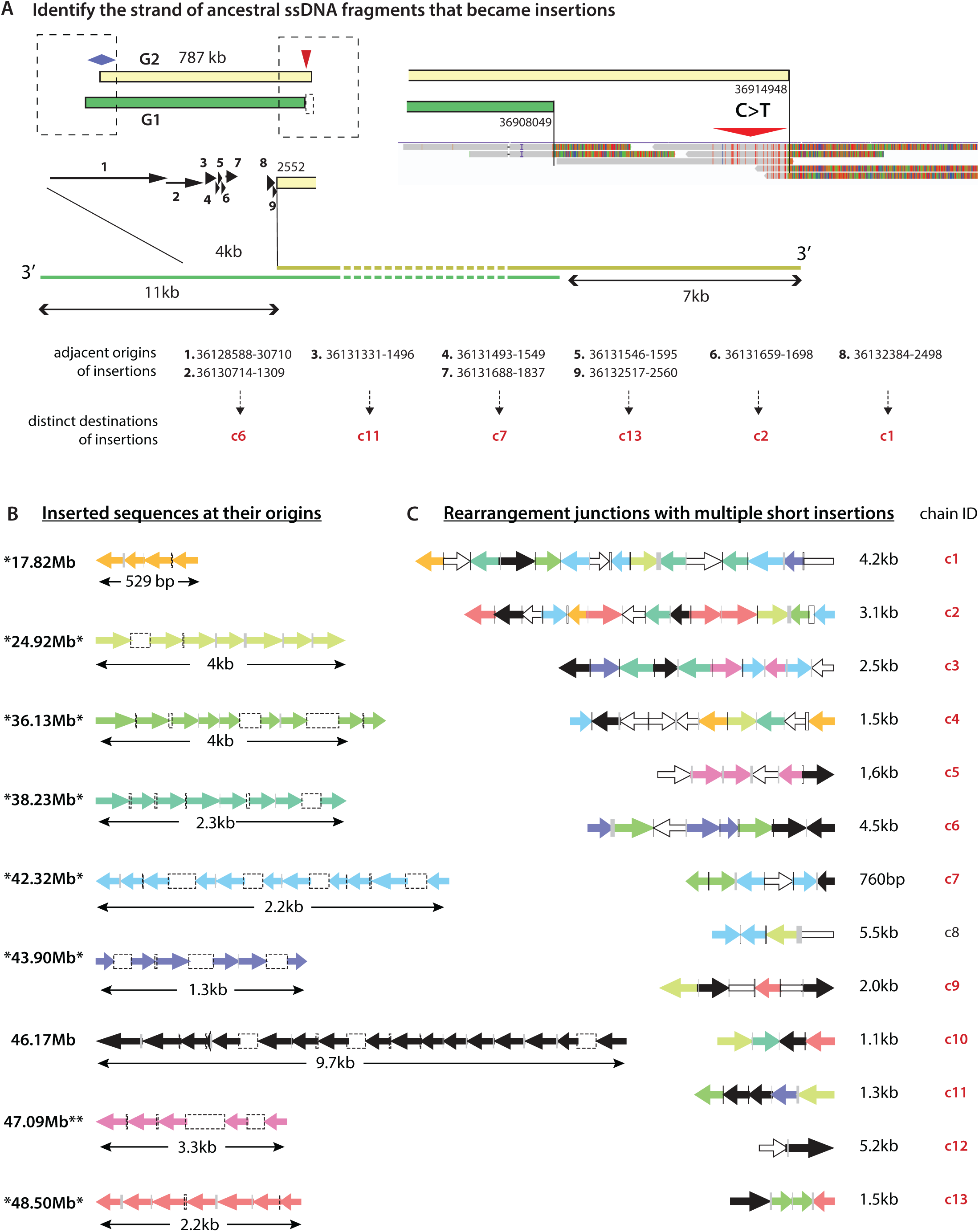
Origin and arrangement of short insertions. **A.** An example of nine short insertions mapped to a region adjacent to the left breakpoint of the **G2** segment. We infer these insertions to have originated as ssDNA fragments of forward strand DNA based on the signature of deamination (C>T) on the opposite (right) end of the **G2** segment. The original loci and destinations of the inserted sequences are shown below and the destination junctions are shown in panel **C**. The sizes of insertions and adjacency are shown true to scale. **B.** Tiling pattern of insertions at their original loci. Each locus contributes four or more insertions that are identified from different junctions (see panel **C** and **Supplementary Table 2**). Both the lengths of insertions (size of the arrows) and the adjacency between neighbor insertions (gaps or overlaps) are shown in logarithmic scale for better visualization (same in panel **C**). Except for the locus at 46.17Mb, all the other loci are adjacent to segmental breakpoints (denoted by *): Insertions at 17.82Mb are adjacent to a minus (-) breakpoint on the left; insertions at 47.09Mb have two adjacent parallel breakpoints on the right; the remaining hotspots are situated right next to the shorter breakpoint (inferred to be a 5’-end of the ancestral DNA) of two adjacent parallel breakpoints. The strands of insertions are inferred based on the polarity of the adjacent breakpoint and represented by arrows (5’->3’ of the original ssDNA sequence): leftward arrows correspond to reverse strand DNA; rightward arrows correspond to forward strand DNA. **C.** Arrangement of insertions at 13 destination junctions with two or more insertions (“chains” of insertions). Insertions are colored based on their origins on the left; open arrows represent insertions not from the hotspot origins shown on the left. The directionality of each arrow indicates the strand of the inserted sequence in the junction. Two open bars without arrowheads (at junctions **c1** and **c8**) represent insertions whose original strands could not be determined. If a chain is generated by sequential “copy-and-paste” of the ssDNA template, *all* the copied sequences have to be added to the same strand by DNA synthesis (5’->3’), i.e., the arrows will have to be colinear (as in **c11** and **c12**). Clear violation of strand coordination (arrows with both orientations) is seen in all the remaining chains.

In summary, our data suggest that short insertions originate from ssDNA fragments near ancestral DNA ends and are inserted between sister segments generated by replication of ancestral DNA fragments. (See **Supplementary Information, Sec. 9** for further discussion.) Therefore, both short insertions and large segmental duplications are intrinsic outcomes of B-R/F.

### B-R/F can lead to DNA re-replication and create unpaired DNA ends

If all fragments of the broken chromatid undergo complete replication, the B-R/F process will produce no net DNA gain or loss. However, the variable timing of DNA replication implies that unreplicated (template) DNA and replicated (daughter) DNA can co-exist during S phase: When replicated DNA is fused to unreplicated DNA with unfired origins, progression of the replisome from the unreplicated DNA into the replicated DNA can cause re-replication. This prediction explains the **l21** segment that creates a triplication (**Extended Data Figure 6**).

### Foldback junctions arise from fusions between sister DNA ends

The joining of two adjacent parallel breakpoints creates a foldback junction (**Figure 5**, bottom). Foldback junctions can result from replication through hairpins generated by self-annealing and ssDNA ligation of staggered ends as suggested in a recent study in yeast^49^. Foldback junctions also can result from the ligation of replicated sister ends (dsDNA ligation). Therefore, foldback junctions are a specific outcome of B-R/F that can occur both at the ends of broken bridge chromosomes and at unrepaired DNA ends after DNA breakage, including chromothripsis and incomplete retrotransposition^50^.

We identified six foldback junctions between duplicated segments of haplotype **A** (**Extended Data Figure 7**; for foldback junctions detected on the other haplotype of bridge clone **a** or in other bridge clones, refer to **Supplementary Table 1** and **Supplementary Information, Sec. 5**). Three were direct foldbacks and the foldback breakpoints are all within 1kb (**Extended Data Figure 7**, top). Three foldback junctions contained insertions (**Extended Data Figure 7**, bottom); in these junctions, the breakpoints were further apart (4.3kb, 10.7kb, and 10.9kb). These observations suggest that foldbacks can arise either by replication of a hairpin end, which cannot accept insertions, or by ligation of replicated sister DNA ends, which can incorporate insertions by a similar repair mechanism that creates long-range junctions with insertions.

For foldback junctions at chromosome ends, including those generated by the BFB cycle, we expect to see both deletion of DNA on the telomeric side of the foldback breakpoints and long-range palindromic structure in the rearranged DNA. By contrast, foldback junctions between sister DNA fragments are not subject to these constraints. Based on the long-range structure of the rearranged chromosome, we inferred that at least three foldback junctions [**I1(-)/I2(-)**, **Cb1a1(-)/Cb1a2(-)**, and **V1(+)/V2(+)**] were formed between sister DNA ends instead of sister chromosome ends (**Extended Data Figs. 6, 7**, and **Supplementary Information, pg. 33**). Therefore, foldback junctions can arise directly from chromosome fragmentation. This provides a parsimonious explanation for the co-ocurrence of foldback junctions and long-range rearrangements in complex rearrangements.

### Footprints of breakage-replication/fusion in cancer genomes

Above, we described three genomic signatures of breakage-replication/fusion. The first is adjacent parallel breakpoints (within ∼10kb) derived from the replication of unrepaired dsDNA ends. The second is junctions with short insertions (<1kb) mapped to regions near the breakpoints of large segments. Finally, for rearranged DNA sequences derived from fragments of a single chromatid, including both large segments and short insertions, the rearranged segments should have little or no partial overlap. We next sought to identify these features in cancer genomes.

We studied the breast cancer cell line HCC1954, one of the first cancer genomes to be analyzed by whole-genome sequencing^51^. The HCC1954 genome displays extraordinary complexity of amplified DNA from chr5, chr8, chr17, and chr21, some of which was suggested to reflect breakage-fusion-bridge cycles^38^. It is technically impossible to determine the complete structure of amplified DNA that consists of dozens or hundreds of highly similar sequences. However, the preservation of breakpoints on many copies of duplicated DNA enabled us to identify features of B-R/F (**Supplementary Table 3**).

First, we identified 121 pairs of adjacent parallel breakpoints (breakpoint distance <25kb) including 59 pairs that form foldback junctions (both direct and with insertions; median: 1244bp; mean: 2506 bp) and 62 pairs that form long-range rearrangements with different partners (median: 4722bp; mean:6744bp). The *trans* relationship of breakpoints was verified either using long reads (117/121) or by the presence of copy number transitions at both breakpoints (4/121). An example is shown in **Figure 6A** of the region of amplification on chr5q between 180 and 181Mb. The amplified DNA is flanked by two breakpoints at the centromeric (left) boundary (179.91 and 179.98Mb) and two adjacent breakpoints (181.24Mb) at the telomeric (right) boundary, the latter forming junctions with different partners (**Figure 6B**). Notably, 50 out of 62 non-foldback adjacent parallel breakpoints were identified in amplified regions from chr5, chr8, and chr21. The prevalence of adjacent parallel breakpoints in these regions with many inter- and intra-chromosomal rearrangements (similar to the example in **Figure 6A**) indicates a significant role of B-R/F in the amplification.

**Figure 6.**
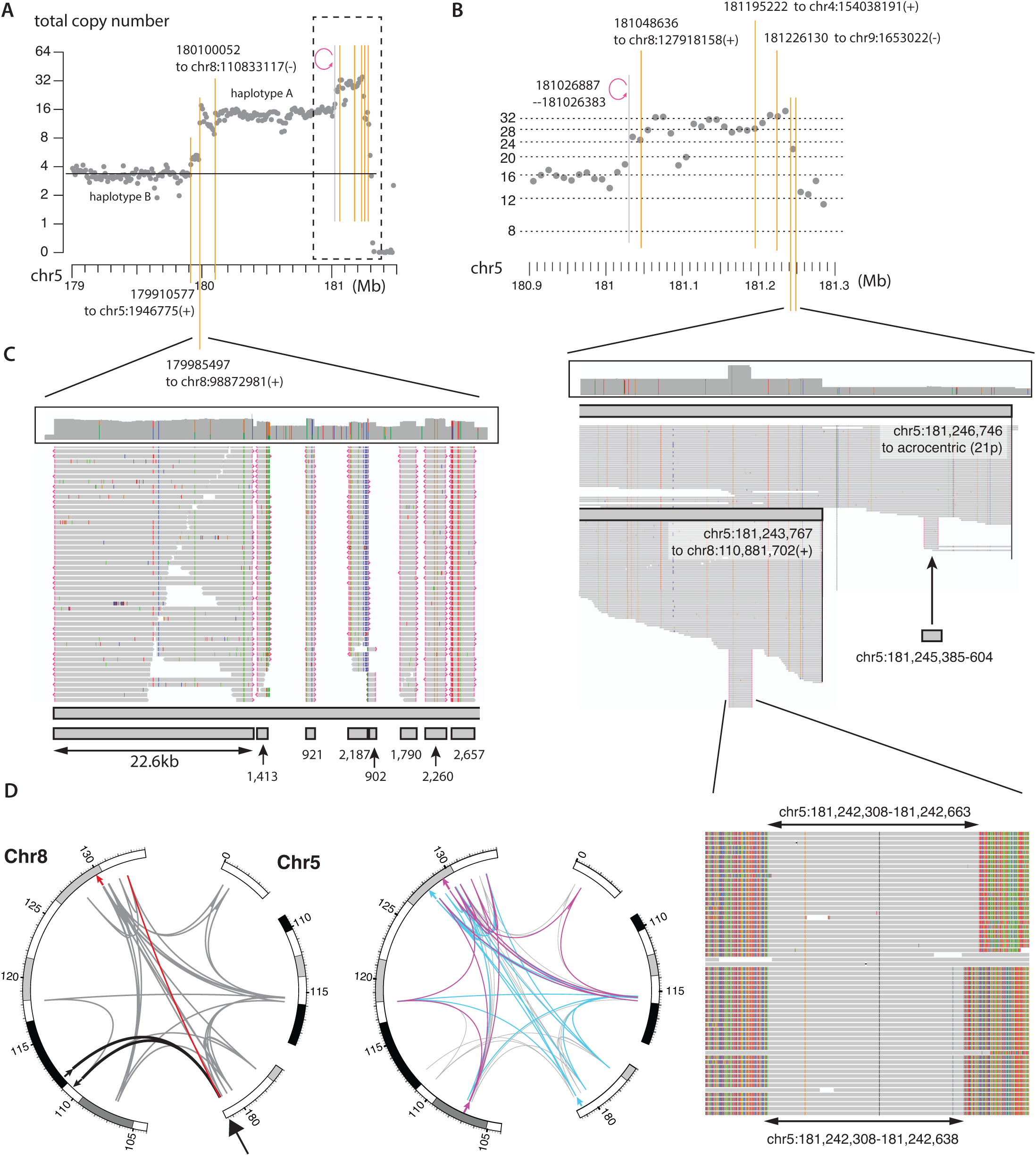
Footprints of breakage-replication/fusion in the HCC1954 genome. **A.** Total DNA copy number (log2 transformed, 10kb bins) in an amplified region (179-181.3Mb) of 5q. Amplified DNA is exclusively from haplotype A. **B.** *Top*: Five breakpoints with inter-chromosomal junctions (vertical orange lines) and a foldback junction (magenta circular arrow) at 181.03Mb between 181 and 181.3Mb. *Middle*: Zoomed view of long reads showing two adjacent parallel breakpoints (181,246,746 and 181,243,767, vertical lines) that are translocated to different chromosomes. *Bottom*: Short insertions (indicated by long reads with soft clipping on both sides) near the segmental breakpoints. Remarkably, there are two different insertions 181,242,308-638 and 181,242,308-663 with identical junctions on the left but different breakpoints/junctions on the right. The destination junctions of these two insertions are shown in panel **D**. **C.** Zoomed view of breakpoints/insertions near 180Mb of chr5. Shown are the total sequence coverage (histogram) and discordant reads in this region. The first row on the bottom represents a single segment with centromeric (left) breakpoint at 179985497; the second row displays non-overlapping insertions. The presence of a long insertion (22.6kb) is verified by Nanopore reads. **D.** Examples of complex insertion junctions containing chained insertions derived from multiple loci from chr5 and chr8. The left plot shows two junctions. The first one (thick solid lines) contains only one insertion, the longer insertion (181,242,308-663) shown in panel **B**. The second junction (gray lines) contains 47 insertions including the short insertion (181,242,308-638) shown in **B**. The junction between the breakpoint at 181,242,638 and the next insertion is shown in red; the final DNA breakpoint is highlighted with the red arrow. The right plot shows two more complex insertion chains (magenta and blue); the flanking breakpoints are highlighted with red and magenta arrows. Note the clustering of insertions from different chains at the same original locations near segmental breakpoints.

Next, we identified 351 insertions shorter than 10kb (median: 613bp; mean: 1552bp). A large number of insertions were derived from amplified regions from chr5 (95), chr8 (122), and chr21 (38). 53 insertions were adjacent (<1kb) to but not having a significant overlap with another insertion: One example of seven insertions whose origins were mapped to chr1:151.74Mb is shown in **Extended Data Fig. 8A**. Another example of non-overlapping insertions near a breakpoint of a large segment in shown in **Figure 6C**. Significant partial overlap was seen only between four pairs of insertions. Remarkably, we identified two pairs of insertions with ‘staggered’ breakpoints, one of which is shown on the bottom of **Figure 6B**: The longer sequence (356bp) was inserted between breakpoints at 111,694,761 and 110,510,787 (black arrows in **Figure 6D**, left); the short sequence (331bp) was one of 47 insertions (gray arcs) between 111,694,761 and 129,889,115 (red arrow). The most plausible explanation for these “sister insertions” is that they were derived from short, complementary ssDNA fragments at an unrepaired DNA end that underwent replication-fusion. Two more examples of junctions with multiple insertions (magenta and blue) are shown in **Figure 6D** (right), highlighting the proximal origins of insertions as shown in **Figure 6C**.

Consistent with the prior finding that replication bypass can generate 10kb duplications between overlapping breakpoints, we identified 50 pairs of adjacent overlapping breakpoints within 25kb. The distance between overlapping breakpoints (median: 6427 bp; mean: 7668 bp) was much longer than the size of insertions. Although we also identified 20 insertions above 10kb, the fraction of long insertions (20/370) is significantly lower than the fraction of large overlaps (17/50), highlighting different underlying mechanisms for insertions (fragmentation/ligation) and adjacent overlapping breakpoints (replication bypass).

Finally, we next assessed the frequency of partial overlapping between amplified DNA segments from chr5 and chr8. As the cancer cells were derived from a single clone with the amplification, we cannot determine the breakpoints of each segment based on subclonal copy-number variation. However, we can determine the segments of amplified DNA based on their *cis* Hi-C contact with non-amplified DNA in the rearranged chromosomes. For chr5, amplified DNA originated from three regions: 0-5Mb, 109-117Mb, and 179-182Mb (**Extended Data Fig. 9A**). Zooming into the largest region (109-117Mb, **Extended Data Fig. 9B**), we identified 11 different segments based on Hi-C contacts between chr5 and chr4 (both homologs). Only two segments displayed partial overlap (i.e., intersecting); the remaining segments were either inclusive or non-overlapping, similar to the pattern seen in the bridge clone (**Figure 1C**). A similar pattern was observed for amplified regions from chr8 (**Extended Data Fig. 9C** and **9D**) and chr21. The rarity of partial overlap between amplified DNA segments, together with the prevalence of adjacent parallel breakpoints, strongly supports an origin of amplified DNA from B-R/F.

A single B-R/F cycle is insufficient to generate high-level DNA amplification. So does the amplification require other processes such as BFB cycles or double-minute amplification? An intriguing feature is the concentration of breakpoints in the region of amplification as shown in **Figure 6A**. The clustering of breakpoints is also identified for foldback junctions on chr8p (**Extended Data Fig. 8B**), chr20q, chr12p, and chr17q (**Extended Data Fig. 8B-E**). In each case, there were three or more foldback junctions concentrated in regions less than 1Mb, with the most extreme case being three foldback junctions within 10kb on chr17. Nested foldbacks were previously attributed to BFB cycles and the location of each foldback junction was taken as the site of presumed chromosome breakage. According to this interpretation, the different foldbacks are generated independently over many generations; however, it is difficult to explain why the independently generated breakpoints are in close proximity. Positive selection may be invoked to explain amplified *ERBB2*, but the same argument does not apply to other regions without bona fide oncogene. We propose that nested foldbacks, and more generally, nested parallel breakpoints (e.g., near 55.35Mb and 55.42Mb on chr20q as shown in **Extended Data Fig. 8C**) are more plausibly and parsimoniously explained by multiple rounds of B-R/F fueled by the creation of new unligated DNA ends by replication (**Extended Data Fig. 10** and **Supplementary Information, Sec. 10**).

In summary, we identified signatures of B-R/F in a cancer genome including: (1) duplications with adjacent parallel breakpoints; (2) short insertions both derived from and inserted between segmental breakpoints; and (3) a largely non-intersecting pattern of duplicated DNA. We also identified nested foldbacks or parallel breakpoints in regions of DNA amplification that suggest the accumulation of high-level DNA amplification by multiple cycles of B-R/F. Therefore, sequential B-R/F cycles after one episode of massive DNA damage can generate a variety of complex rearrangement patterns with DNA copy-number gains or high-level DNA amplifications.

## DISCUSSION

The profound impact of gene copy-number variation on cellular and organismal phenotypes has been known since early studies by Alfred Sturtevant^52^. However, what processes produce somatic copy-number alterations, especially segmental duplications and amplifications, remain poorly understood. From an in-depth analysis of copy-number gains that arose spontaneously (i.e., without selection) during the evolution of a broken bridge chromosome, we have identified a general mechanism of DNA amplification that we term the breakage-replication/fusion (B-R/F) cycle. We suggest B-R/F explains several patterns of complex rearrangements with copy-number gain or amplification.

### B-R/F, BFB cycles, and chromothripsis

The central feature of B-R/F is the duplication/amplification of ss- and dsDNA ends by normal (S-phase) DNA replication prior to repair (fusion). Duplication of DNA ends is a feature of the chromatid-type Breakage-Fusion-Bridge cycle^16^ where the broken end of a broken bridge chromosome is replicated and then undergoes sister-chromatid fusion. Here we show that the same replication/fusion process can occur to dsDNA ends generated by various forms of chromosome fragmentation, creating adjacent parallel breakpoints as a specific genomic signature. Duplication of dsDNA ends is expected for damaged chromosomes from micronuclei^43^ or bridges^44^, as these chromosomes will contain DNA ends with significant ssDNA overhangs that are not readily repaired by c-NHEJ in G1^45^. Moreover, many forms of unrepaired dsDNA breaks carried over from the previous cell cycle (e.g., due to deficient homologous recombination or replication stress) will also have a similar structure due to DSB end resection. Therefore, duplication of unrepaired dsDNA ends should be a frequent outcome of DNA repair defects or genome instability, generating adjacent parallel breakpoints in cancer genomes.

In a single B-R/F cycle, the fusion of replicated DNA segments inherently produces both deletions and duplications, providing a unifying explanation of chromothripsis^28^ or chromoanasynthesis^11^ with three copy-number states. In contrast to the copy-and-paste model of classical BFB cycles or chromothripsis, a distinct feature of B-R/F is that fusions can occur either before or after replication, creating both foldback junctions (fusions between sister DNA ends generated from the same ancestral dsDNA end) and long-range rearrangement junctions (fusions between DNA ends generated from unrelated ancestral DNA ends). We suggest that the generation of these rearrangement outcomes by a single B-R/F cycle (e.g., after a single episode of chromosome fragmentation in a micronucleus or bridge) provides the most parsimonious explanation for the co-occurrence of long-range and foldback junctions in complex DNA rearrangements, which were previously attributed to sequential episodes of chromothripsis and BFB cycles^42^. We speculate that survival after a single episode of chromosome fragmentation, although rare, should be much more frequent than survival after multiple sequential chromosome catastrophes. Most importantly, sequential chromothripsis or BFB events cannot explain adjacent parallel breakpoints that do not form foldbacks or duplications that do not exhibit partial overlap, two specific predictions of B-R/F that are observed in the cancer genome analyzed here.

### B-R/F, BIR, and short insertions

It was previously suggested that resected DNA ends may invade non-allelic loci based on sequence homology or microhomology and initiate DNA synthesis by BIR^22,23^ or MMBIR^24^, creating both large segmental gains and short insertions^3,11^ by a copy-and-paste process. This proposal assumes that both large segmental gains and short insertions represent putative gains due to extra DNA synthesis. It is important to note that there has been no definitive evidence supporting this presumption^41^: The identification of net DNA gain requires copy-number information of both sister chromatids or daughter cells that is unavailable from the bulk average copy number of either a single cancer or an experimentally generated progeny clone. Moreover, we conclude that MMBIR type processes cannot be the major mechanism of either large segmental duplications or short insertions based on several lines of reasoning.

First, the rate of DNA synthesis by BIR or MMBIR is too low to produce large duplications. A recent study of BIR in yeast suggested the rate of BIR synthesis to be 0.5kb per minute^53^. At this rate, the upper limit of BIR synthesis over 20 hours (the entire interphase duration for RPE-1 cells) is 600kb. Moreover, BIR is interrupted by secondary structures or gene transcription^53^, further limiting its processivity. Therefore, a single round of BIR cannot produce the megabase-scale duplications that are commonly seen in cancer genomes or congenital disorders (**Supplementary Information, Sec. 6**). In B-R/F, there is no size limit of the duplicated sequence as duplication is generated by conventional DNA synthesis that can be carried out by replisomes fired from multiple origins within the duplicated segment.

Second, (micro)homology-dependent strand invasion cannot explain the prevalence of adjacent breakpoints in both experimentally generated clones (**Figure 3** and **Extended Data Figure 3**) and in the cancer genome that is analyzed here (**Figure 6** and **Extended Data Figure 8**). It also cannot explain why junction microhomology is only observed at staggered breakpoints but not at flush breakpoints.

A final argument against BIR and copy-and-paste processes in general is that duplicated DNA generated by sequential, random copy-and-paste events would generate duplications with frequent partial overlap. This contradicts the observation that partial overlap is rare, either between large duplications (>10kb) or among short insertions (<10kb). Instead, we find that duplications either have near identical breakpoints (flush or staggered), consistent with an origin from a single ancestral dsDNA segment that is replicated, or have little or no overlap, consistent with an origin from non-overlapping dsDNA segments from a single fragmented chromosome.

The most extraordinary non-overlapping pattern is observed for short insertions at their original loci (**Figure 5**; **Extended Data Figure 5**; **Extended Data Fig. 8A**). The tiling pattern of insertions (i.e., little overlap or gaps) suggests an origin from non-overlapping short DNA fragments that are difficult to produce by a series of strand invasion, DNA synthesis, and template switching events from an intact donor template. Moreover, conservative DNA synthesis in MMBIR implies that sequences copied to the original DNA end should all maintain the same 5’-3’ orientation (strand coordination) in the rearranged DNA. By contrast, we demonstrate that the strand orientation of insertions (at their destination loci) is essentially random (**Figure 5B**). Both the lack of partial overlap between insertions at the origin sites and the lack of strand coordination between insertions at the destination sites strongly argue against MMBIR as the mechanism producing chains of short insertions. Based on the adjacency between these insertions to breakpoints inferred to be the 5’-ends of ancestral DSBs, we speculate that these insertions could originate from ssDNA fragments generated by fill-in synthesis mediated by CST (CTC1, STN1, TEN1) and its associated DNA polymerase alpha/primase^47,54^ (see **Supplementary Information**, **Sec. 9** for further discussion). However, a detailed molecular mechanism awaits future studies.

In summary, the B-R/F model can explain all the features of large segmental duplications and small insertions, including many that are inconsistent with MMBIR or copy-and-paste processes.

### B-R/F, replication timing, and DNA amplification

In addition to creating DNA duplications through uneven sister DNA segregation, B-R/F can further lead to DNA re-replication by conventional DNA synthesis without origin re-licensing/firing: When a replicated DNA segment is fused to an unreplicated DNA segment with late-firing origins, replisomes from the late-replicating segment can progress into and re-replicate the early replicating segment (**Extended Data Fig. 6**). Notably, this mechanism will preferentially re-replicate segments with early-replication timing. We can also envision a scenario where late-replicating, origin-poor segments are fused together to create large “fragile sites:” These segments would be at risk of never completing DNA replication and being lost during clonal expansion. Therefore, the B-R/F model can produce putative DNA losses and DNA gains that are directly related to replication dynamics. Finally, the dynamic interplay between DNA replication and DNA repair may also influence the copy-number outcomes after chromosome fragmentation. For example, delayed DNA replication may enable more complete DNA repair and reduce the frequency of duplications due to B-R/F.

Iterations of B-R/F can generate high-level DNA amplification. Iterative B-R/F cycles can occur if some DNA ends (both ss- and ds-DNA ends) are replicated but not ligated, and undergoing replication/fusion cycles in subsequent generations (**Extended Data Figure 10** and **Supplementary Information, Sec. 10**). We predict that such iterative B-R/F cycles generate clusters of parallel breakpoints, including nested foldbacks, in regions of DNA amplification as seen in cancer genomes (**Extended Data Figs 8B-E**).

Counterintuively, DNA amplification can also result from the replication/fusion of many copies of ssDNA templates generated over multiple rounds of incomplete replication (**Supplementary Information, Sec. 11**). If a chromosome is trapped in a micronucleus in one generation, partial replication can generate two newly synthesized DNA strands; after reincorporation into a normal nucleus, these DNA strands can be converted into two extra copies by normal DNA replication. When a chromosome is trapped in a micronucleus over many generations, many rounds of partial replication could produce an onion-skin (nested) structure of amplified DNA, which may resemble the structure of amplified chorion genes in *Drosophila* follicle cells^19,55^. After reincorporation into a normal nucleus, the amplified DNA can be replicated and ligated to produce complex duplications consisting of nested segments as revealed in the analyzed cancer genome (**Extended Data Figure 9**). Therefore, the deficient replication of chromosomes entrapped in micronuclei may paradoxically generate high-level focal amplification of early-replicating DNA by a re-replication mechanism^14^ that does not involve deregulation of replication origin firing^20,21^.

In summary, our analysis suggests that a single B-R/F cycle provides a unifying explanation for seemingly contradictory features of cut-and-paste and copy-and-paste rearrangements and iterative B-R/F cycles enables rapid DNA amplification.

## Acknowledgment

We thank Dr. James Haber for suggesting the connection between adjacent parallel breakpoints and resected double-strand DNA ends, and discussions on break-induced replication, Dr. Heng Li for suggestions about long-read alignment and for discussions on the HCC1954 genome, and Dr. Neil Umbreit for discussions in the early stage of this study. The whole-genome sequencing libraries of bridge subclones were generated by Dr. Jinyu Wang with help from Dr. Umbreit. Dr. Ricardo Pinto assisted in the assembly of complex junctions. The Hi-C sequencing data of HCC1954 cells were generated by Dr. Jinyu Wang and the Hi-C contact map was generated by Dr. Richard Tourdot. Funding for this study comes from the Claudia Adams Barr Program for Innovative Cancer Research and the Innovations Research Fund from Dana-Farber Cancer Institute (to C.-Z. Zhang), Alex’s Lemonade Stand Foundation for Childhood Cancer (C.-Z. Zhang), BreakThroughCancer (C.-Z. Zhang), NCI (K22CA216319 to C.-Z. Zhang, R01CA213404 to D.P.), the Lustgarten Foundation for Pancreatic Cancer Research (to D.P.). DP is supported by the Howard Hughes Medical Institute.

## Methods Summary

### Data Availability

The long-read sequencing data of the HCC1954 genome are downloaded from the NCBI Short Read Archive (PRJNA1086849:SRR28305162). The Hi-C and short read sequencing data of the HCC1954 and its matching germline reference HCC1954BL are available under PRJNA1079784. The long-read sequencing data of the Bridge Clone 1a are available as SRR13579109. Short read sequencing data (Illumina) of the bridge subclones are available as SRR14017897-SRR14017905. Short read sequencing data of Bridge Clone 5a are available as SRR14017896. All these data are part of BioProject PRJNA602546 that also include short read sequencing data of single bridge daughter cells from a previous study. Short read sequencing data of RPE-1 clones from Ref. 27 are available from the European Genome-Phenome Archive under EGAD00001001629.

### Sequencing data processing

All short read sequencing data were aligned to the GRCh38 reference. Alignment and post alignment processing were described previously^56^. Long-read data were aligned using minimap2 (“-ax map-pb” for the bridge clone; “-a -k19 -w19 -U50,500 -g10k -A1 -B4 -O6,26 -E2,1 -s40” for the HCC1954 data).

### DNA copy number calculation

Haplotype-specific and total DNA copy number for all the RPE-1 samples (**Figure 1**, **Extended Data Figs 1** and **5**) was calculated using the RPE-1 haplotype data and the workflow as described previously^56^. Single-cell copy-number data (**Extended Data Figs. 1 and 5**) were calculated previously^25^. Copy number and Hi-C contact density for the HCC1954 genome were calculated as described previously^56^ and the results were reported in a recent study^57^.

### Detection of rearrangement junctions

Rearrangement junctions are detected from both long reads and short reads using the same algorithm as described previously^25^. Briefly, long reads with multiple split alignments were represented as “discordant reads” at each split junction. Rearrangement junctions were identified by clustering of discordant reads (both short reads and long reads).

### Haplotype phasing of rearrangement breakpoints

For the primary RPE-1 bridge clones generated in Ref. 25 and Ref. 27 and the subclones of primary bridge clone 1a, the breakpoints were assigned to the parental haplotype based on haplotype-specific copy-number transitions. We manually reviewed all copy-number transitions without detectable rearrangement junctions to identify the breakpoints: Nearly all of these breakpoints form junctions with repetitive sequences (e.g., centromeric/telomeric/rDNA repeats). We aligned the junction sequences to the CHM13 reference to identify the most likely locations of these sequences (e.g., chr13p) without the exact coordinates. The breakpoints of each segment and its copy-number state in each subclone were listed in **Supplementary Table 1**.

For the HCC1954 genome, all the breakpoints are affecting only one haplotype in the regions of amplification as shown in **Figure 6**, **Extended Data Figs. 8** and **9**. To show fine-scale copy-number transitions, we used total read depth in the zoomed copy-number plots.

### Determination of the segmental boundaries of copy-number gains

The central problem in the determination of rearrangement segments is to identify the two boundaries (breakpoints) of each segment. For the subclones of primary bridge clone 1a, this was achieved based on the following strategies. First, we determined the copy number of each breakpoint based on the copy-number transition. For adjacent breakpoints for which the copy-number transition at each breakpoint cannot be determined directly, we used the copy number of the partner breakpoint to determine the copy-number transition. Second, if a region of copy-number gain or retention is flanked by deletions, the flanking breakpoints directly determine the boundaries of the segment. This was applied to segments in the **a6** subclone where the intact chr4p was deleted. Once two breakpoints were determined to be in *cis* in a segment, we assumed their relationship to be preserved in all subclones. Third, in regions with variable or multi-copy gains, we assessed the copy number of each breakpoint in different subclones to identify *cis* breakpoints that always have the same copy-number state. Finally, if two breakpoints were determined to be in *cis* and each had an adjacent parallel breakpoint, then the adjacent breakpoints were also in *cis*. This is a direct prediction of the mechanistic constraint that adjacent parallel breakpoints are generated by replication of an unrepaired dsDNA end. The segmental copy number and breakpoints of all segments in primary clone 1a were listed in **Supplementary Table 2**. For additional information about this analysis, including the copy number and structure of all rearranged segments in each subclone, see **Supplementary Information**. The segmental breakpoint data were used to generate **Figures 2** and **4**, and **Extended Data Figs. 2-4.**

### Detection of single-nucleotide substitutions

Detection of short variants (single-nucleotide substitutions and small insertions/deletions) was performed using HaplotypeCaller as described previously^25^. We manually reviewed clustered mutations near breakpoints that are reported in **Supplementary Table 2**. The clustered mutation data were used in **Figure 4** and **Extended Data Figs. 2** and **4**. Both copy-number calculation and short variant discovery were performed using only short read data.

### Assembly of complex junctions with multiple insertions

We assembled insertions at rearrangement junctions by two approaches. When long-read data were available (for primary bridge clone 1a and HCC1954), the insertions were identified directly from the split alignments. For short read data, complex junctions were assembled from split reads and discordant reads. The assembly consists of five steps. First, for each rearrangement breakpoint, soft-clipped reads near the breakpoint (within 10kb up- or downstream) and their pairmates were collected. Second, soft-clipped reads were grouped based on the positions of split alignments and assembled into junctions based on sequence overlap across the junction. The assembled sequences were referred to as junction contigs since they were assembled from split reads at each junction. Only junction contigs with two or more supporting reads were considered. Third, the junction contigs were concatenated based on overlap determined by alignment (Smith-Waterman). This was performed first for contigs linked by the same sequencing read (i.e., a sequencing read with three split alignments supports two junctions), then between contigs with overlapping sequences spanning at least one junction, and finally between contigs supported by different pairmates of the same sequencing fragment (i.e., a fragment consisting of two reads each with two or more split alignments). We referred to the concatenated sequences as contigs. The supporting reads for all the junctions within each contig were grouped together. Fourth, the (concatenated) contigs were extended using the non-split pairmates of the supporting reads. The extended contigs were then concatenated based on overlap if they were linked by the same sequencing fragment. Finally, the supporting reads were aligned to the extended contigs (Smith-Waterman) and both the extended contigs and the supporting reads were aligned to the GRCh38 reference using bwa. The alignments were plotted for manual review.

For primary bridge clone 1a, the complex junctions were first assembled from the short reads data and then independently validated by the long reads. As a large fraction of the insertions were mapped to a few hotspots (**Figure 5**), we re-aligned the long supporting reads of the complex junctions to these regions using minimap2 with different parameters “-k13 -H -n2 -m14 -s28 -w5 -g 35” and “minimap2 - k17 -H -n2 -m20 -s40 -w5 -g 35” to increase the alignment sensitivity for short insertions. The original locations of the insertions and the order of insertions in the rearranged DNA were both listed in **Supplementary Table 2**.

For the HCC1954 genome, we selected three examples of complex junctions with multiple insertions (**Supplementary Table 3**).

### Identification of adjacent breakpoints in the HCC1954 genome

We chose an operational threshold of 25kb for adjacent breakpoints in the HCC1954 genome. We identified adjacent parallel breakpoints, insertions, and adjacent overlapping breakpoints based on the breakpoints detected jointly from long reads and the PCR free Illumina data. We then manually reviewed these adjacent breakpoints to validate the *cis*/*trans* relationship between breakpoints using long reads.

## DNA duplications from replication and fusion a single broken chromosome

**Extended Data Figure S1.**
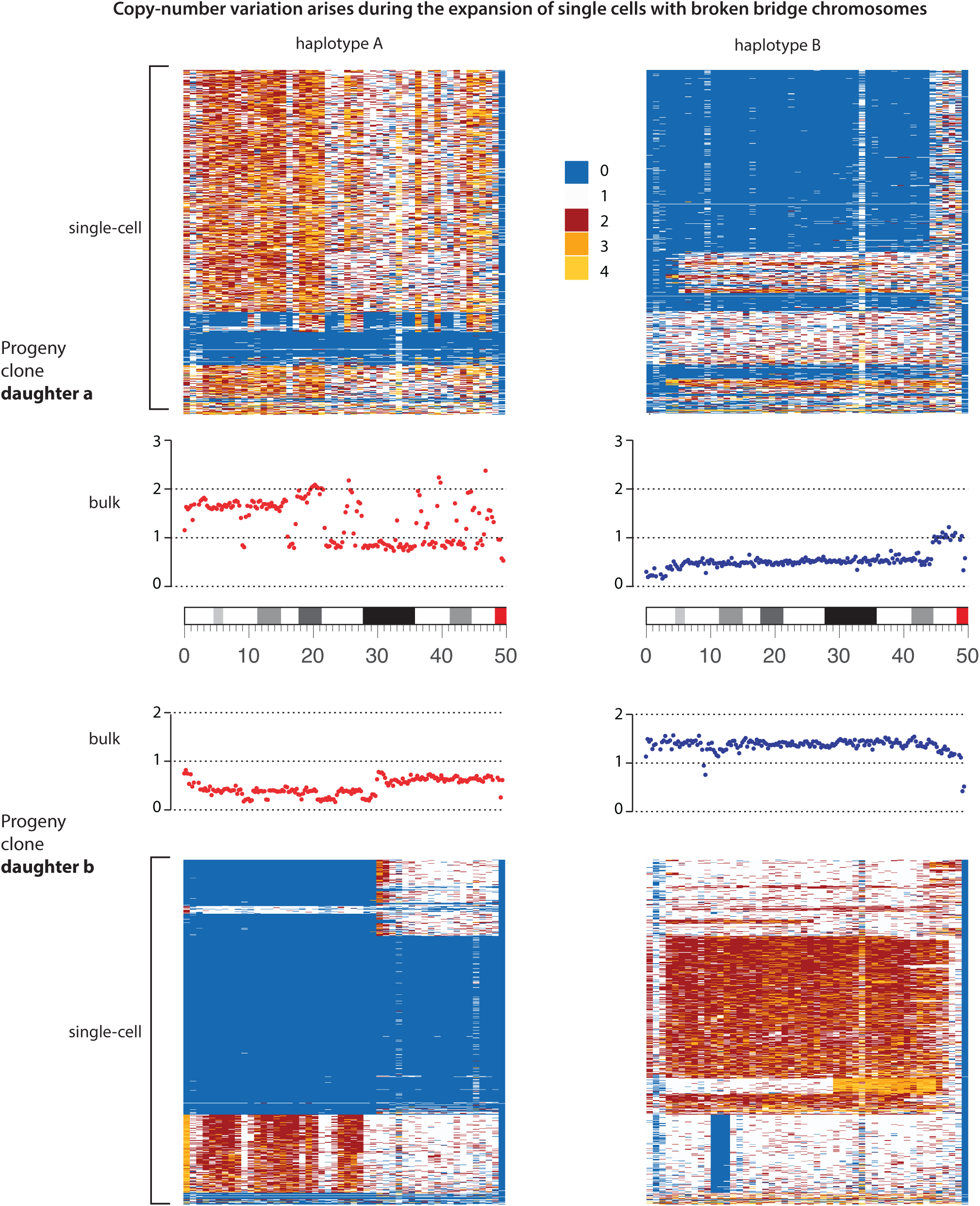
Copy-number variation of chr4p in the primary bridge clone **a** (top) and its sister clone **b** (bottom). Single-cell copy-number data are shown as heatmaps for haplotype **A** (left) and haplotype **B** (right). Bulk copy-number plots are shown in the middle rows: haplotype **A** in red and haplotype **B** in blue. The retention of most of chr4p in some cells in each clone implies that the ancestors of both clones (a daughter pair) must have inherited a near complete 4p. Therefore, the copy-number gains in either clone (e.g., haplotype **A** in clone **a** or haplotype **B** in clone **b**) must have arisen during the downstream evolution of the ancestral broken chromosome 4. In contrast, the reciprocal duplication and deletion of haplotype **A** between 0 and 30Mb in bridge clone **b** indicates a reciprocal distribution of two fragments generated by a secondary breakage.

**Extended Data Figure S2.**
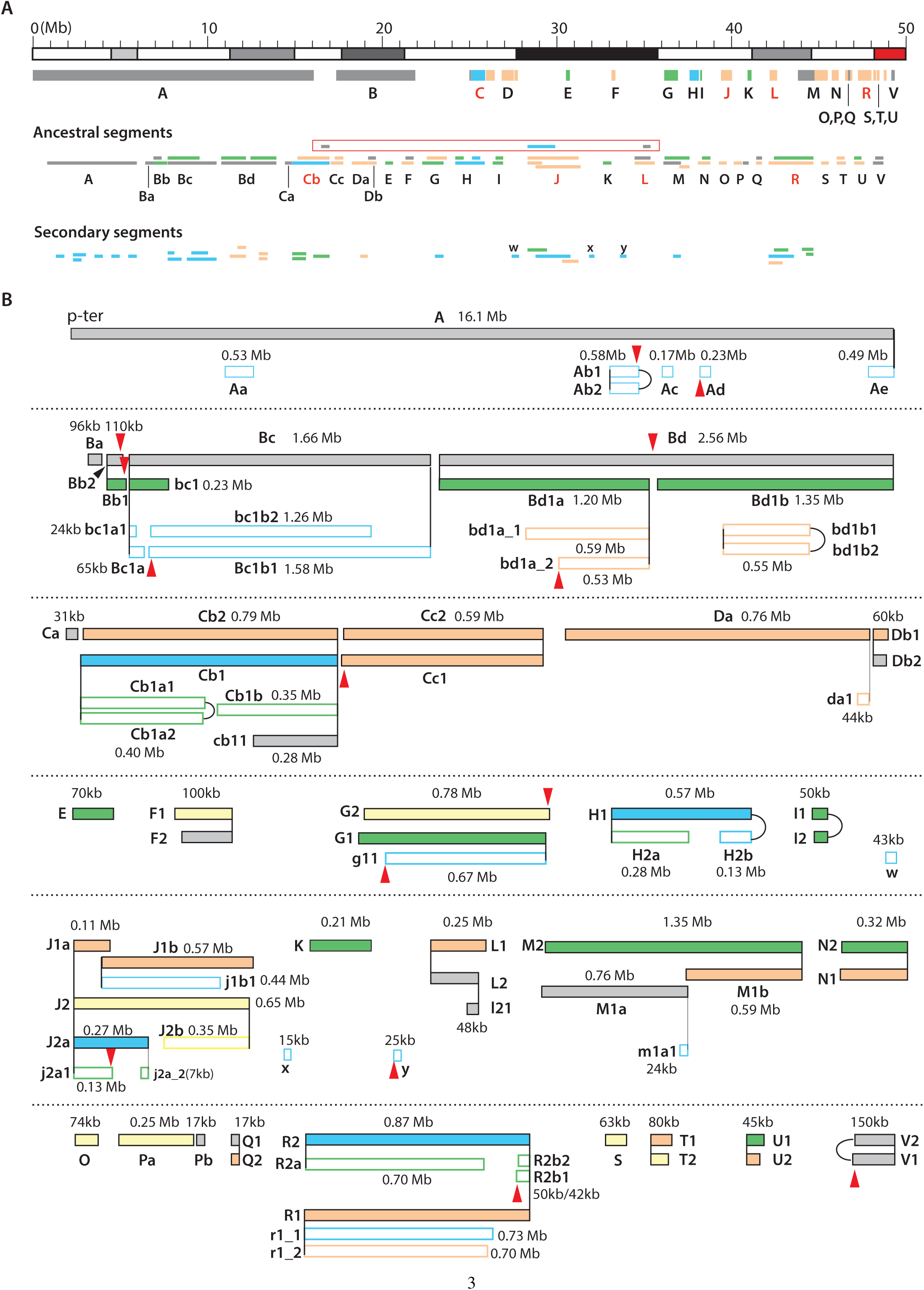
Rearranged segments of 4p identified in the subclones of bridge clone **a**. **A.** Locations, sizes, and relative positions of rearranged segments. *Top row*: segments are grouped (**A**-**V**, **w**,**x**,**y**) based on their original locations on 4p. *Middle and bottom rows*: Relative positions and sizes of ancestral and secondary segments. For better visualization, the spacing between breakpoints *x* is scaled by a nonlinear function *f* (*x*) = arctan(*x/*10, 000). The sizes of each segment is annotated in panel **B**. **B.** A detailed view of breakpoints of both ancestral (filled bars) and secondary (open bars) segments at each locus. Breakpoints shared by more than one segment are highlighted with vertical lines. Duplicated segments with identical breakpoints but present at multiple copies at different locations on the rearranged chromosome (exact duplications) are shown only once. A breakpoint is considered ancestral if (1) it is preserved in more than one subclone; or (2) it has an adjacent breakpoint and both breakpoints are identified in different subclones. A breakpoint is considered secondary otherwise. A segment is considered ancestral only when both boundaries correspond to ancestral breakpoints. Secondary segments have at least one boundary being a secondary breakpoint. *Color of segments*: Segments in both **A** and **B** are colored based on the subclone where it is preserved at the lowest copy-number state (gray:subclone **a1**; yellow:**a2** and **a3**; orange:**a4**; green:**a5**; blue:**a6**). *Naming of segments*: The names of segments reflect both their original positions (**A** to **V** from the telomeric end to the centromere) and ancestry. Segments inferred to have been derived from sister DNA fragments are indexed ‘1’ and ‘2’ with ‘1’ being the longer sibling. Segments inferred to have been derived from adjacent fragments from the ancestral chromosome are appended with ‘a’,‘b’, etc. (e.g., **Ba-Bd**). Segments that are truncated in comparison to the ancestor have lowercase letters, e.g., **bc** is a truncated copy of **Bc**. When more than one truncated segments are present, they are annotated using ‘_1’, ‘_2’, e.g., **r1_1**, **r1_2**. *Additional sequence features*: Clustered substitutions (≥3) with the signature of cytosine deamination are highlighted by arrowheads: downward arrows when deamination occurs to cytosines on the forward strand; upward arrows when deamination occurs to cytosines on the reverse strand. Junctions between rearranged segments are not shown except for foldbacks that join sister segments (e.g., **I1**/**I2**) or adjacent breakpoints (e.g., **H1**/**H2b**). *Note*: **H1** is only preserved in subclone **a6** but is inferred to be ancestral because its telomeric (left) breakpoint is shared with **H2a** (preserved in subclone **a5**) and its centromeric (right) breakpoint has an adjacent breakpoint on **H2b**. Based on the adjacent centromeric (right) breakpoints of **H1**/**H2b** and the shared telomeric breakpoint of **H1**/**H2a**, we infer that **H2a** and **H2b** are derived from an ancestor **H2** that is a sister of **H1**.

**Extended Data Figure S3.**
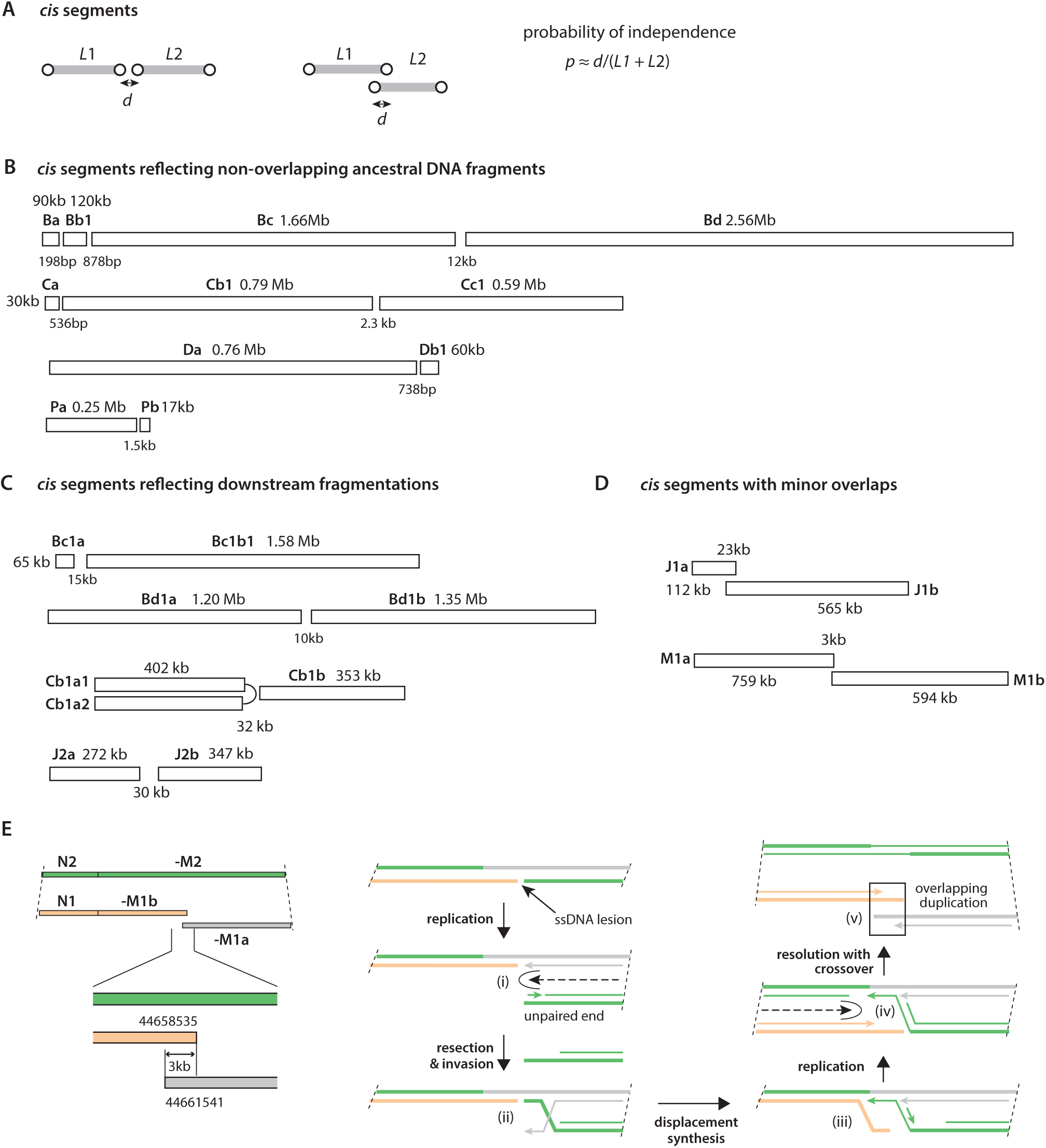
Adjacent breakpoints on *cis* segments generated by DNA fragmentation. **A.** Assessment of the probability that breakpoints on adjacent segments are generated independently. **B.** Non-overlapping segments with small gaps originate from fragments of an ancestral segment. Some of the segments are preserved in different subclones (see **Extended Data Figure S2**), but the adjacency between breakpoints suggests that they were generated simultaneously but later segregated into different subclones. **C.** Non-overlapping segments inferred to have arisen by secondary fragmentation of ancestral segments. **D.** Two instances of segments with small overlap likely generated by the replication bypass mechanism. **E.** *Left*: Structure of the **M1a**/**M1b** segments, their sister segment **M2**, and their neighbor segments **N1**/**N2** in the rearranged chromosomes. Colors reflect the subclonal origin of each segment as shown in **Extended Data Figure S2**. Note the pairing of the shorter breakpoint on **N2** with the shorter breakpoint on **-M2**). *Right*. A proposed model for the generation of **M1a**, **M1b** and **M2** by replication bypass. Thick lines represent template DNA strands; thin lines represent newly synthesized DNA; arrowheads denote the 3’-ends of newly synthesized DNA; DNA strands are colored by the same scheme as shown on the left. (i) A left-moving fork collapses (e.g., at a ssDNA gap) and creates an unpaired dsDNA end (green). (ii) The dsDNA end (green) is resected and invades the sister DNA (gray). (iii) The invading end initiates DNA synthesis and displaces the orange template, similar to break-induced replication (BIR). (iv) A right-moving fork encounters the BIR fork and collapses. (v) A resolution involving cleavage of the gray strand causes a sister-DNA exchange (crossover) that results in the opposite coordination between the DNA ends, i.e., the shorter end on the left is in *cis* with the shorter end on the right. There are three possible outcomes of the two break ends (orange and gray) with partial overlap. First, they could anneal with each other by homology; this would result in error-free repair with sister-DNA crossover. Second, if the 3’-ends (arrowheads) are extended (i.e., gap-filling synthesis), the dsDNA ends can join each other by c-NHEJ or MMEJ, creating a tandem duplication. This outcome was previously demonstrated in cells with BRCA1 deficiency. Finally, the two ends may be separately ligated to other DNA ends, as is seen here.

**Extended Data Figure S4.**
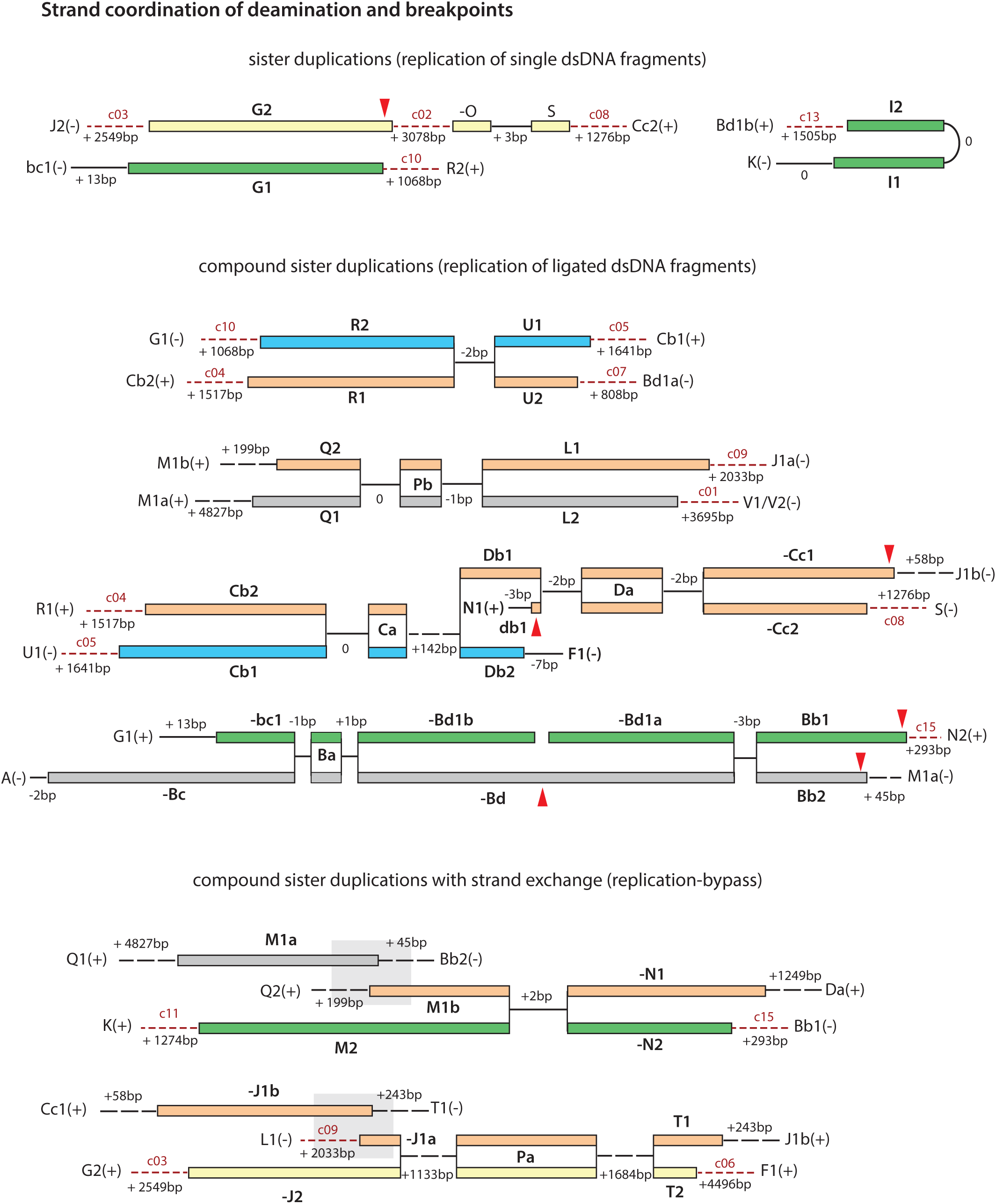
Long-range strand coordination of deamination and DNA ends in compound segments. Segments are colored by the subclonal origin (**Extended Data Figure S2**) and arranged as in Figure 3. Strand coordination is observed for two features. (1) Shorter breakpoints on one side (5’-ends of the ancestral DNA) is in *cis* with longer breakpoints on the opposite side (3’-ends of the ancestral DNA). (2) Deamination signature is consistent with the ancestral DNA strand of the overhang as inferred from the breakpoints. Note that the red arrows are oriented relative to the segment where deamination occurred. Therefore, if the segment is shown in a reverse orientation (e.g., **-Cc1** and **-Bd**), the signature of deamination needs to be reversed accordingly. There are two instances with apparent violations of strand coordination of breakpoints. (1) The shorter end of **M2**(+) is in *cis* with the shorter end of **N2(**+); (2) the shorter end of **J2(-)** is in cis with the shorter end of **T2(-)**. We attribute both instances to sister-DNA exchange during the replication-bypass repair inferred from the small duplication (gray boxes). We note that strand coordination only applies to breakpoints that originate from ancestral ssDNA ends, but not junctions that form between dsDNA ends generated by replication of ssDNA ends. For example, the left breakpoint of **G2(**+) (originally 5’- end of forward strand DNA) is ligated with the right breakpoint of **J2(-)** (originally 5’-end of reverse strand DNA). However, junctions between sister DNA ends show more frequent insertions and more extensive microhomology than junctions between flush breakpoints that are formed before replication. Junction complexity: Solid lines for direct joining; long dashed lines for the presence of one insertion; red short dashed lines for the presence of two or more insertions. ‘+’ basepairs indicate the length of total inserted DNA, ‘-’ base pairs indicate the length of microhomology. The complex junctions (c1-c15) are discussed in Figure 5.

**Extended Data Figure S5.**
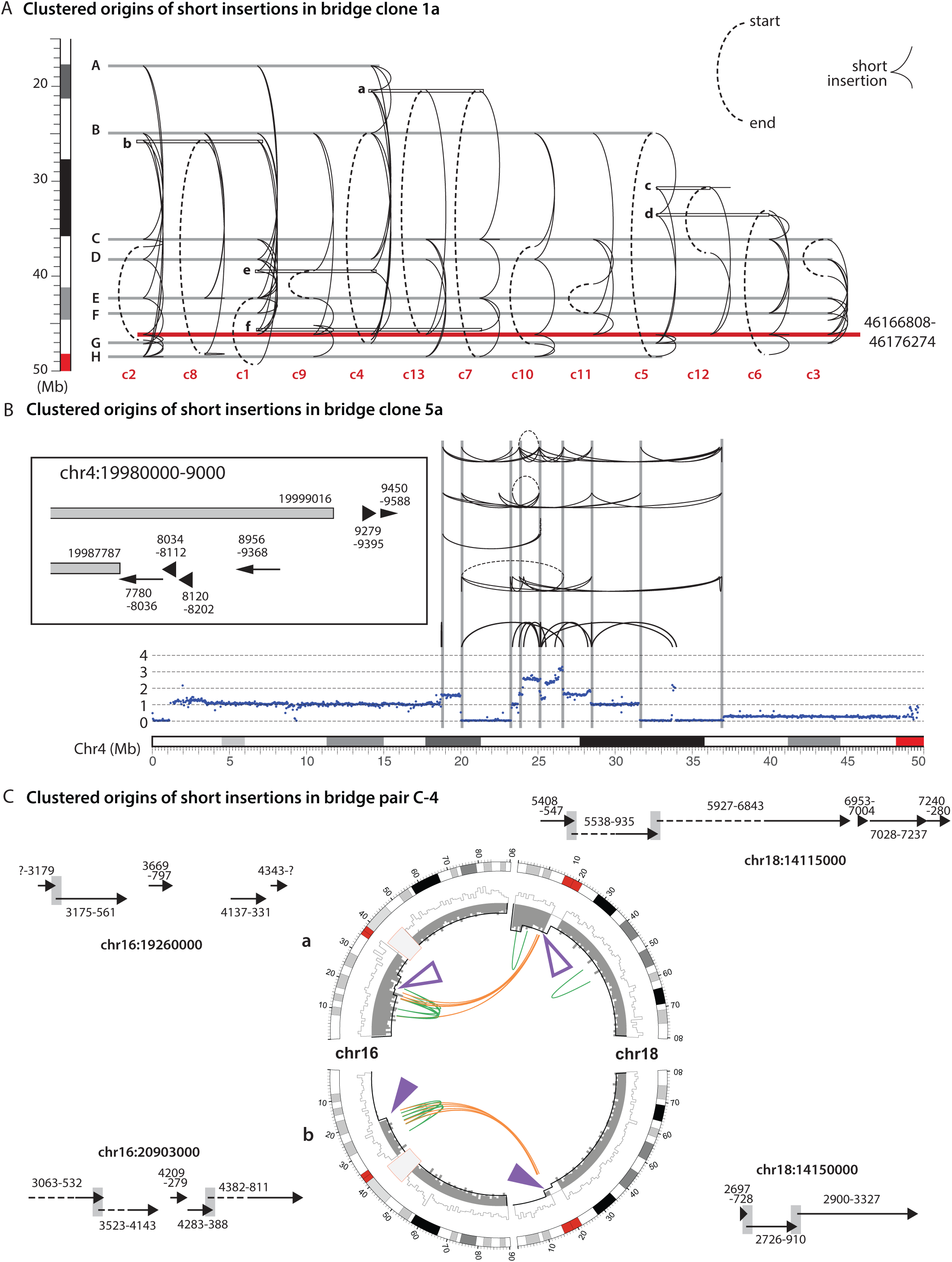
Further examples of short insertions mapped to regions near large segmental breakpoints. **A.** Footprints of the 13 chains of insertions as shown in Figure 5C. For each chain (c1-c13), the breakpoints of the flanking segments are shown as a dashed line, arcs represent insertions that are strung together between these breakpoints. Regions **A-H** correspond to the eight hotspots shown in Figure 5B; regions **a-f** each contribute two or three insertions. **B.** Five complex insertion junctions identified in primary bridge clone 5a from Umbreit et al. (2020). Chained insertions are shown similarly as in **A** except that the chromosome is drawn horizontally. The inset shows an example of multiple insertions mapped to regions adjacent to parallel breakpoints (at 19,999,016 and 19,987,787, 11,229bp difference). **C.** Two daughter cells each having inherited two broken chromosomes from a bridge (bridge pair C-4 from Umbreit et al. (2020)). In the CIRCOS plots, the DNA copy number of the broken chromatid is shown in gray (inner histograms) and the copy number of the intact homolog is shown in the outer histograms; intra- and inter-chromosomal rearrangement junctions are shown as green and orange arcs. The sites of chromosome breakage (purple arrows) are identified from the pattern of reciprocal DNA gain and loss (∼ 20Mb on chr16 and ∼ 14Mb on chr18). All four chromosome ends contribute multiple insertions as shown. Dashed lines indicate insertions whose continuity cannot be ascertained as their sizes exceed the size of sequence fragments. See **Supplementary Information** for additional examples.

**Extended Data Figure S6.**
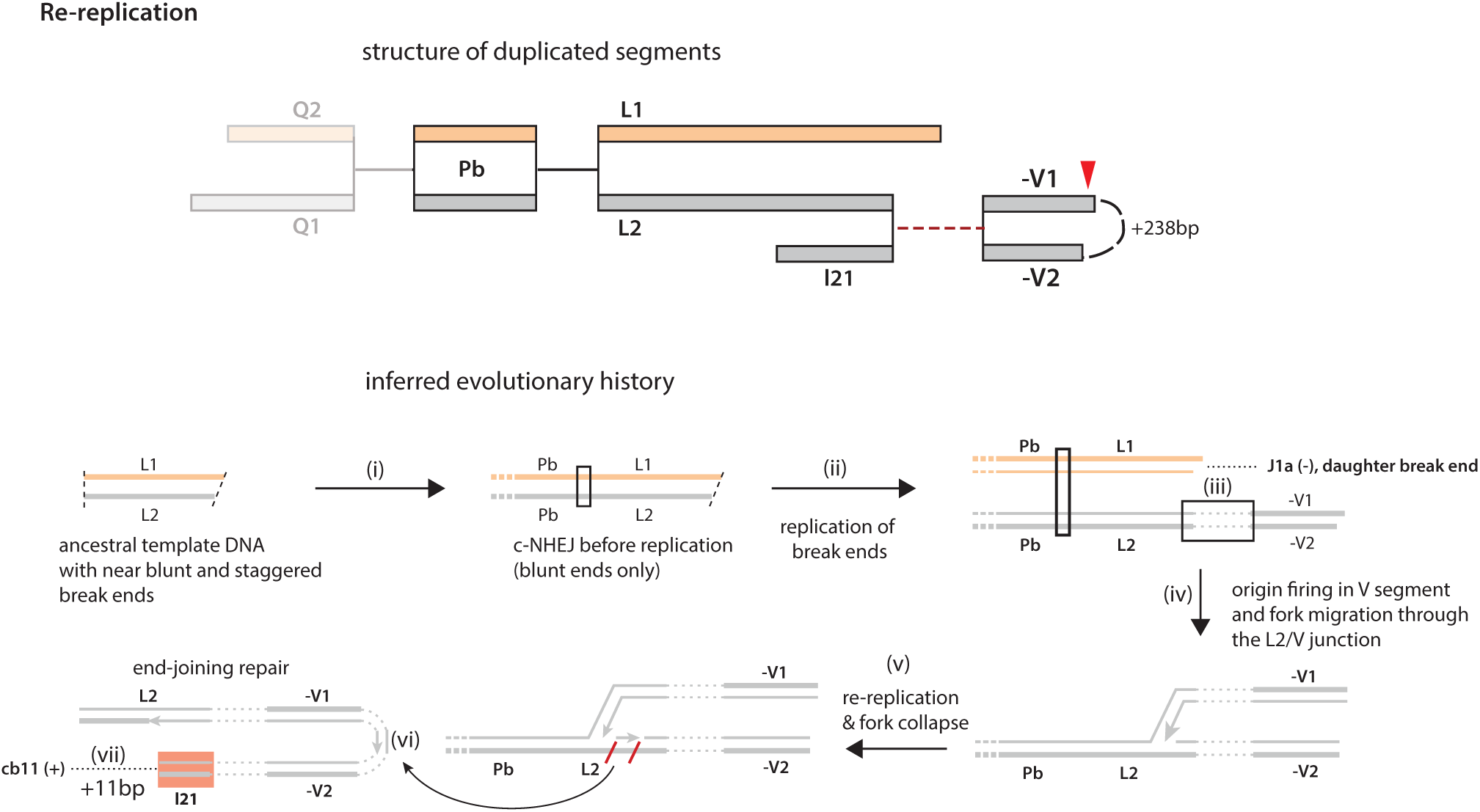
An example of DNA re-replication due to the progression of a replication fork through a junction between pre- and post-replicated DNA. In this example, a small duplication **l21** is inferred to be generated by the replication fork from the ancestor dsDNA fragment of **V1**/**V2** passing through a junction with a replicated sister DNA end on the **L2** segment. This inference is based on the following reasoning. First, the right breakpoint of **L2** has an adjacent breakpoint on segment **L1**, suggesting that both segments are generated by replication of an ancestral fragment (i and ii). The inference that the duplicated junction between **L2**/**l21** and **-V1**/**V2** results from a fusion between sister DNA ends is further supported by the complexity of the fusion junction (c1). Second, the **V1**/**V2** segments are sister segments generated by replication of an ancestor fragment. This inference is based on adjacent parallel breakpoints (49,172,365/49,176,671) on **V1**/**V2** and the presence of two G>A substitutions in the overhang region that reflect deamination of DNA overhang on the reverse strand. Third, the replication that produces **V1**/**V2** must have occurred in the same generation as the replication that produces **L1**/**L2**. This is because the **V1**/**V2** segments are located in the middle of the rearranged chromosome with sister duplications **Q1:Pb:L2** and **Q2:Pb:L1** on opposite sides (**<u>Supplementary Information</u>**). If **V1**/**V2** were generated in a later generation, the parallel breakpoints would have been a terminal DSB end on the rearranged chromosome. Therefore, the junction between **L2**/**l21** and **V1**/**V2** is formed between the replicated **L2** segment and an unreplicated **V** segment (iii) that is replicated later in the same cell cycle. Following the firing of replication origins within the **V** segment, bidirectional fork progression generates a pair of break ends (staggered breakpoints of **V1**/**V2**), duplicates the **L2:-V** junction, and further causes partial re-replication of **L2** (iv and v). Importantly, the replication fork passing into the **L2** segment will not be able to merge with an opposite fork since the **L2** is already replicated: This replication fork eventually collapses (vi), creating an unpaired break end on **l21** that is then ligated to an unpaired break end on the **cb11** segment. The presence of an insertion within the foldback junction between **V1**/**V2** that is mapped to chr4:42,275,638-873 right next to the breakpoint of **l21** at 42,275,875 supports that steps (iii-iv) occur with in a single cell cycle. A similar model may explain the triplicated **cb11** segments (**Supplementary Information**).

**Extended Data Figure S7.**
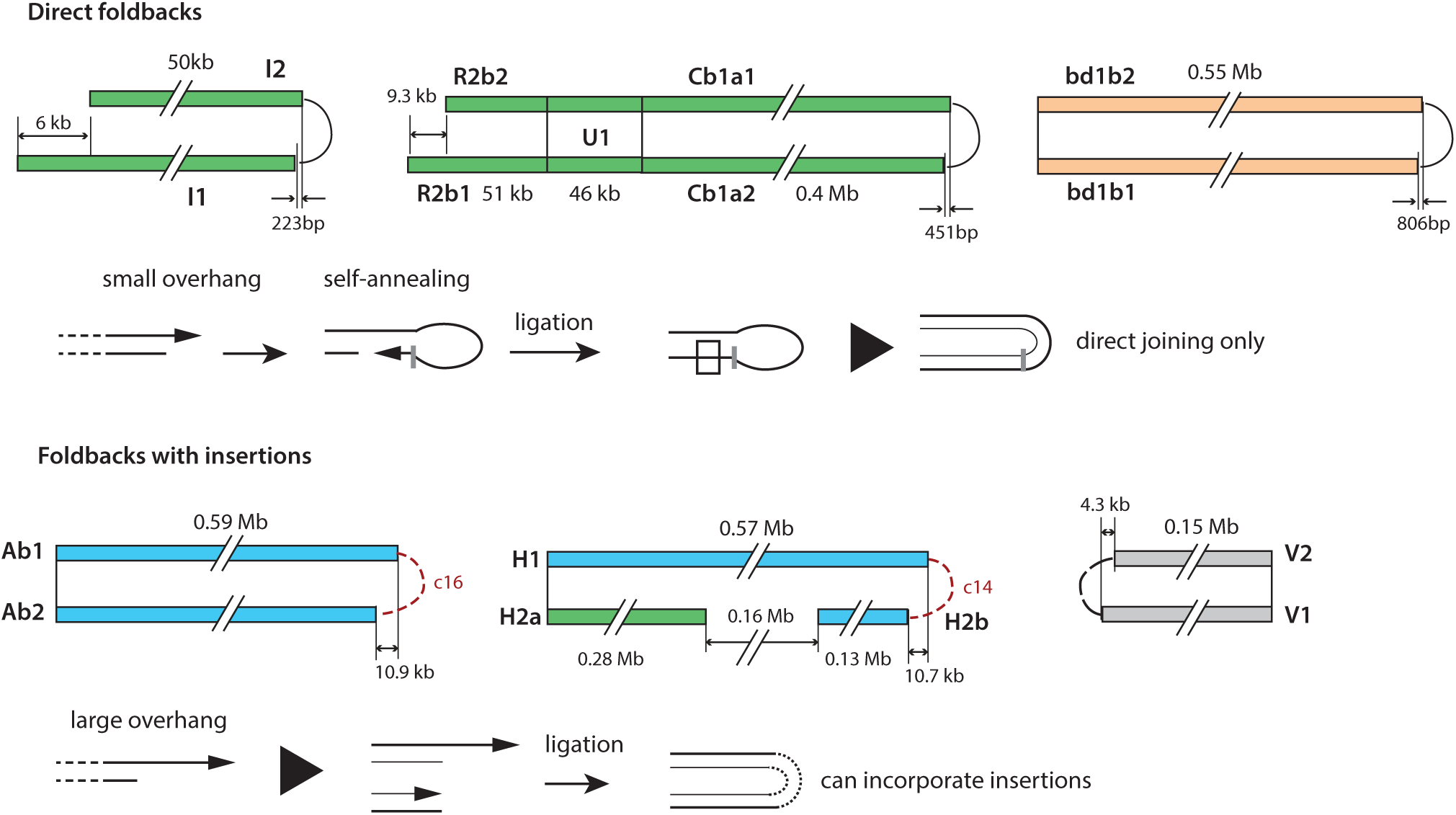
Foldback junctions between sister DNA segments. *Top*: Direct foldbacks often join breakpoints that are in close proximity (<1kb). For the foldback junctions between **I1**/**I2** and **Cb1a1**/**Cb1a2**, the foldback breakpoints are in *cis* with staggered breakpoints on the opposite side (left side of **I1**/**I2** and left side of **R2b1**/**R2b2**). The pairing of staggered breakpoints flanking sister duplications establishes the origin of these breakpoints as staggered dsDNA ends on an ancestral DNA fragment. *Bottom*: Foldbacks with insertions likely occur after the replication of ancestral dsDNA ends and can accept insertions. The sister breakpoints in insertion-mediated foldbacks are further apart. In the three examples shown here, the first two both contain multiple insertions (insertion chain 14 and 16), and the third contains a single insertion. *Note*: **Ab1**/**Ab2** are secondary segments only preserved in subclone **a6**; accordingly, the insertions in the **c16** junction are mapped to chr4:40.41-40.42Mb that is not adjacent to any ancestral breakpoints, but adjacent to insertions in the **c17** junction that is also private to **a6**. By contrast, the **c14** junction is inferred to be ancestral and all the insertions are mapped to regions near ancestral breakpoints (**Supplementary Table 2**); this inference is also consistent with the inference of **H2a**/**H2b** derived from an ancestral **H2** segment that is a sister of **H1**. See also **Extended Data Figure S2**.

**Extended Data Figure S8.**
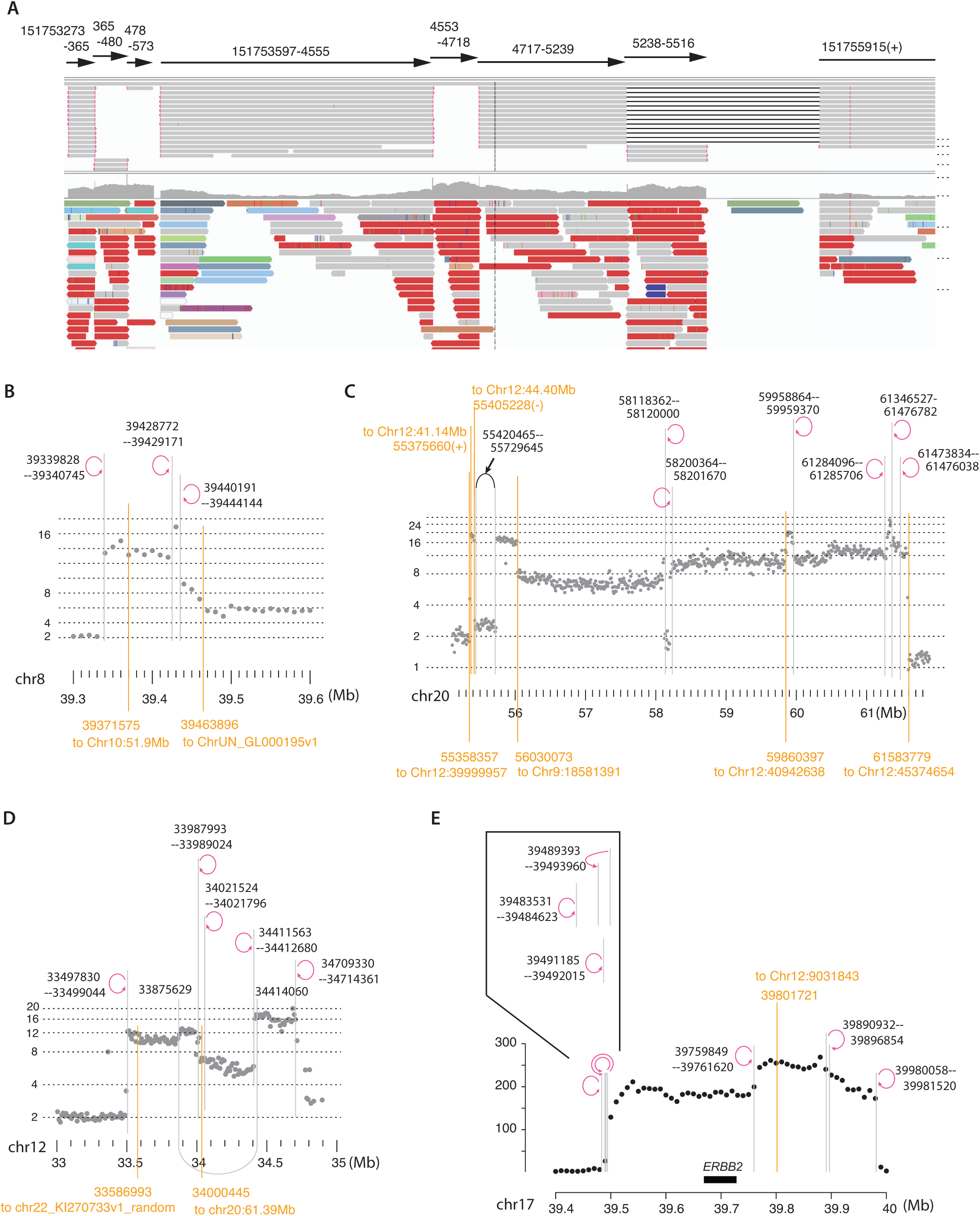
Tiling insertions and nested foldbacks in the HCC1954 genome. **A.** Seven insertions mapped to a region on chr1q near the breakpoint chr1:151755915(+). Two insertions, 151753273-365 and 151753597-4555 are joined in the same junction but at opposite orientations (violation of strand coordination). Insertion 151754717-5239 joins the breakpoint at 151755915(+). **B.** Total DNA copy number in a 0.3Mb region on 8p containing two breakpoints (orange) plus three foldback junctions. **C.** Haplotype-specific copy number (log2 transformed) in a 7Mb region on 20p. Note the presence of two pairs of adjacent breakpoints flanking the duplications between 55.36Mb (55.358Mb and 55.376Mb) and 55.42Mb (55.405Mb and 55.420Mb). **D.** Total DNA copy number (log2 transformed) in a 2Mb region on 12p, showing two breakpoints plus four foldback junctions. **E.** Six foldback junctions and one inter-chromosomal junction in the amplified region on 17q including the *ERBB2* gene.

**Extended Data Figure S9.**
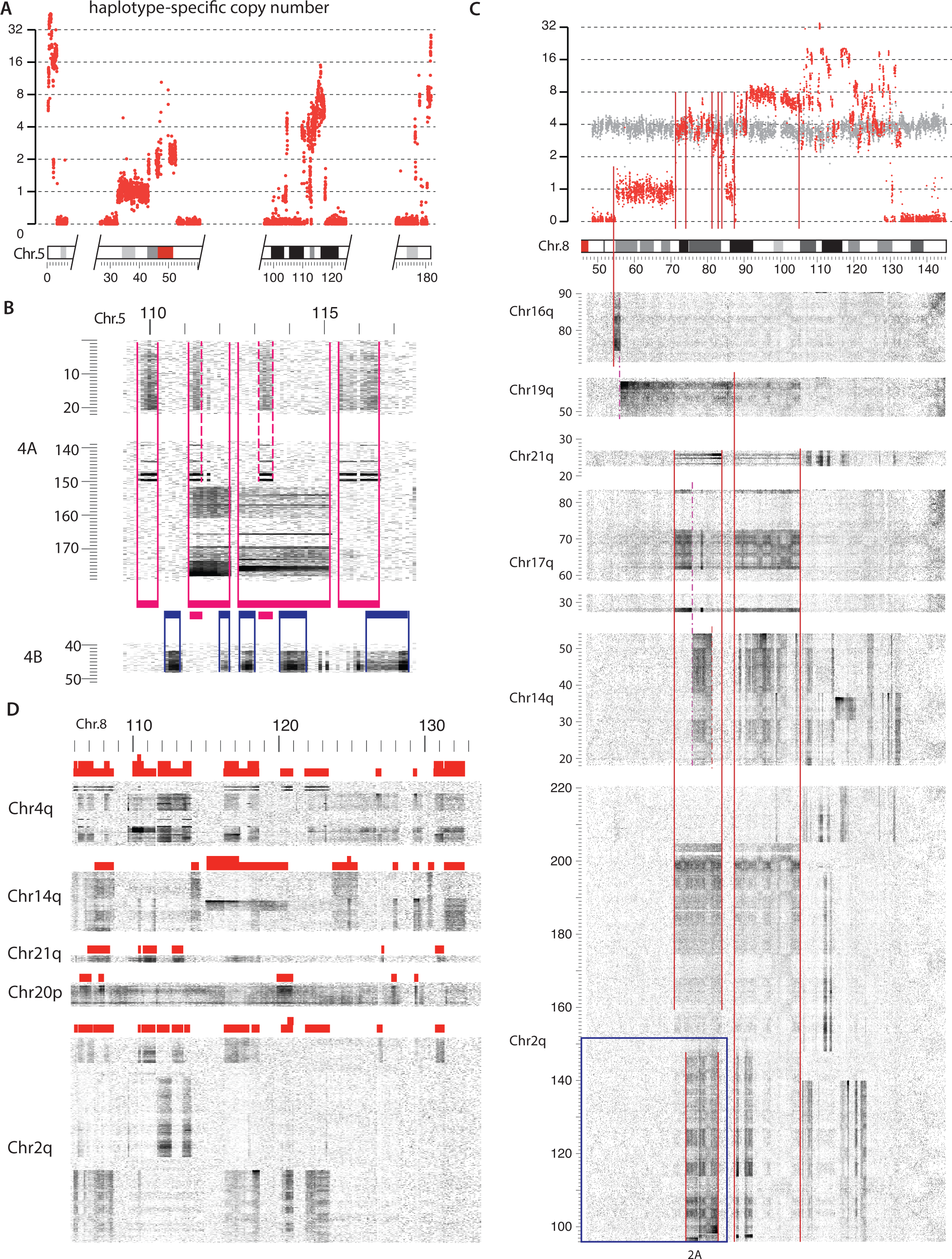
Evidence of ancestral chromosome fragmentation in regions of complex duplications. **A.** DNA copy number (log2 transformed) of the chr5 homolog with segmental amplifications. **B.** Hi-C contact map between chr5:109-116Mb and two chr4 homologs showing different segments from chr5 that are joined to different regions on chr4 identified by de novo *cis* contacts. The individual segments are shown as magenta (joining 4A) and blue bars (joining 4B). Partial overlap is only seen between two segments (magenta and blue); the remaining segments are either inclusive or non-overlapping as expected for fragments from a single chromatid. **C.** Haplotype-specific DNA copy number (log2 transformed) of chr8q (gray for the unaltered homolog, red for the homolog with amplifications). Hi-C contact maps between chr8q and various chromosomes reveal multiple segments that are either non-overlapping, or inclusive. **D.** Zoomed view of Hi-C contact maps between the amplified region on chr8q (106-132Mb) and other chromosomes. The scarcity of partial overlap between duplicated segments (the only exception being the segment from 115-121Mb that joins 14q) also supports an origin that these segments were originally derived from fragments from a single chromatid. In both **B** and **D**, the Hi-C contact density is normalized for haplotype-specific DNA copy number.

**Extended Data Figure S10.**
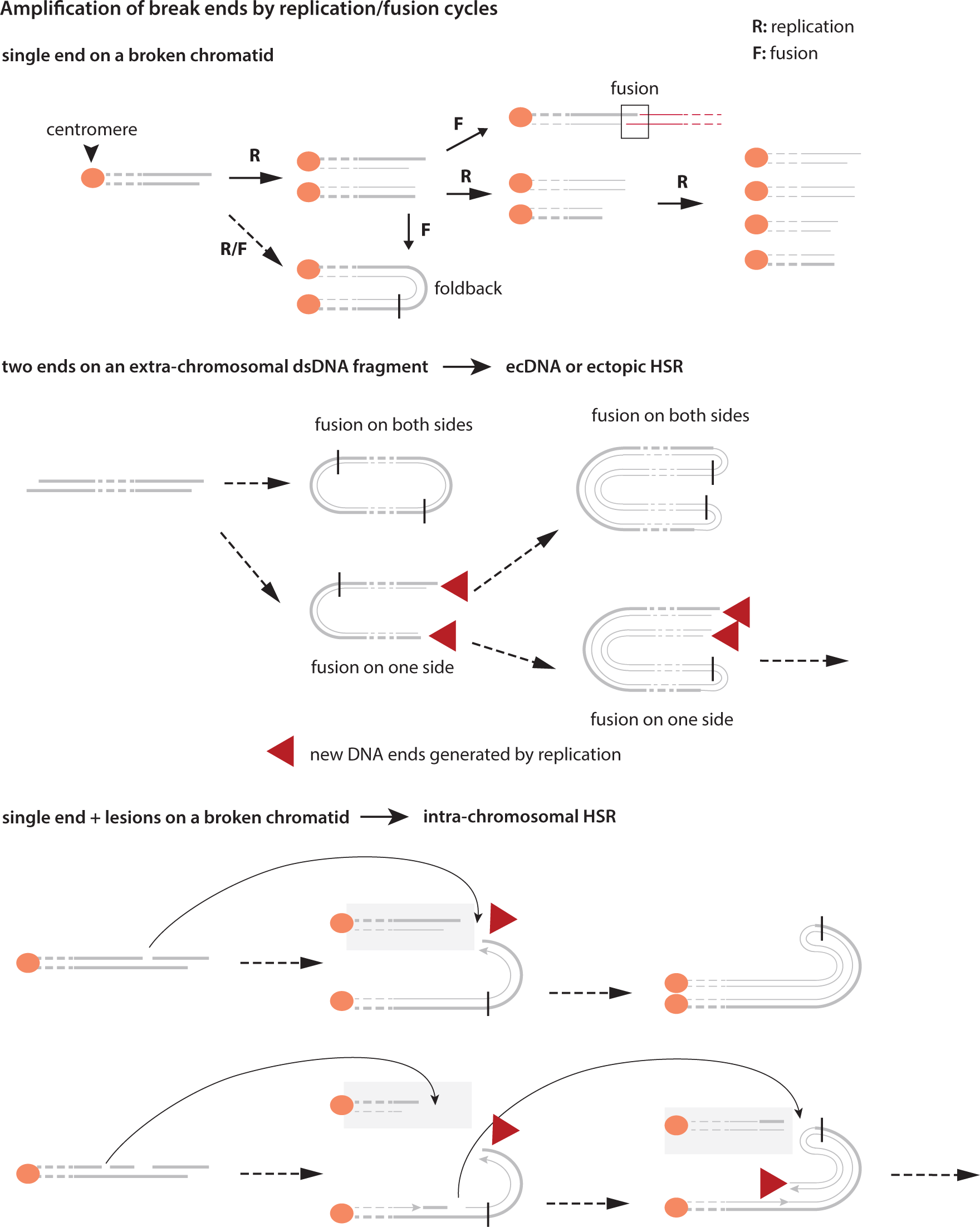
A proposed model for the amplification of a single ancestral DNA end into many adjacent parallel breakpoints over multiple rounds of break end replication. In the BFB cycles (top), the unligated chromosome ends are segregated into different daughter cells. Foldback junctions formed by self-ligation (black vertical line) are segregated into one daughter, but secondary breaks generated by subsequent bridge breakage events will be unrelated to the foldback breakpoints. For an extrachromosomal dsDNA fragment with two ends, the formation of a foldback junction at either end will cause sister segments to segregate into one cell. If both ends self-ligate, it will create a circular acentric chromosome (double-minute, or ecDNA circle). If only one end undergoes self-ligation, it will create an inverted duplication with two ends that can undergo further amplification iteratively. The amplicon will consist solely of foldback junctions and there should be adjacent foldback junctions generated from a single ancestral DNA end. The end of a broken bridge chromosome often contains ssDNA lesions (gaps or nicks) that can be converted into dsDNA breaks by replication. The regional concentration of foldback junctions or adjacent parallel breakpoints can therefore be explained by regional DNA breaks incurred on the initial bridge chromosome without successive cycles of bridge formation and breakage. In both the 2nd and the 3rd scenario, amplification is reiterated because each round of replication creates new DNA ends (red arrows) are not ligated before the next round of replication.

