## Supplementary Information for "Chromosome breakage-replication/fusion enables rapid DNA amplification"

##### This PDF file includes:

Supplementary text
SI Figures 1 to 20

### Supplementary Discussion and Data

#### 1. Definitions

We start by defining commonly used terminologies related to DNA rearrangement with examples shown in **SI Figure 1**.

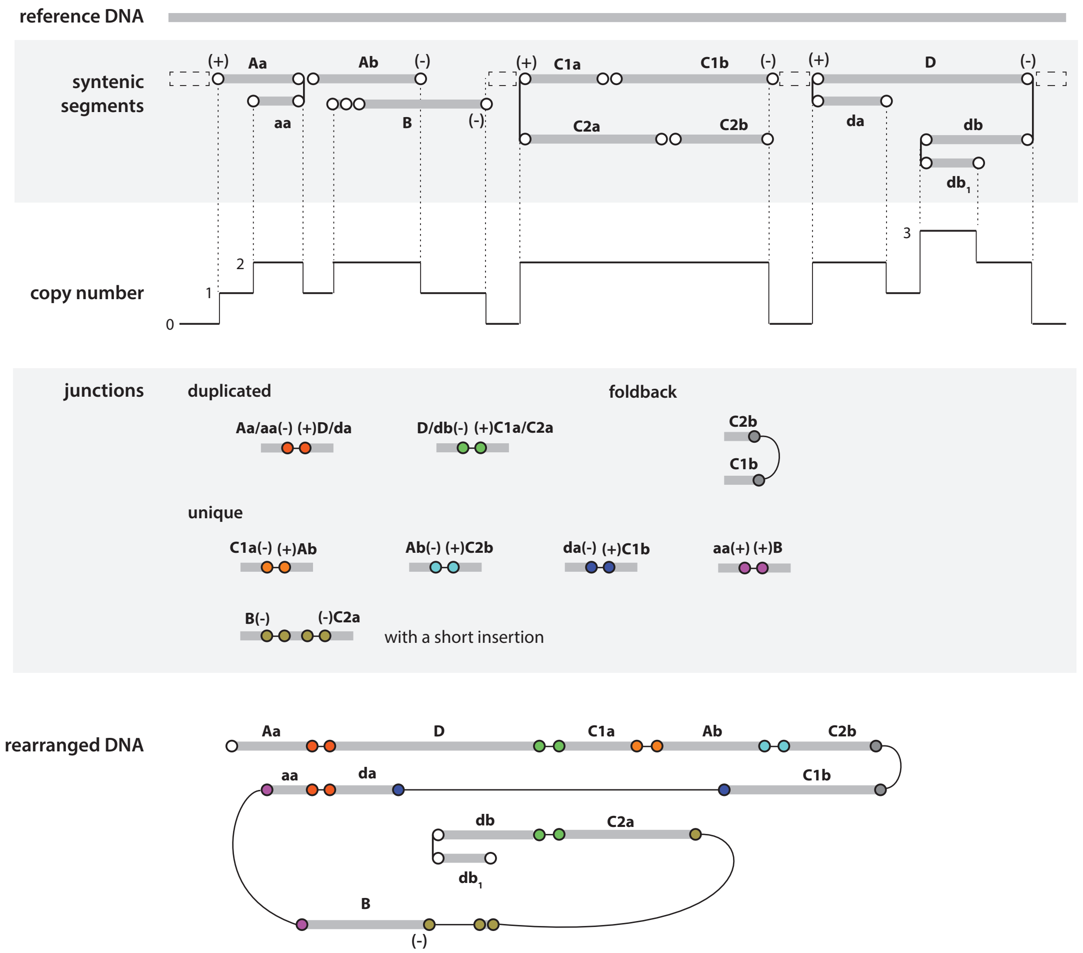


**SI Figure 1** An example of rearranged DNA (bottom) with segments (gray bars), breakpoints (open circles), DNA copy number, and junctions (colored circles). (+) and (-) denote the directionality of copy-number transition across the breakpoint. The naming of segments follows the same convention as in **Extended Data Figure 2**.

(i) **Rearrangement** A DNA sequence consisting of two or more subsequences (**Segments**) that are conserved in another (reference) sequence. See DNA rearrangement and "structural variant" and Bioinformatic analysis of DNA rearrangement.

(ii) **Segment** A DNA subsequence in the reference DNA that is conserved (i.e., contiguously aligned) in the rearranged DNA. Gray bars in **SI Figure 1**. See Segment and synteny mapping and **SI Figure 2** below.

(iii) **Breakpoint** The boundaries of segments define breakpoints (open circles in **SI Figure 1**). Breakpoints are represented by locations in the reference DNA. We use (+) and (-) to denote an increase or decrease of copy number at a breakpoint.

(iv) **DNA copy number** The copy number of a sequence in the reference DNA is the total number of segments from a rearranged chromosome or genome that are mapped to this locus. DNA copy number only takes integer values. But the bulk average DNA copy number in a heterogeneous population may take non-integer values, which is the algebraic average of the integer copy-number states weighed by their subclonal fractions.

(v) **DNA duplication** A sequence in the reference genome is duplicated if it is preserved in more than one segment (with different breakpoints) or multiple copies of the same segment in the rearranged genome. For cells with heterozygous chromosomes, only sequences derived from the same parental chromosome are considered to be duplicated, but not those from different parental (homologous) chromosomes.

(vi) **Junction** A DNA subsequence in the rearranged DNA that connects two rearranged segments (spans two breakpoints). These are shown as color-filled circles in **SI Figure 1**. Also see **SI Figure 3** below.

(vii) **DNA ends** Termini of single-strand (ss)DNA or double-strand (ds)DNA molecules. See Breakpoints and DNA ends.

(viii) **Adjacent breakpoints** Adjacent breakpoints are those in close proximity in the reference/pre-rearranged DNA such that they are unlikely to have been generated independently by chance.

(ix) **Insertion** A short DNA sequence inserted between large segments in the rearranged DNA. See Insertions and segments. A provisional criterion for distinguishing between insertions and segments is that insertions are flanked by *adjacent* (+) and (-) breakpoints in *cis*. For *cis* and *trans* breakpoints, see below.

(x) **Phasing of breakpoints, segments, and junctions** Phasing of breakpoints or segments refers to the inference of whether they originate from the same (*cis*) or different (*trans*) ancestral chromosomes. Phasing of junctions refers to the determination of the linkage and order of junctions in the rearranged chromosome. The *cis*/*trans* relationship between breakpoints or segments (with regard to the ancestral chromosome) is distinct from the *cis*/*trans* relationship of junctions (only relevant in the context of the rearranged chromosome). For example, in a reciprocal translocation between two chromosomes, the junctions are in *trans* as they are located on two different chromosomes, but they are formed between two pairs of *cis* breakpoints. By contrast, in a tandem duplication, there is only one junction but the two breakpoints can originate from sister chromatids; therefore, they are *trans* breakpoints.

(xi) ***cis* and *trans* breakpoints** *cis* breakpoints originate from a single ancestral DNA molecule; *trans* breakpoints originate from different DNA molecules or different copies of duplicated DNA.

(xii) ***cis* segments** DNA segments that are derived from fragments of a single ancestral chromosome by *fragmentation*. *cis* segments can be identified by *adjacent* (-) and (+) *cis* breakpoints that originate as opposite DNA ends after DNA fragmentation. In **SI Figure 1**, *cis* segments are labelled using lower case letters a, b, $\cdots$, e.g., **Aa**;**Ab**. Adjacency between breakpoints is, however, not strictly required for the inference of *cis* breakpoints or segments. See Evolutionary analysis of *cis* breakpoints and segments.

(xiii) ***trans* segments** DNA segments originating from different DNA molecules. The most common examples of *trans* segments are duplicated segments, e.g., **Aa**/**aa**, **Ab**/**B** in **SI Figure 1**. But two segments can be inferred to be in *trans* even if they do not overlap. For example, the **C1a**;**C1b**/**C2a**;**C2b** pair are in *trans*; therefore, **C1a** and **C2b** are in *trans* even if they do not overlap. This inference can be derived even if **C1b** is lost. See Evolutionary analysis of *trans* segments and breakpoints.

(xiv) **Mechanisms of DNA breakage ("cut")** Exogenous or endogenous processes that cause ss- or dsDNA breaks. Inferences about the processes of DNA breakage are drawn from the copy number, the nucleotide context, or the transcriptional and chromatin state of breakpoints.

(xv) **Mechanisms of DNA duplication ("copy")** Processes that produce extra DNA copies, including both semiconservative DNA synthesis, i.e., copying from both DNA strands as in normal DNA replication, and conservative synthesis, i.e., copying from only one DNA strand as in break-induced replication. Distinguishing between these two modes of DNA synthesis requires both the presence of *strand-specific* mutations in the DNA template and knowledge of their segregation in the duplicated DNA sequences. In the current study, we have taken advantage of strand-specific deamination and staggered ssDNA ends to demonstrate the origin of DNA duplication from semiconservative synthesis.

(xvi) **Mechanisms of DNA repair/recombination ("paste")** Processes that create rearranged DNA subsequences (junctions) by ligation (end-joining repair) or recombination (homology-dependent repair). Inferences about the processes of DNA repair are commonly drawn based on features of the junction sequence, e.g., extent of sequence homology, insertions, etc. However, the association between junction sequence features and different repair processes is promiscuous.

(xvii) **Mechanism of rearrangement** A cascade of molecular events that create one or more rearranged DNA molecules or chromosomes from the chromosomes of a single cell. A rearrangement mechanism (e.g., in immunoglobulin gene rearrangement) usually involves processes of DNA breakage (**cut**) and DNA repair (**paste**), but may also include DNA duplication (**copy**). A rearrangement mechanism may also reiterate itself over multiple generations (e.g., breakage-fusion-bridge cycles) to create multiple rearranged DNA sequences. In bulk DNA sequencing analysis, only a single snapshot of the rearranged sequence is available, causing uncertainty in both the evolutionary history and the mechanism. See Inference of the evolutionary history and mechanism of DNA rearrangement for further discussion.

#### 2. Further discussion on DNA rearrangement analysis

**A. DNA rearrangement and "structural variant".** A structural variant is a representation of the rearranged sequence as an *alteration* of the reference sequence (i.e., a variant). Common (simple) structural variants include deletions, duplications, inversions, transpositions, and translocations. There are also patterns of complex structural variants that cannot be reduced to combinations of simple structural variants. Here we opt to represent rearrangement using segments and junctions instead of structural variants because of the following reasons.

First, segments and junctions can be uniquely defined for any rearranged sequence based on its alignment to the reference sequence, but many complex rearrangement patterns (local $n$-jumps, chromothripsis etc.) can have more than one representations by simple or complex structural variants.

Second, the segments and junctions of a rearrangement always define a unique rearranged sequence, but a complex structural variant can represent more than one rearranged sequence. See Figure 1 of Li et al. (2020) for examples. This ambiguity is because structural variants are defined solely based on the joining pattern of breakpoints (i.e., junctions), but do not include breakpoint linkage information as illustrated in **SI Figure 2**.

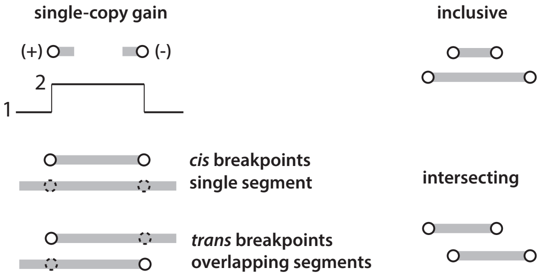


**SI Figure 2** Ambiguities of breakpoint linkage on rearranged segments. *Left*: *cis* and *trans* breakpoints associated with a single-copy gain. *Right*: Two segmental configurations reflecting different *cis* breakpoints.

Third, breakpoints and segments are directly associated with molecular features of the ancestral DNA, for which the evolutionary timing and correlation can be inferred by statistical analysis (e.g., adjacent breakpoints in **Figure 2**). By contrast, a structural variant often involves two or more breakpoints or segments; it is more difficult to infer the relative timing of different structural variants than the timing of individual breakpoints or segments.

Finally, breakpoints and segments provide more insight into the mechanism of rearrangement than structural variants. For example, tandem duplication (TD), a common structural variant, can arise in different sizes and by different molecular processes. It has been demonstrated that 10kb TDs are generated by the replication-bypass mechanism associated with BRCA1 deficiency, which produces a net gain of DNA. However, this mechanism does not produce larger TDs (>100kb or >1Mb). Moreover, as demonstrated both in a recent study of breast cancers (Setton, Hadi, Choo et al., 2023) and in the current study, replication bypass can also create non-tandem duplications. Therefore, the central genomic feature of the replication-bypass mechanism is the presence of two DNA segments with a 10kb overlap, but not a head-to-tail joining pattern of breakpoints as indicated in the TD classification.

The common usage of structural variants for DNA rearrangement analysis is at least partially due to the technical limitation that shotgun sequencing reads can only resolve junctions between breakpoints, but not their linkage on the rearranged DNA. With long-read sequencing and Hi-C sequencing, we can determine the linkage between breakpoints and resolve the segmental structure of rearranged DNA. Due to reasons listed above, it is advantageous to build a general framework for DNA rearrangement analysis based on the segmental structure instead of the conventional representation based on structural variants.

**B. Bioinformatic analysis of DNA rearrangement.** A complete description of DNA rearrangement includes three features: the rearranged segments, the junctions between segments, and the order of junctions and segments in the rearranged chromosome. Among these features, junctions can be efficiently detected by shotgun sequencing reads with non-contiguous (discordant) alignments to the reference. From the aligned subsequences of these reads, we can further identify breakpoints or segments that are shorter than the read length. In the absence of DNA duplication, all the rearranged segments are present at one copy and can be uniquely determined from consecutive (+) and (-) breakpoints on a chromosome.

When there are duplicated segments, determination of their segmental structure requires knowledge of the linkage between breakpoints flanking each duplicated copy (**SI Figure 2**). For small duplications, the breakpoints can be directly resolved from a single sequencing read or from a unitig (a unique path on the assembly graph) assembled from the reads. For large DNA duplications (≥100kb) with little sequence variation (the average frequency of de novo substitutions in somatic genomes is usually $\lesssim 10^{-5}$), even long read assembly cannot resolve the linkage between breakpoints. In the current study, we devised two strategies to derive the long-range linkage between breakpoints: The first is to link/phase breakpoints by segmental copy number variation (e.g., *cis* breakpoints always have the same copy number). This strategy can be applied both to a single genome with multiple copy-number states, and to multiple related subclones with variable copy-number states. The second strategy, which we applied to the analysis of a complex cancer genome, is to separate overlapping duplications based on their *cis* Hi-C contacts to segments from different chromosomes. Regarding the second strategy, it is in principle also possible to phase breakpoints based on the contacts of junction-spanning Hi-C reads, i.e., Hi-C reads containing the rearrangement junction sequences that can be determined from whole-genome sequencing. The most complete inference should combine both approaches and solving this technical problem will remove a major roadblock towards the assembly of complex cancer genomes.

**C. Segment and synteny mapping.** Rearranged segments are identified by aligning the rearranged sequence to the reference sequence. **Synteny** (blocks of genes with conserved order) mapping can be regarded as a special case of segment determination when alignment is performed at the gene level. Synteny mapping is used when comparing the genomes of *different species*, where significant sequence divergence interferes with the identification of long-range segmental conservation. For *somatic cells* from the same individual, the segments of rearrangement can be directly identified by sequence-level alignment. However, the alignment should distinguish between segments derived from different homologous chromosomes based on the *haplotype phase*. For duplicated sequences that are highly similar, their rearrangement can only be determined based on the non-duplicated flanking sequences. An example can be found in the determination of segmental gains in 39-41Mb in bridge clone **a** presented in Breakpoints of rearranged segments in bridge clone **a**.

**D. Breakpoints and DNA ends.** A DNA end is a *transient* molecular structure on the ancestral chromosome that can only be identified by *in situ* DNA sequencing. A breakpoint describes a feature of the rearranged DNA sequence that can be either retained by a single chromosome or preserved in a clonal population; therefore, breakpoints can be identified by either single-cell or bulk DNA sequencing.

There are several key differences between breakpoints and DNA ends. First, a double-strand (ds)DNA end consists of two single-strand (ss)DNA ends that can be either flush (same position for both strands) or staggered (different positions on opposite strands); a breakpoint is a sequence feature that is identical for both DNA strands. Second, DNA ends are not duplicated exactly due to the end-replication problem or end processing such as resection; breakpoints (when they descend from ancestral DNA ends) are features of the ligated sequence and are duplicated exactly. Finally, in addition to arising from dsDNA ends, breakpoints can also arise from an intact DNA donor template that is invaded by a dsDNA end. See **SI Figure 3**. This strand invasion followed by DNA synthesis can create a rearranged segment from the intact donor, with breakpoints corresponding to the sites of invasion or template switching, but not sites of DNA breakage. Given the uncertainty about the origin of individual breakpoints, an integrative analysis of all the breakpoints is necessary to infer whether a breakpoint originates from DNA breakage or strand invasion, and to determine the relationship between the breakpoint and the ancestral DNA end.

**E. Insertions and segments.** Putative insertions are derived from foreign DNA (i.e., not from the host genome), such as viral DNA or integrated transgenes. Insertions are also used to refer to short sequences sandwiched between large segments, even if the inserted sequences originate from the reference. An example is a short segment (unlabelled) to the left of segment **B** in **SI Figure 1**. When the inserted sequence can be mapped to one or multiple locations in the reference (in the latter case, these sequences represent repeats), it is often referred to as being "templated" (or more precisely, mappable). However, there is no difference between templated insertions and segments in the representation of rearranged DNA sequence: <u>Any subsequence that can be mapped to a location in the reference DNA should be viewed as a rearranged segment regardless of its size.</u>

Although there is no difference between templated insertions and segments from the perspective of bioinformatic analysis (both are determined by sequence alignment), a distinction may be made from the *evolutionary* analysis of breakpoints. For short insertions, the close adjacency of breakpoints of the inserted sequence suggests that they are generated in a concerted manner, at the same time. For large segments, the breakpoints may arise from independent events occurring at different timepoints.

In the current study, we use 'insertions' to refer to short sequences for which the breakpoints are closer than expected (i.e., "adjacent breakpoints" in **Figure 2**). In the experimentally generated clone (bridge clone **a**), we chose an operational threshold (10kb) to distinguish insertions from large segments. This classification is supported by the distinct origins of short insertions and large segments in the reference chromosome, as well as their distinct joining patterns in the rearranged chromosome. In the HCC1954 cancer genome, we identified 20 segments between 10 and 25kb in addition to 350 insertions that were shorter than 10kb. Whether these 'long' insertions are generated by similar or different processes from the short insertions is unclear. A systematic analysis of rearranged segments in large cancer genome datasets is required to both refine the criteria for identifying insertions and to generate hypotheses about their mechanisms.

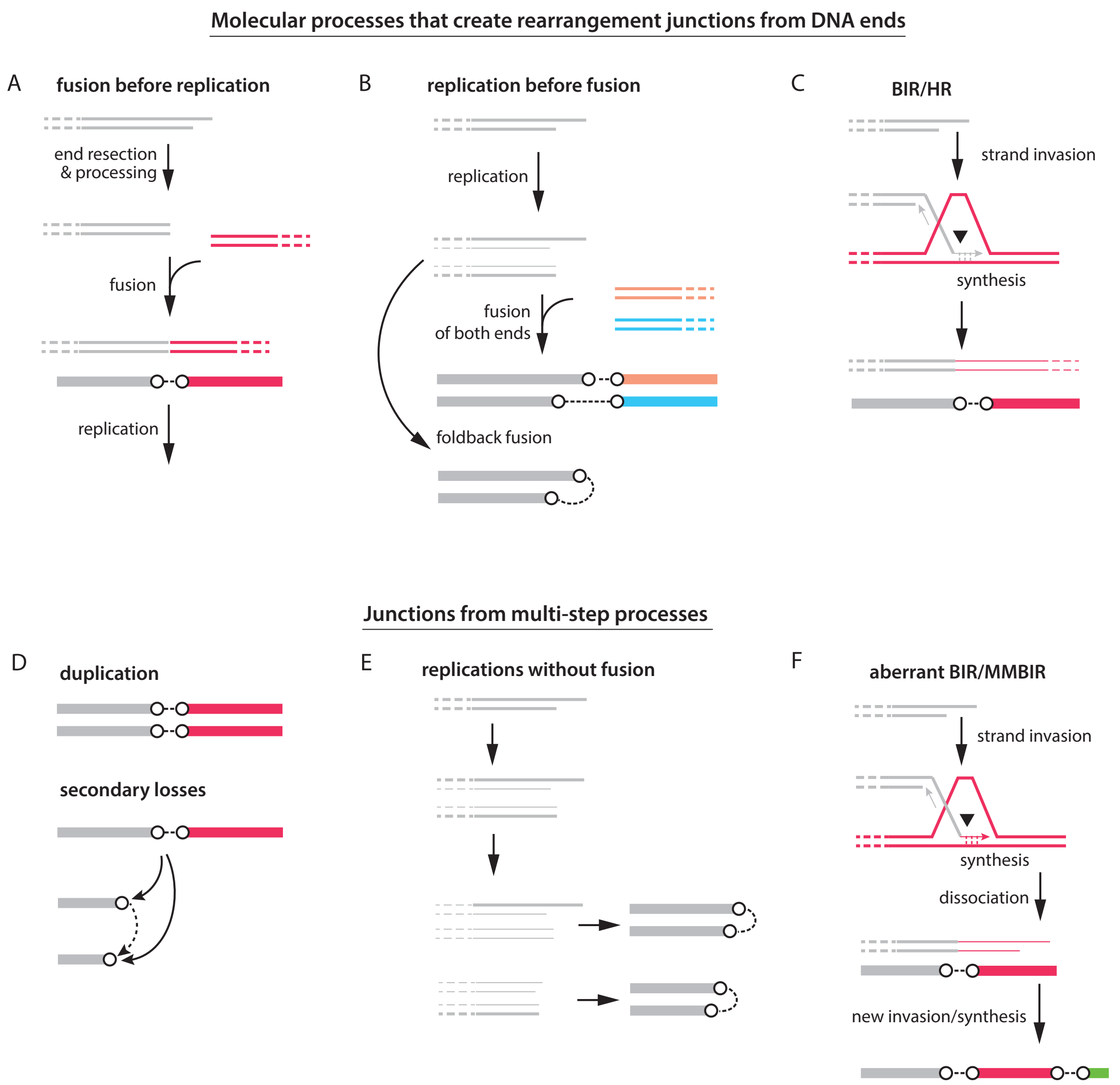


**SI Figure 3** Molecular processes that create rearrangement junctions.
**A**. Fusion (end-joining repair) between two DNA ends can create a single junction with two breakpoints. Each breakpoint originates from a single ancestral dsDNA end.
**B**. An unrepaired dsDNA end may be duplicated by replication and the sister DNA ends may form separate junctions. A special case is when sister DNA ends join each other to form a foldback junction. Newly synthesized DNA strands are shown as thin lines; template DNA strands are shown as thick lines
**C**. A dsDNA end may also invade an intact DNA template and initiate synthesis by break-induced replication (BIR). The 3'-end may be extended continuously, but the 5'-end cannot. Studies of BIR in yeast suggested that second strand synthesis uses the leading strand as template (i.e., conservative synthesis); however, whether the same process also occurs in mammalian cells, and whether BIR is always conservative remain unclear. BIR may be resolved in different ways, but the junction created by the initial strand invasion event consists of two breakpoints, one originating from a dsDNA end, the other from strand invasion. See Rearrangement outcomes of break-induced replication.
**D**. Junctions/breakpoints can be duplicated or deleted.
**E**. Replication of naked DNA ends over multiple generations can create two or more adjacent DNA ends as seen in regions of DNA amplification. See Clusters of adjacent parallel breakpoints and foldback junctions.
**F**. Microhomology-mediated break-induced replication, or MMBIR, posits that the BIR fork can dissociate from one template (red) and invade a different template (green), creating a series of junctions by "copy-and-paste."
Note that breakage-fusion (**A** and **D**), BIR (**C**), or MMBIR (**F**) can produce similar breakpoint/junction patterns, but adjacent parallel breakpoints are a specific feature of breakage-replication-fusion (**B** and **E**).

**F. Evolutionary analysis of *cis* breakpoints and segments.** For rearranged DNA that is preserved in a clonal population, it is impossible to definitively exclude the possibility that two segments or two breakpoints originate from two ancestral DNA molecules (e.g., sister chromatids). However, two strategies may be employed to infer *cis* breakpoints or *cis* segments indirectly.

The first strategy is by the identification of adjacent gapped or overlapping breakpoints (**Figure 2**). The *cis* relationship is established by two lines of reasoning: (1) the distance between these breakpoints suggests that the probability that they are generated independently on different DNA molecules is very small (an evolutionary argument); (2) there is ample evidence that such breakpoints can result from opposite ends created by a single DNA breakage event (a molecular argument). As adjacent *cis* breakpoints are inferred to have been generated at the same time, their partners in the rearrangement junctions are also generated at or around the same time. Iteration of this strategy can identify evolutionarily concurrent breakpoints even when they are not adjacent in the original chromosome.

As an example, we consider three segments as shown in **SI Figure 4**. For the proximal breakpoints on segment 1 and 2, if the breakpoint distance $d_1$ is very small compared to the size of the segments ($L_1$ and $L_2$), we can infer that the two breakpoints are generated from a single breakage event. Similar argument can be applied to breakpoints between segment 2 and 3 (when $d_2 \ll L_2 + L_3$). Moreover, based on the junctions between breakpoints (same color for each junction) in the rearranged chromosome (bottom), we can infer that the breakpoints of segment 3 are also generated concurrently as the adjacent breakpoints between segment 1 and 2. Note that the inference of adjacent *cis* breakpoints cannot establish the long-range *cis* relationship between breakpoints on each segment. The latter can only be established if the size of the segment (e.g., $L_2$) is very small (i.e., it is a short insertion) compared to the average distance between breakpoints. But even without knowledge of the long-range *cis* relationship between breakpoints, we can still identify adjacent breakpoints based on breakpoint distance.

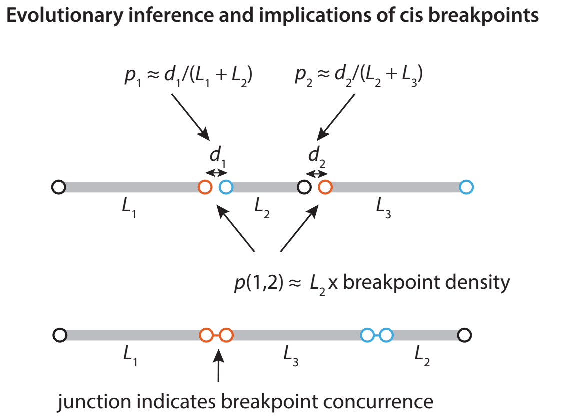


**SI Figure 4** *cis* breakpoints/segments from DNA fragmentation.

We can also assign *cis* breakpoints or *cis* segments based on the absence of evidence for a *trans* relationship. By definition, the presence of *trans* breakpoints or segments requires duplicated DNA segments. In the absence of DNA duplication, each segment or breakpoint can be parsimoniously assigned to a single ancestral chromosome and therefore taken to be in *cis*. This line of reasoning is weaker than the inference based on adjacent breakpoints: the breakpoints can be generated either simultaneously or sequentially.

A well-known example of *cis* segments is in chromothripsis, where all the segments are derived from fragments generated in a single catastrophe. The oscillation between deletion and retention suggests all the retained segments are derived from the same ancestral chromosome, but does not establish that they are generated all-at-once. The argument that the rearrangements in chromothripsis arises all-at-once instead of by gradual evolution was originally made by contrasting the simulated copy-number outcomes of gradual evolution with the observed two-state oscillating copy-number pattern (Stephens et al., 2011). A counterargument to this original argument was made subsequently by Kinsella, Patel, and Bafna (2014). We later provide a revised argument supporting the all-at-once model of chromothripsis in Inference of the evolutionary history and mechanism of DNA rearrangement.

**G. Evolutionary analysis of *trans* segments and breakpoints.** Here we only consider *trans* segments/breakpoints on duplicated segments. For simplicity, we do not consider pre-existing duplications in the reference chromosome and assume all the segments of the rearranged chromosome are mapped to unique locations in the reference. We want to infer (1) the phylogenetic relationship between duplicated segments; (2) the timing of breakpoint formation on duplicated segments. These inferences can be drawn based on three types of variation that distinguish different duplicated copies: (1) local sequence variants; (2) different breakpoints; (3) different flanking sequences.

*Mutations* The evolutionary timing of duplicated segments can be inferred from the number of mutations generated after duplication. These mutations should accrue independently in each duplicated copy (e.g., assessed by the positions of mutations). As the average frequency of de novo mutations is generally low ($\lesssim 10^{-5}$), the most useful scenario is when there is hypermutation (e.g., kataegis). See Origin of insertions that share a breakpoint with large segments for examples.

*Breakpoints* The following rules apply to parallel breakpoints on duplicated segments (**SI Figure 5**). (1) Each breakpoint is generated only once at a specific timepoint (a direct consequence of the infinite-sites model of genome evolution). See Ma, Ratan, Raney et al. (2008). (2) Flush breakpoints (identical position and identical junction sequence) are generated before duplication. (3) Adjacent parallel breakpoints are generated by replication through an unrepaired dsDNA end; therefore, they are generated at the same time as the (initial) duplication. Most adjacent parallel breakpoints are staggered, but they may have identical positions by chance; in both cases, the junction sequences are different because they are generated by ligation to different partners. (4) For non-adjacent parallel breakpoints, the shorter breakpoint can be derived from a duplicated segment with the longer breakpoint by a secondary deletion, but the longer breakpoint cannot be derived from a duplicated segment with the shorter breakpoint by a secondary extension. The reasoning is as follows. First, as deletion is irreversible, a short DNA segment can be derived from a long segment by deletion but cannot produce a long DNA fragment by extension ("reverse" deletion). Second, although it is possible to extend a DNA end by homologous recombination (HR), this process can only generate one breakpoint that extends the ancestral DNA end, but cannot create two parallel breakpoints. Finally, when an ancestral DNA end is duplicated into two sister DNA ends by replication, neither end can be extended by HR as there is no intact sister chromatid to be used as the template for HR.

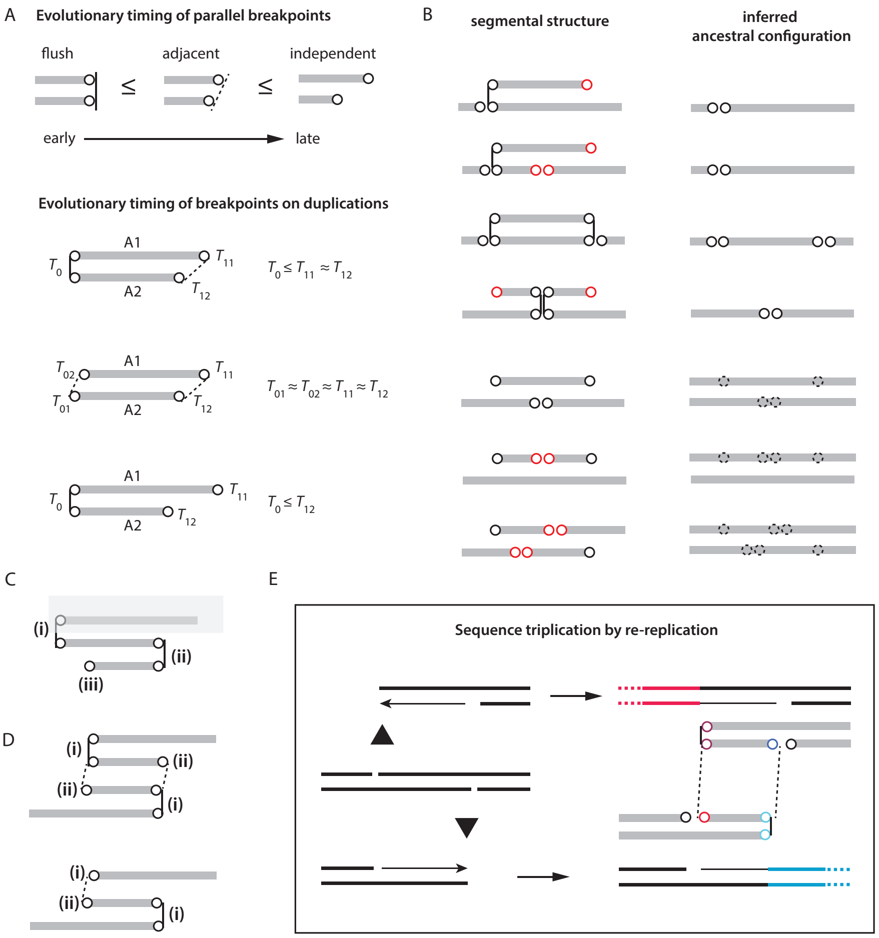


**SI Figure 5** Timing of *trans* breakpoints on duplications. **A**. Relative timing of parallel breakpoints on duplications. **B**. Different segmental configurations of single-copy duplications with the inferred ancestral segments (right) and secondary breakpoints (red circles). The top four suggest a chromosome breakage event before duplication; the bottom three suggest a duplication before breakage. **C**. An example of a non-flush breakpoint (i) that is generated before the flush breakpoints (ii). The long breakpoint (i) is initially duplicated but becomes a single breakpoint after deletion of the long segment (shaded box). Note that the short breakpoint (iii) is generated after the flush breakpoint. **D**. A special scenario when adjacent parallel breakpoints (i and ii) are generated sequentially by two rounds of replication as shown in **E**. **E**. The replication forks originating from the middle segment collapse at the nicks on opposite strands, creating adjacent ssDNA nicks and dsDNA ends. Re-replication (through replication forks fired from the partner segments) can create up to two internal segments whose breakpoints are adjacent to the breakpoints of the two overlapping segments. However, each internal segment must share exactly one flush breakpoint with the two long segments. This constraint excludes both *trans* segment configurations shown in **Figure 1B**.

The relative timing of *trans* breakpoints can be summarized as (**SI Figure 5A**):

$$T[\text{flush breakpoints}] < T[\text{replication}] = T[\text{staggered breakpoints}] \le T[\text{short non-adjacent breakpoint}].$$

Applying these constraints to single-copy gains with various breakpoint configurations (**SI Figure 5B**), we can infer the relative timing of duplication and breakpoint generation. The example in **SI Figure 5C** shows why the timing constraint applies only to the short breakpoint but not the long breakpoint when both are in *cis* with flush breakpoints.

*Segments* We can apply the timing constraint of breakpoints to segments (**SI Figure 6**). (1) For two segments in an 'inclusive' configuration (i.e., one is contained in another), the shorter segment is either a descendent of the longer segment, or the two segments arise independently from two precursor duplications. These two scenarios cannot be distinguished. (2) For two segments that are partially overlapping ('intersecting'), they must have been generated independently from two DNA copies, which indicates either independent breakage events on duplicated DNA copies, or independent copy-and-paste events. (3) For two segments with identical breakpoints, their timing can only be inferred based on mutations generated after duplication or by distinct breakpoints in the flanking segments.

We next consider the possible configurations of triplications. For the *trans* configurations shown in **Figure 1B**, the timing constraint implies that all the breakpoints must have been generated at the same time; this requires re-replication as there are three overlapping segments. The only possible outcomes are shown in **SI Figure 5D**; these outcomes can arise from the hypothetical model shown in **SI Figure 5E**. In this model, two partially overlapping *trans* segments are generated by replication through ssDNA gaps on opposite strands. Up to two internal segments (thin lines) can be generated by re-replication by replication forks fired from the unreplicated segments (red on top; blue on bottom). Note that to enable the replication fork to pass through, both strands have to be ligated; therefore, the short segment has to share a flush breakpoint with one of two long segments, but not both. Thus, neither configuration of *trans* segments as shown in **Figure 1B** is possible. Additional examples of triplications are shown in **SI Figure 6**.

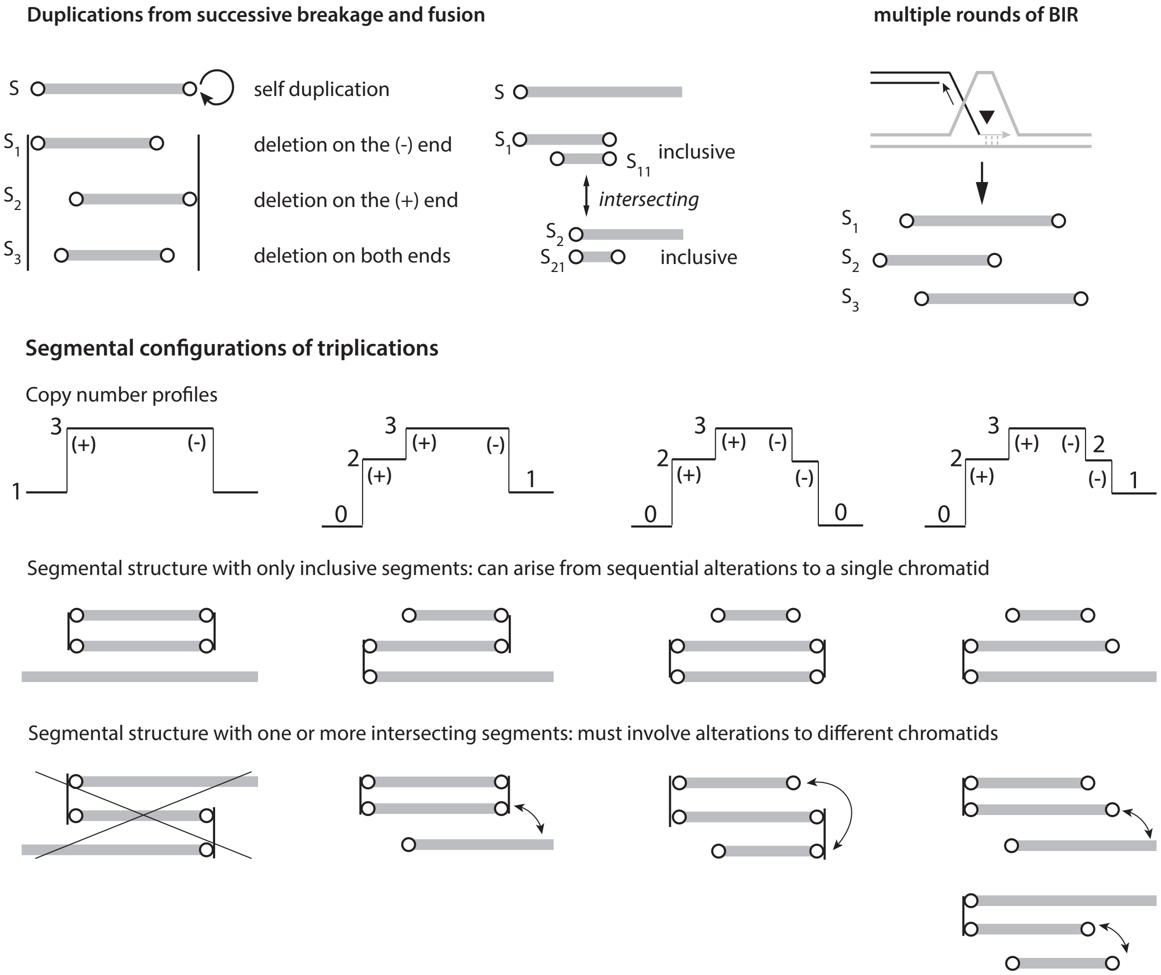


**SI Figure 6** Segmental configurations of multi-copy gains. *Top left*: Configurations of breakpoints on secondary duplications generated by breakage and fusion. Each secondary duplication is shorter than its ancestor ("inclusive"); two segments with partial overlap ("intersecting") must have descended from different ancestors (different strands of a single dsDNA fragment, or independent dsDNA segments). *Top right*: Duplications generated by break-induced replication from an intact donor template can be partially overlapping as the breakpoints generated in different rounds of BIR are independent. *Bottom*: Examples of segmental configurations of triplications. The first row show configurations without partially overlapping segments; the second row show configurations with partially overlapping segments. Note that the first example of triplication is not permitted based on the model shown in **SI Figure 5**.

**H. Inference of the evolutionary history and mechanism of DNA rearrangement.** The question of how a rearranged DNA sequence arises is often addressed from two angles. The first is to infer the evolutionary history of rearrangement, i.e., to deduce the series of sequence alterations that create the rearranged DNA sequence from the reference sequence. The second is to infer the mechanism of rearrangement, i.e., to provide a mechanistic explanation of the rearranged sequence using a cascade of molecular events that can involve DNA breakage, DNA repair, or DNA replication.

These two problems are different but intimately related. Knowledge of the rearrangement mechanism will help identify genomic features that are produced in a single cascade of events, which simplify the search of evolutionary trajectories that produce complex rearrangements. On the other hand, sequence features that are inferred to be evolutionarily concurrent can lead to specific mechanistic inferences or hypotheses. One such example in our current study is adjacent parallel breakpoints that indicate replication of unrepaired dsDNA ends.

Here, we argue that to draw evolutionary or mechanistic inferences about DNA rearrangement does not require knowledge of the complete evolutionary trajectory that produces the rearranged sequence. Instead, we can draw inferences directly from the sequence features, i.e., breakpoints, junctions, and segments of rearranged DNA.

As an example, we revisit the genomic evidence that suggests rearrangements in chromothripsis arise in a single catastrophe instead of by gradual evolution. The original argument was based on the prediction that a random series of deletions, tandem duplications, or inversions as inferred from the junctions of the rearranged DNA will produce DNA duplications. This prediction contradicts the observation that the rearranged chromosome only shows two copy-number states, retention and deletion. A caveat in this otherwise elegant argument is that it implicitly assumes that head-to-tail junctions reflect tandem duplications. However, the head-to-tail junctions can also arise from two inversions or a transposition, both of which do not cause copy-number gains. If we accept these alternative interpretations of the junctions, then it is straightforward to come up with a series of inversions and deletions, or a series of transpositions and deletions, that produce only deletions but not duplications. Therefore, the argument for chromothripsis as a one-off catastrophe hinged on one of many possible interpretations of rearrangement junctions.

Here we provide an evolutionary argument for the one-off model of chromothripsis that is solely based on features of the rearranged DNA including segments, breakpoints, junctions, and does not invoke the "structural variant" interpretation of rearrangement junctions. First, the concentration of breakpoints on a single chromosome cannot be explained by random DNA breakage that is expected to affect all chromosomes. Such concentration can only arise from an *unstable* chromosome. The question is whether these breakpoints accumulate over multiple generations (cell division) or in one generation. Second, if the unstable chromosome is replicated completely but unevenly segregated between the daughter cells, it should create both copy-number gains and losses. (By definition, an unstable chromosome cannot be stably inherited by both daughters.) Note that this argument for copy-number gains based on uneven DNA segregation is different from the original argument where duplication was derived from an interpretation of the head-to-tail junction. Third, if the unstable chromosomes undergoes deficient replication such that deletion is the predominant outcome in both daughters, then the breakpoints of each deletion (due to deficient replication) are evolutionarily concurrent. Moreover, the formation of junctions between the breakpoints of different deletions suggests that these breakpoints are generated concurrently. Therefore, the copy-number, breakpoint, and junction features of chromothripsis suggests a process that creates multiple deletions in one generation.

We can extend this analysis to complex rearrangements with copy-number gains. We first consider a *single* duplication. This region is contained in two segments that can be either inclusive or intersecting. If the segments are intersecting (i.e., with partial overlap), they must have been derived independently, e.g., from fragments from sister chromatids. If one is contained in the other (inclusive), then either they arose independently, or the short segment was derived from a duplicated copy of the long segment. Now we consider *all* duplicated regions. If they were generated independently from sister chromatids, we would expect to see at least some instances of intersecting segments. If, however, only inclusive segments are observed, then we can conclude that the sister-chromatid fragmentation model is unlikely.

For regions with even higher copy-number states, we expect many of them to be contained in three or more different segments. If these breakpoints are derived independently from different DNA copies, there is a high chance that at least some of the segments have partial overlap. To avoid partial overlap, the duplicated segments must be arranged in a hierarchy (like Russian dolls). This feature can be recognized in the amplified regions on chr5 and chr8 in the HCC1954 genome (**Extended Data Figure 9**). The hierarchical pattern of amplified segments cannot be explained by sequential duplications and deletions, because independent duplications of an ancestral segment can produce truncated segments that have partial overlap (see **SI Figure 6**). Therefore, the hierarchical pattern suggests a chronological order of duplications, i.e., the second largest segment is derived from the largest segment, and the third largest is derived from the second largest (but not the largest), etc. Based on this argument, we speculate that the segments in the amplified regions as shown in **Extended Data Figure 9** are generated by a re-replication process that produces an onionskin-like pattern. See Copy-number gain and amplification after micronucleation.

The above analysis of copy-number gains can only be made when the segmental structure of duplicated DNA segments can be determined. Such information cannot be obtained from shotgun sequencing, but can be derived from long-range sequencing data such as Hi-C and strand-seq. We therefore expect these data to be critical for both the bioinformatic analysis and the mechanistic interpretation of cancer genomic rearrangements.

#### 3. Breakpoints of rearranged segments in bridge clone a

In this section we show how the breakpoints of rearranged segments in bridge clone **a** are determined from copy-number variation in the subclones. We first show the DNA copy number of haplotype A in different subclones (**a1-a6**) together with the segmental structure (shown only once for samples with identical copy-number profiles). A summary of segments identified in all subclones and the rationale for the segmental decomposition are presented afterwards.

Figures in this section are not numbered but ordered by regions. The subclones are ordered based on the number of segments they retain: **a1** has the fewest number of segments and **a6** has the most. Segments are colored by the subclone where they are present at the **lowest** copy number state: gray for **a1**; yellow for **a2** and **a3**; orange for **a4**; green for **a5**; blue for **a6**.

**Copy-number variation in bridge clone a: p-ter–16.5Mb**

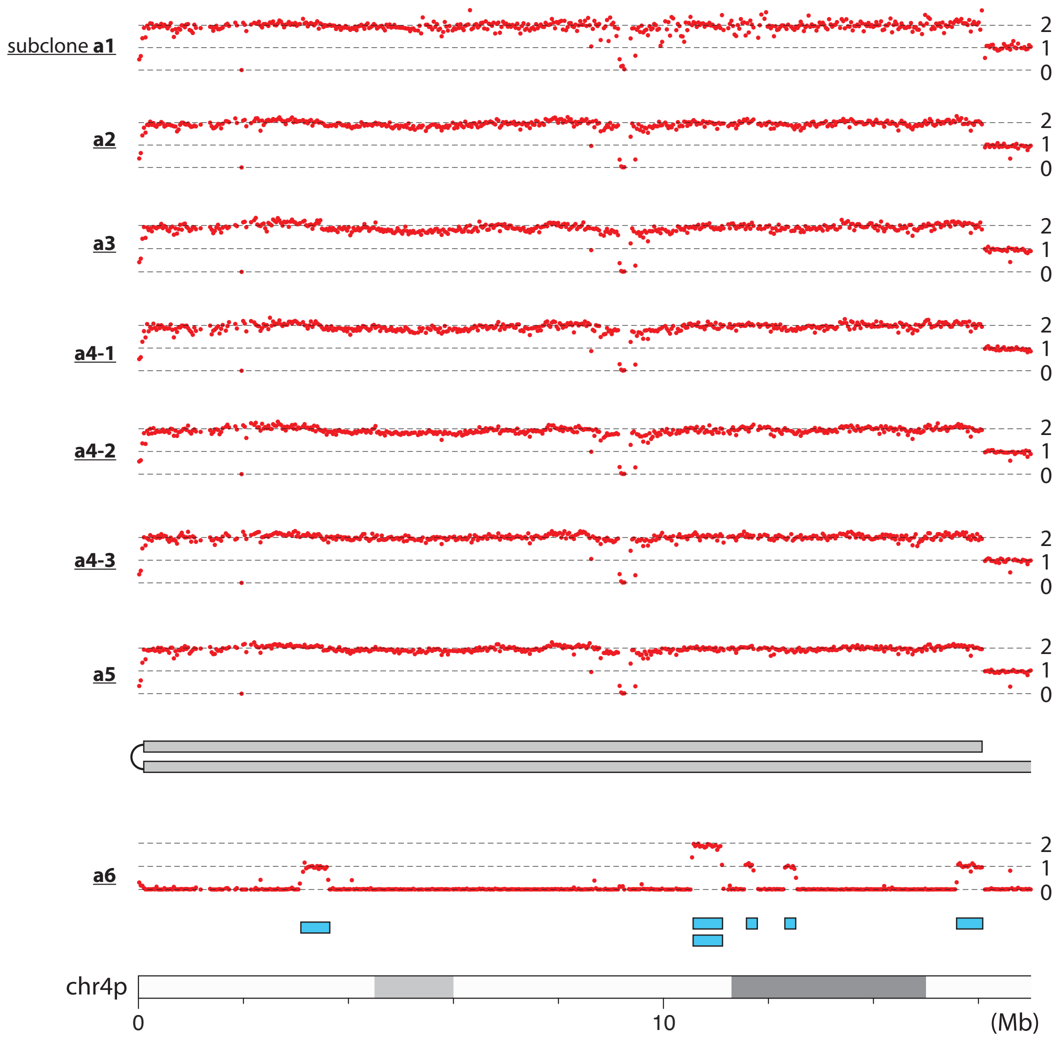


**Segmental structure of copy-number gains: p-ter–16.5Mb**

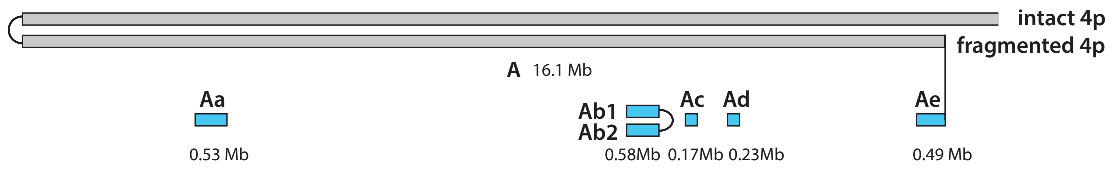


**a1-a5** have the same copy-number pattern and segmental composition; segments in **a6** are generated by a secondary chromothripsis event that also involves chr14q.

**Copy-number variation in bridge clone a: 17-22Mb**

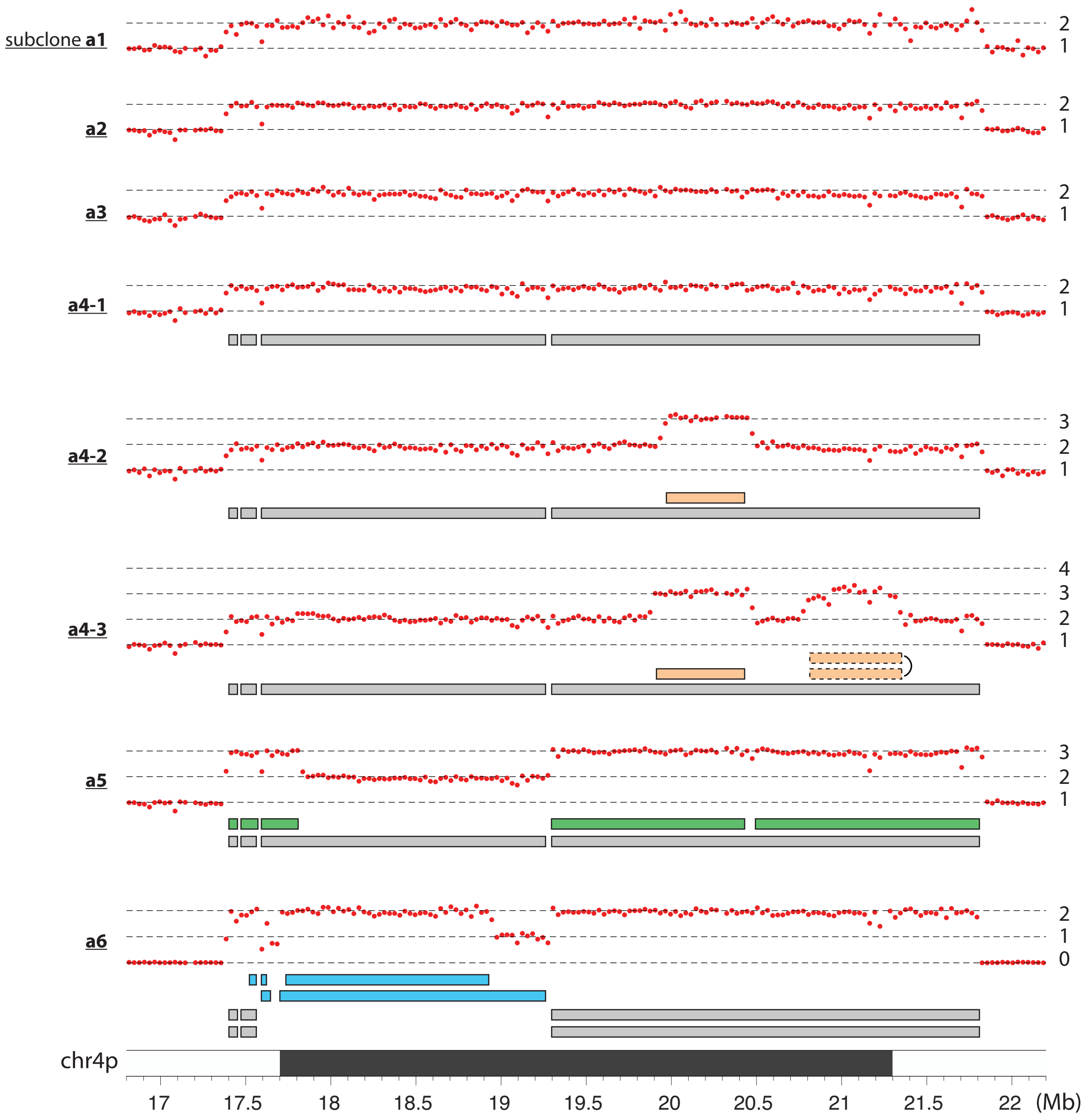

**Segmental structure of copy-number gains: 17-22Mb**

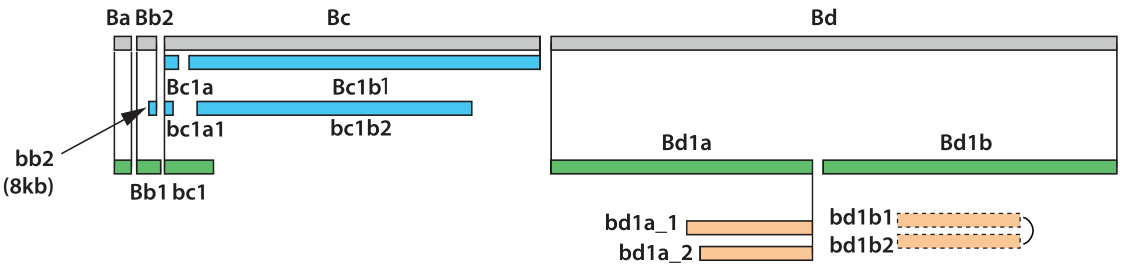


Segments **Ba**,**Bb2**, and **Bd** are defined in subclone **a6** where the flanking sequences are deleted. The same reasoning leads to the definition of *cis* segments **Bc1a** (17.60-17.66Mb) and **Bc1b1** (17.68-19.26Mb), and their descendants/cousins **bc1a1** (17.60-17.63Mb) and **bc1b2** (17.69-18.95Mb). We note that the telomeric (left) breakpoints of **Bc1b1** and **bc1b2** are adjacent (17.68 and 17.69Mb); their pairing with the centromeric (right) breakpoints (18.95 and 19.26Mb) cannot be uniquely assigned but this uncertainty does not impact the relationship between **Bc1b1** and **bc1b2**. A small (8kb) segment **bb2** is also annotated. The **Bc1a/Bc1b** segments then determine the ancestor segment **Bc**. Segment **Bb1** is defined in subclone **a5** as a sister of **Bb2** based on adjacent centromeric breakpoints. Segments **Ba-Bd** account for the copy number gain between 17.38 and 21.83Mb in subclone **a1-a3** and **a4-1** besides an intact copy of 4p.

Assuming the copy of intact 4p is preserved in all subclones except **a6**, we can then define the remaining segments as follows: Segments **bc1** and **Bd1a/Bd1b** account for the additional copy-number gains in **a5**. The copy-number gains near 20Mb in **a4-2** and **a4-3** are attributed to segments **bd1a1** and **bd1a2** that are truncated copies of **Bd1a**. Both segments are inferred to be located near the ends of the rearranged chromosome. The copy number gain near 21Mb in **a4-3** is attributed to duplications **bd1b1/bd1b2** that are joined by a foldback junction.

We can further determine the segmental structure of rearranged DNA combining segmental breakpoints and junctions.
For **B** segments, we have

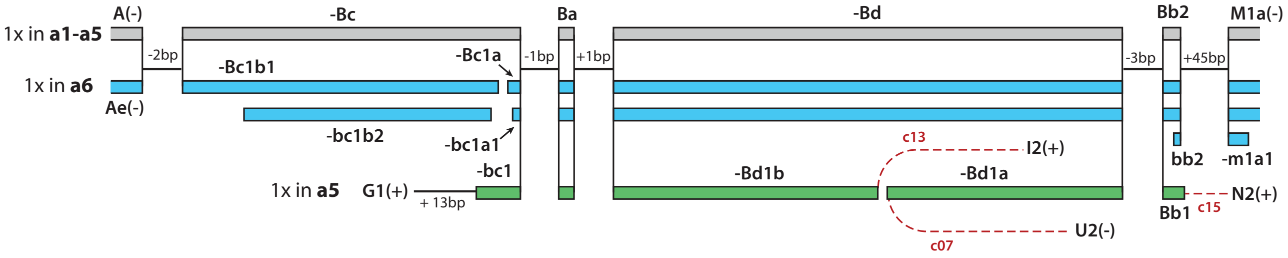


We have labelled microhomology or insertion features at the junctions.
In subclone **a6**, several breakpoints cannot be mapped exactly (the junction sequences suggest locations in the acrocentric arms); the long range linkage between segments **-bc1a1/-Bc1a**, **Bb2/bb2**, and **-M1a/-m1a1** also cannot be determined. One possible arrangement of the rearranged segments is shown below.

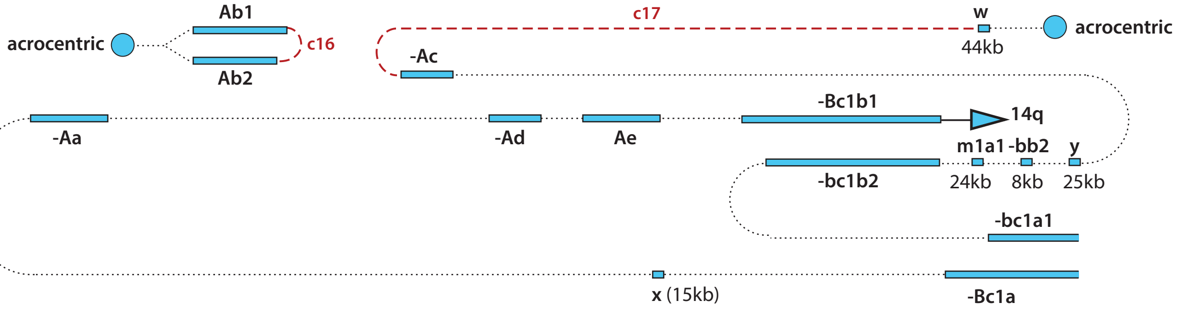

**Copy-number variation in bridge clone a: 17-22Mb**

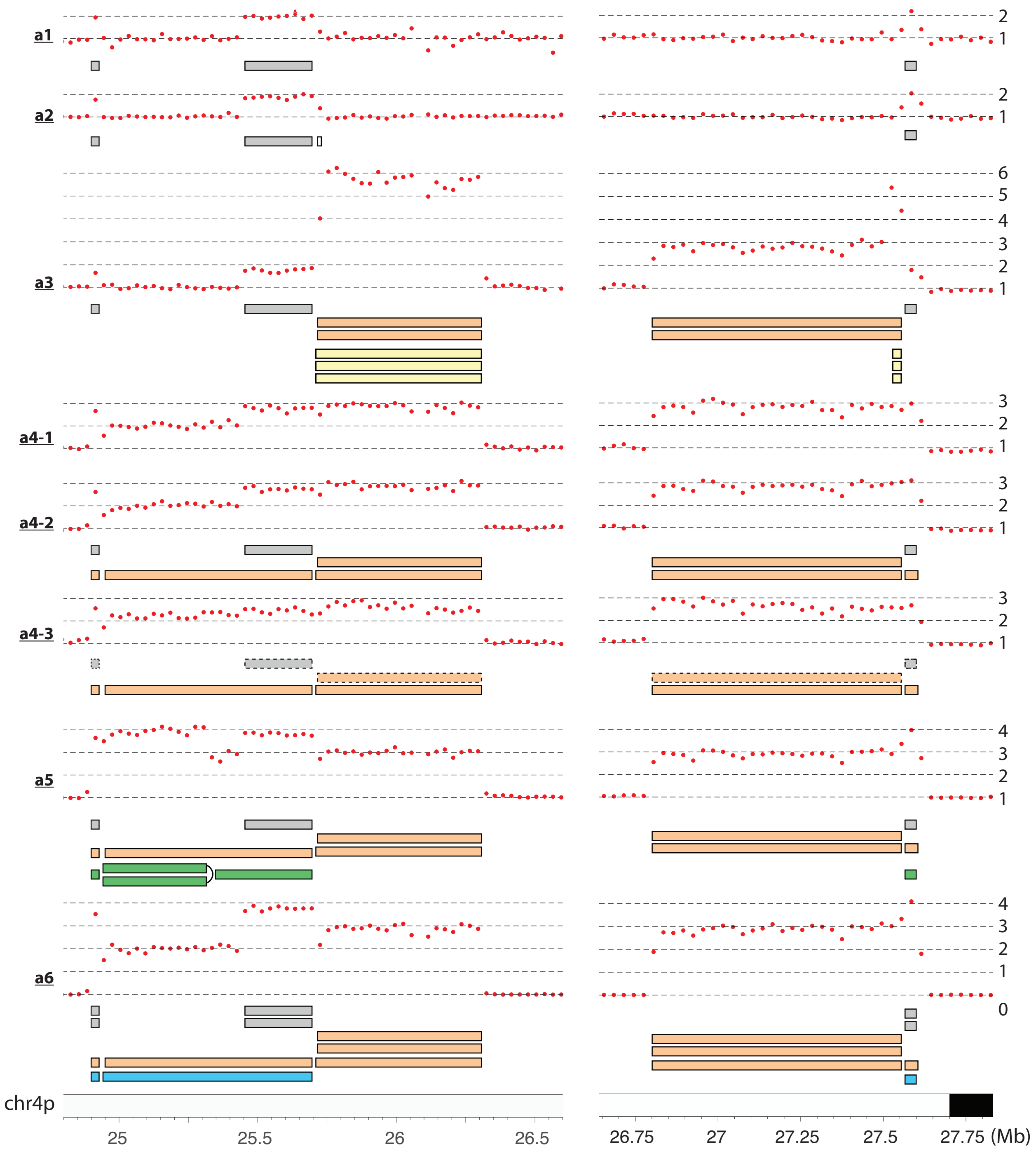

**Segmental structure of copy-number gains: 17-22Mb**

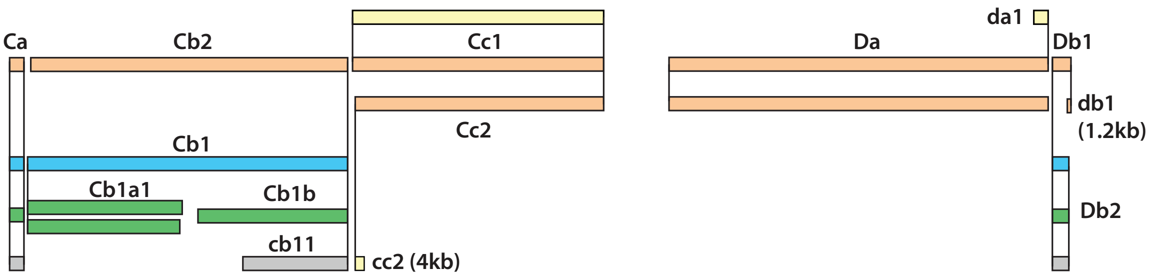


We demonstrate the inference of ancestral DNA structure from these segments. First, we define the following segments based on breakpoints in subclone **a6**: **Ca**(4x), **Cb1**(1x)/**Cb2**(1x)/**cb11**(2x), **Cc1**(1x)/**Cc2**(2x), **Da**(3x), **Db1**(1x)/**Db2**(3x). Numbers in parentheses indicate segmental copy number. From the segmental copy number in different subclones (**A**), we can identify ancestral segments (preserved in more than one subclone) and secondary segments (**B**). From the ancestral segments, we can identify adjacent *cis* and *trans* breakpoints based on breakpoint distance and infer the structure of ancestral DNA ends (**C**).

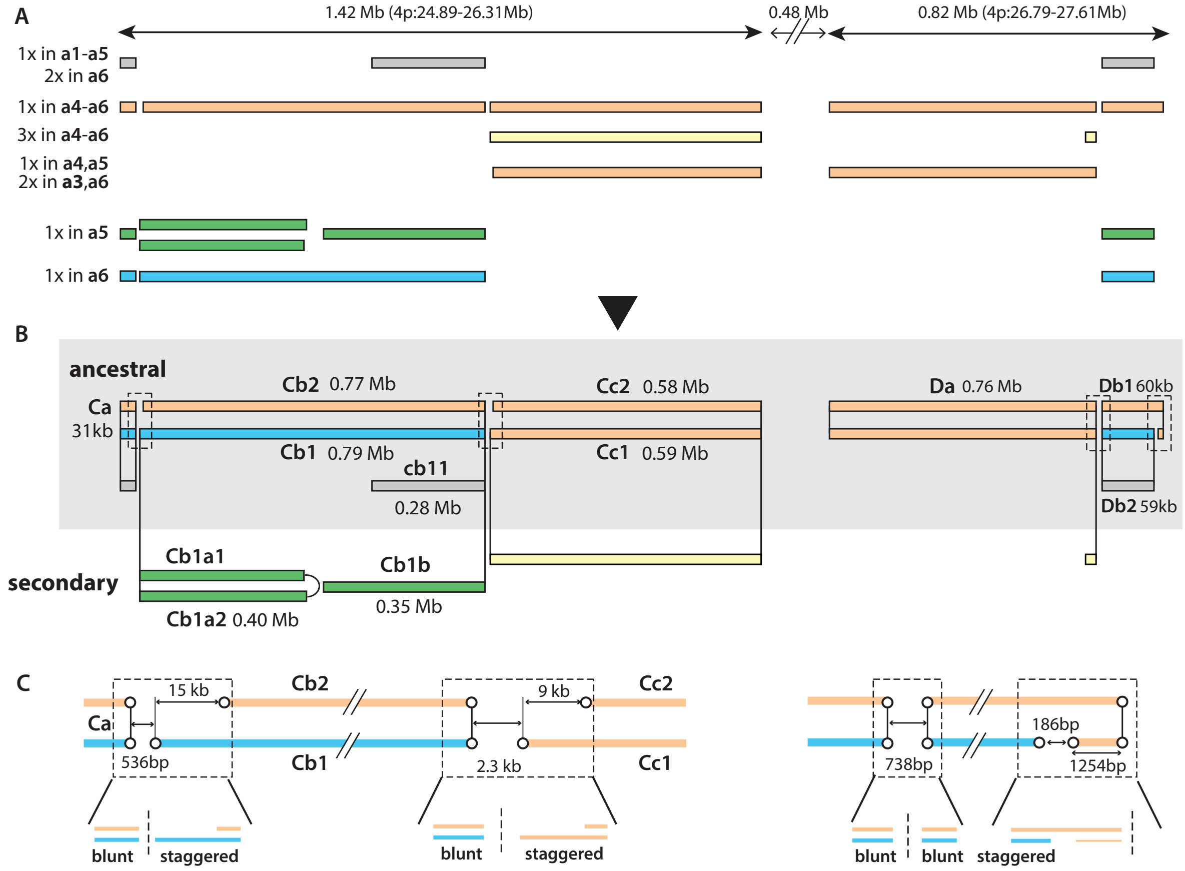


Finally, we can determine the segmental structure of rearranged DNA from the junctions between breakpoints:

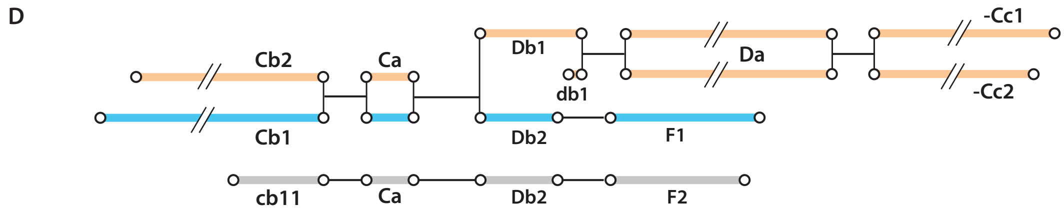


We conclude that there are two copies of **Ca** and **Db2**: one in *cis* with **cb11** and **F2**, the other in *cis* with **Cb1** and **F1**.

#### Copy-number variation in bridge clone a: 30-39Mb

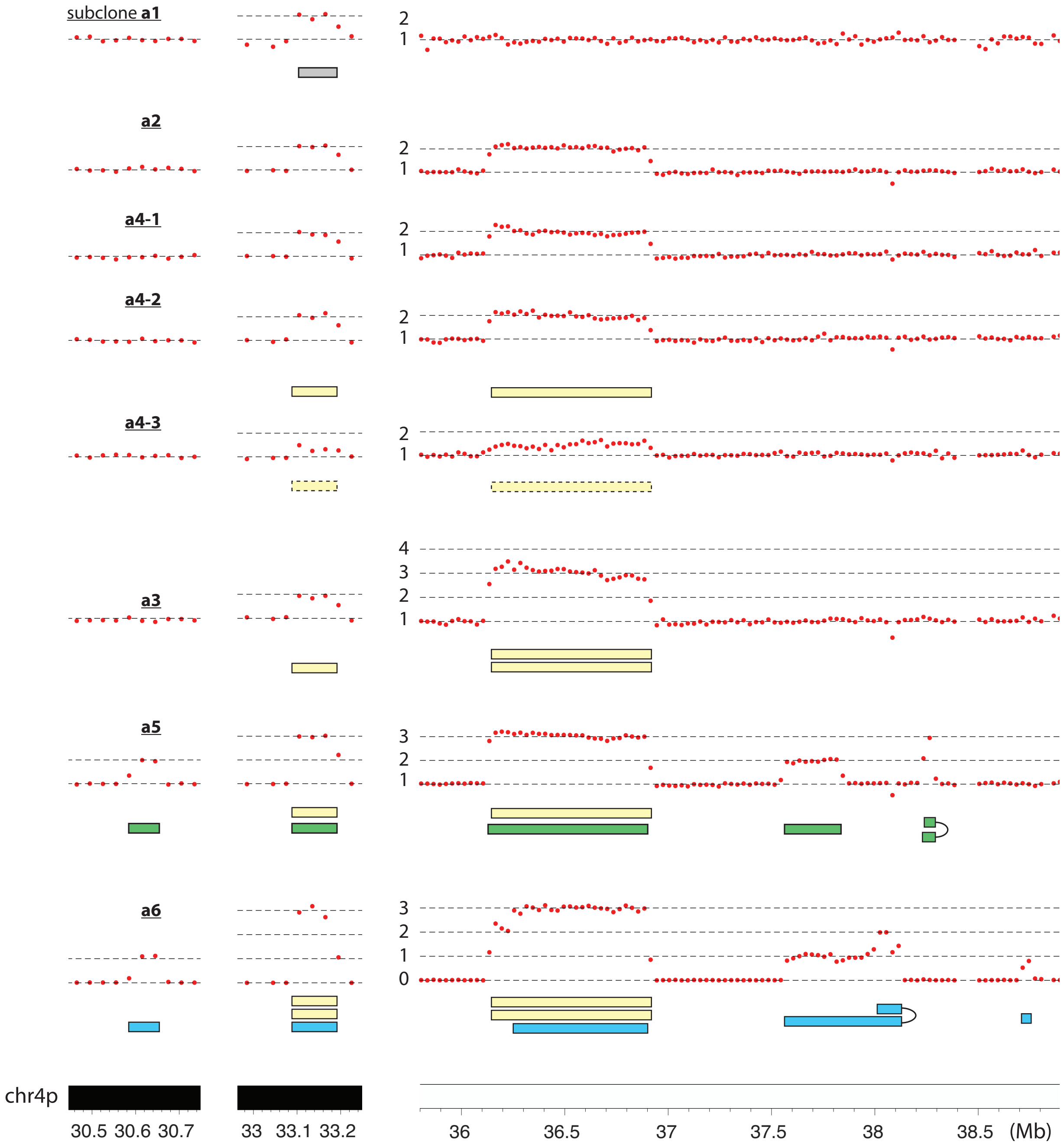

**Segmental structure of copy-number gains: 30-39Mb**

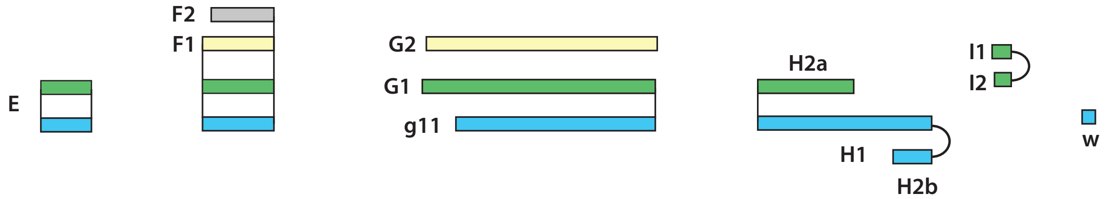


From subclone **a6** we determine the breakpoints of the following segments: **E**(1x), **F1**(3x), **G2**(2x), **g11**(1x), and **H1**(1x)/**H2b**(1x). **H1** and **H2b** are joined together by a foldback junction with insertions. There is also an **a6**-private segmental gain (**w**:38.71-38.75Mb) that could have either *cis* or *trans* breakpoints. This private segmental gain does not impact the analysis of ancestral segments in other subclones. Segment **G1**, with breakpoints identified in subclone **a5**, is determined to be a sister of **G2** based on adjacent breakpoints at both centromeric and telomeric boundaries. Segment **F2** is inferred to be a sister of **F1** based on adjacent breakpoints at the centromeric side (right).

We further determine the two copy gain near 38.2Mb in subclone **a5** to be due to a pair of sister segments **I1/I2** that are joined by a foldback junction. The other segmental gain in **a5** between 37.55 and 37.83Mb is inferred to be a fragment **H2a**: both **H2a** and **H2b** are derived from an ancestral segment (**H2**) that is a sister of **H1**.

The segmental structure of rearranged DNA is as follows (**F** segments will be treated at the end, **w** is included in the rearranged **A** segments in subclone **a6**):

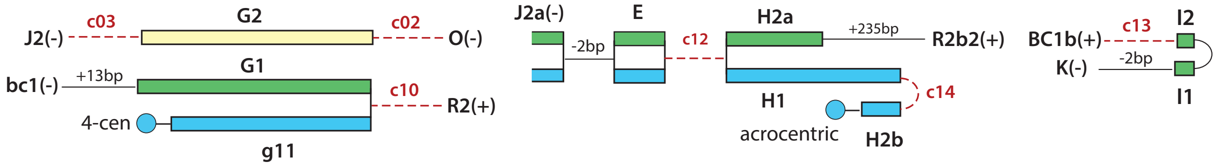

#### Copy-number variation in bridge clone a:  41-46Mb

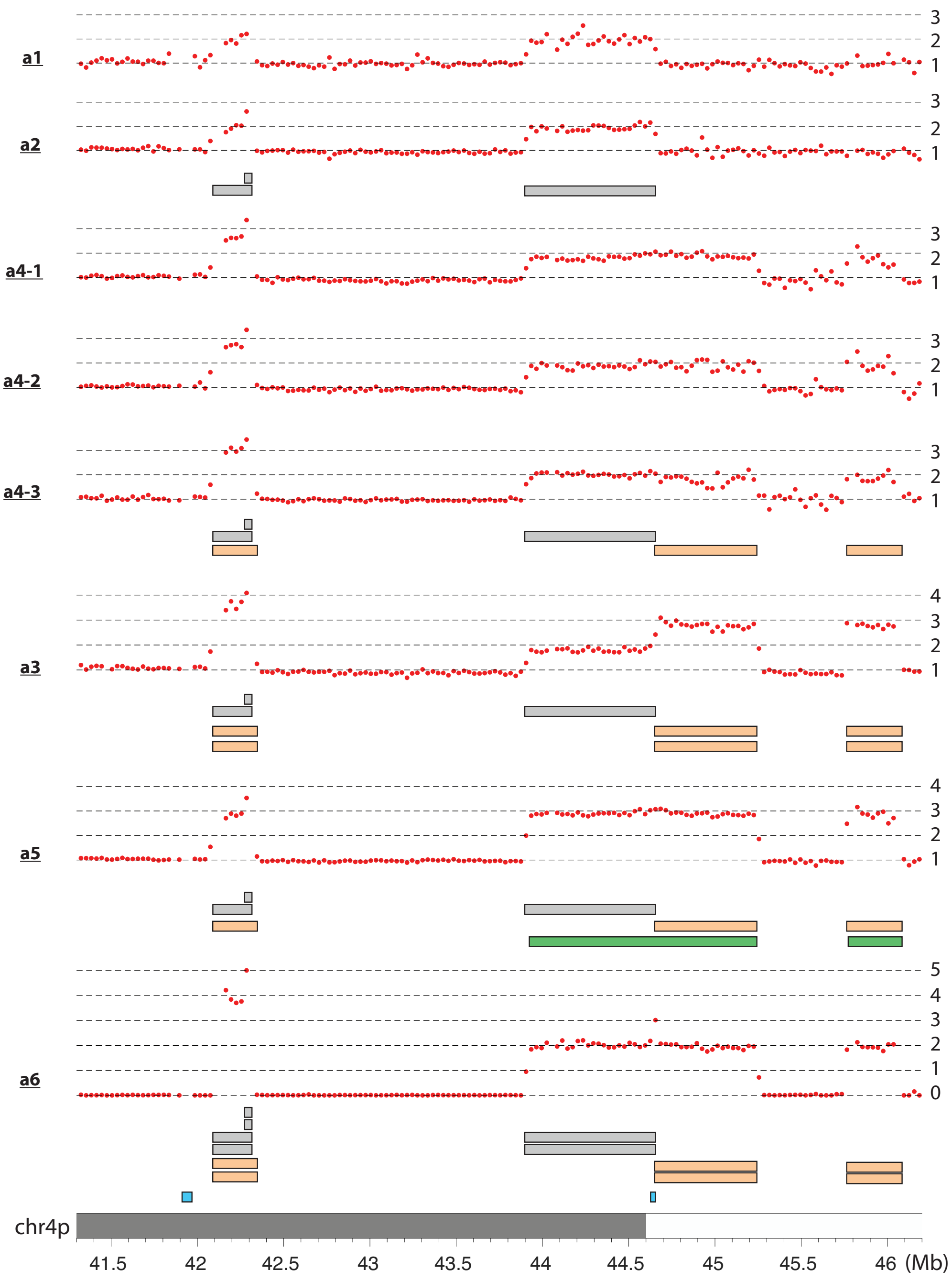

**Segmental structure of copy-number gains: 41-46Mb**

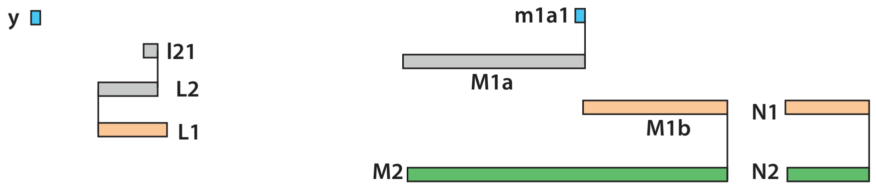


From subclone **a6** we determine the breakpoints of the following segments: **L1/L2/l21**, **M1a/M1b**, and **N1** (all 2x). The **N2** segment is defined in subclone **a5**; **N1/N2** have adjacent breakpoints on the telomeric side. In subclone **a5** we also identify segment **M2** that we infer to be a sister of the ancestor of **M1a/M1b**. We further infer the partial overlap between **M1a** and **M1b** to have been generated by a replication-bypass mechanism (cf. **Extended Data Fig. 3E**).

There are two **a6**-private segmental gains: one (**y**) at 41.91-41.94Mb (25kb), the other (**m1a1**) at 44.64-44.66Mb (24kb). They are included in the rearranged **A** segments in subclone **a6**. The order of the remaining segments in the rearranged chromosome will be presented later.

**Copy-number variation in bridge clone a: 46-47Mb**

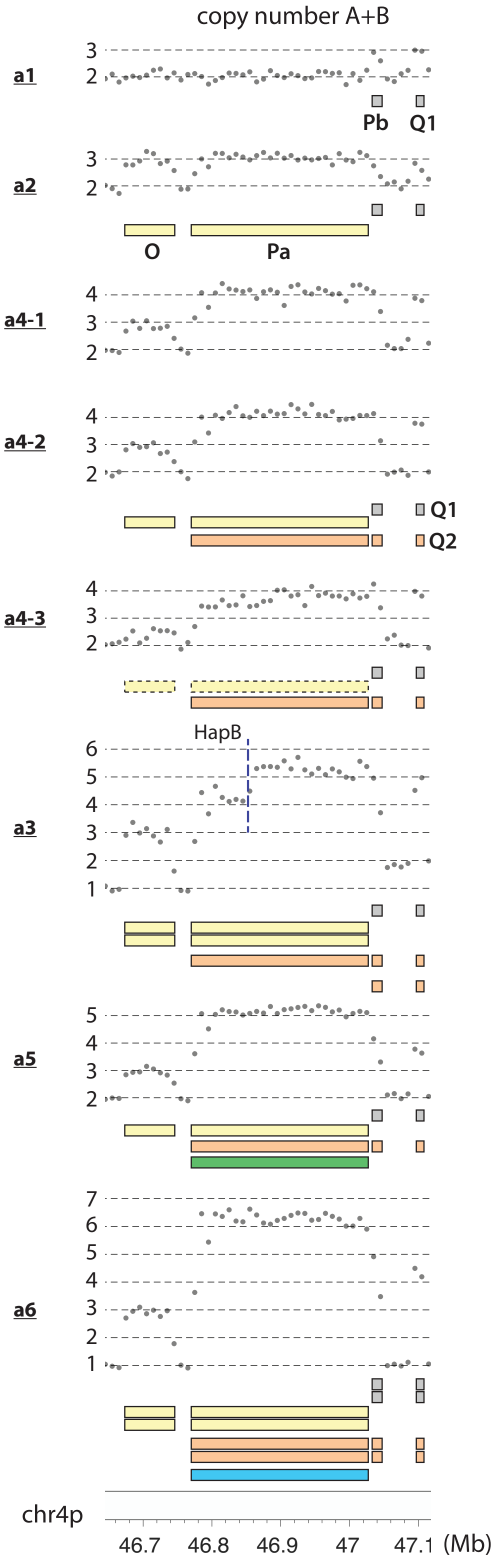


 

**Copy-number variation in bridge clone a: 48-49Mb**

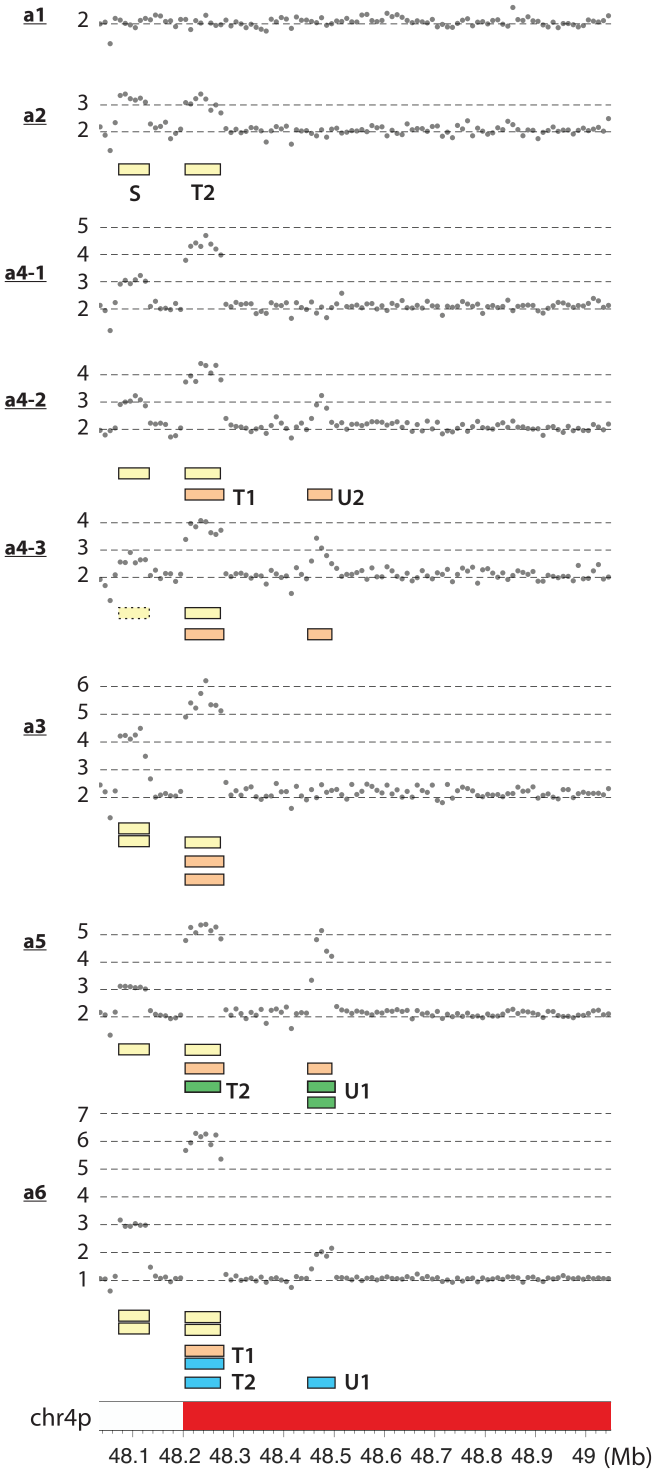

**Segmental structure of copy-number gains: 46-49Mb**

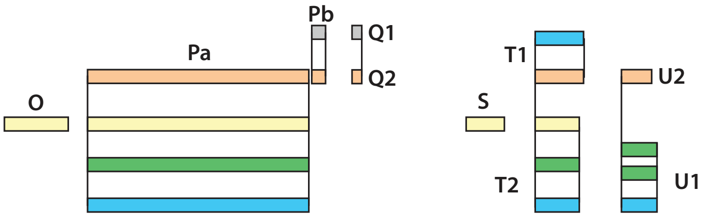


For segmental copy-number variation in 46-49Mb, we use both the total DNA copy number (left) for breakpoint identification and haplotype-specific copy number (right) for breakpoint phasing. We note that subclone **a6** has an intact copy of homolog B. From subclone **a6** we determine the breakpoints of the following segments: **O**(46.67-46.74Mb;2x), **Pa**(46.78-47.029Mb;5x), **Pb**(47.03Mb-47.047Mb;4x), **S**(48.07-48.13Mb;2x), and **U1**(48.46-48.50Mb;1x). We can also determine the breakpoints of **Q1/Q2** (47.09-47.11Mb) and **T1/T2**(48.20-48.28Mb), but their copy-number states need to be determined based on the flanking segments in the rearranged chromosome.

The assembly of segments **L-N** and **Q,Pb,V** and their copy number states in all subclones are shown below.

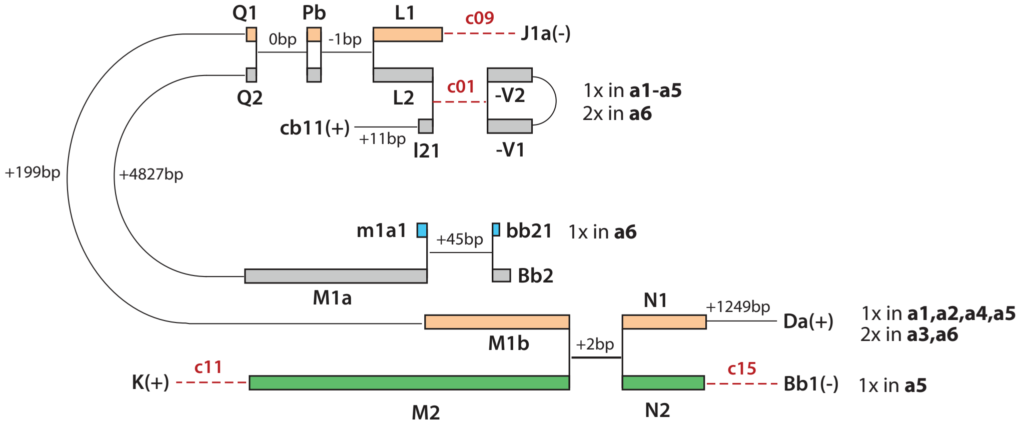

#### Copy-number variation in bridge clone a: 47-48Mb

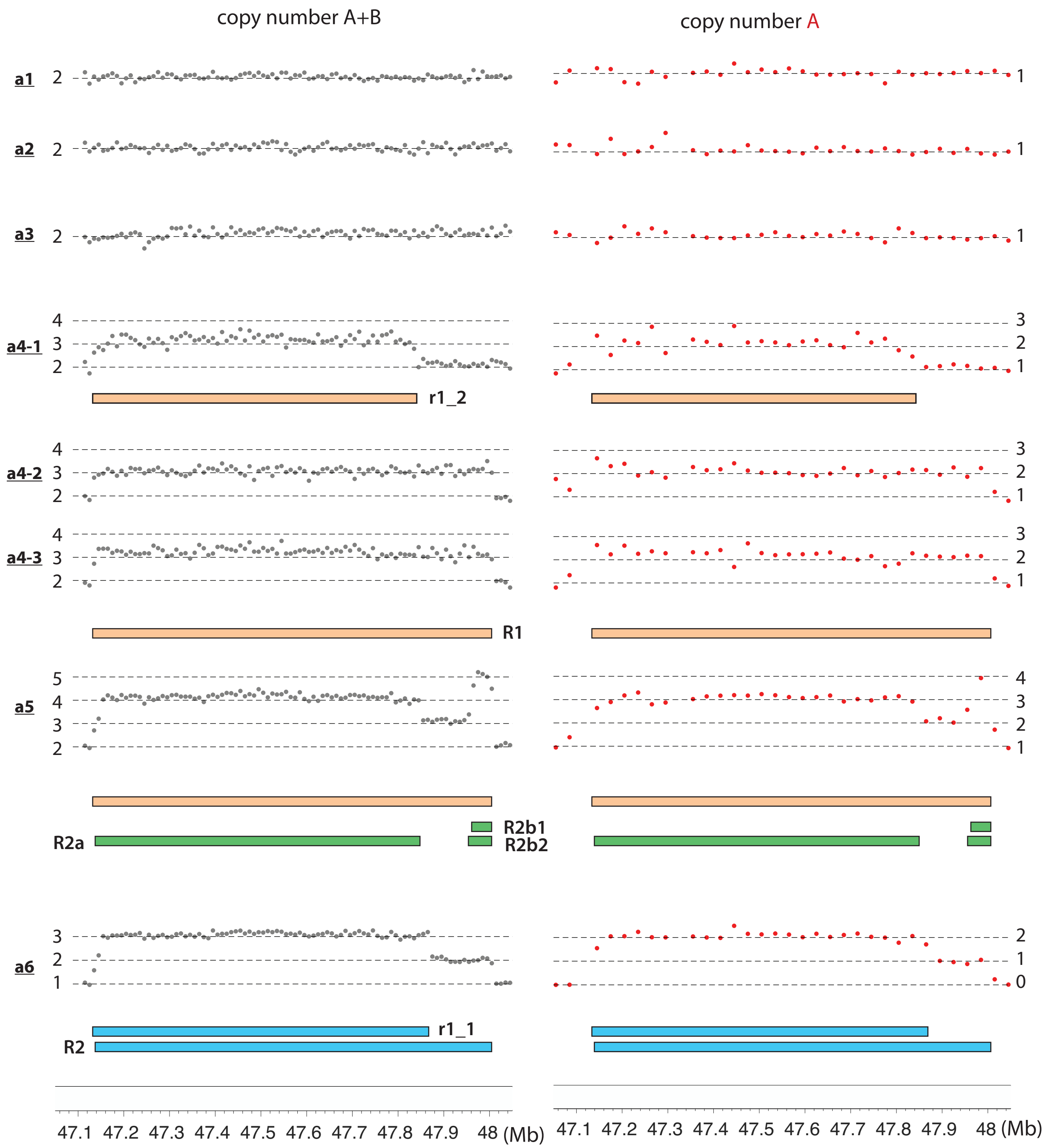

From the segmental copy number in subclone **a4-2** and **a4-3**, we can determine the breakpoints of segments **R1** (47.13-48.01Mb). A truncated segment **r1_2** (47.13-47.84Mb) accounts for the gain in subclone **a4-1**. Based on breakpoint junctions in subclone **a4**, we can assemble the following compound segment: **Cb2(+):R1:U2:Bd1a(-)**. From the segmental copy number and breakpoints in **a5** and **a6**, we infer the presence of a sister segment **R2**: **R1** and **R2** share the same centromeric (right) breakpoint but have distinct, adjacent telomeric (left) breakpoints. The duplications in **a5** and **a6** can have two configurations (#1 and #2 as shown in the right figure). To distinguish between these two configurations, we need to consider the copy number of the flanking segments in the rearranged chromosome.

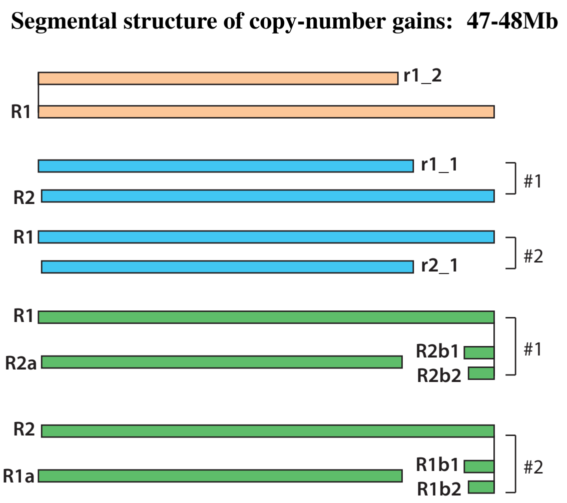


Based on junctions in **a5** and **a6**, we can assemble the following segments flanking **R2**: **G1/g11(-):R2(+)** and **R2(-):U1:Cb1/Cb1a(+)**. The segments flanking **R1** are **Cb2(+):R1(+)** and **R1(-):U2:Bd1a(-)**. The copy-number states of these flanking segments (middle figure) in **a5** and **a6** are determined as follows: **Cb2**(1x in **a5** and **a6**), **U2:-Bd1a**(1x in **a5**; 0x in **a6**), **G1/g11(-)**(1x in **a5** and **a6**), **U1:Cb1(+)**(1x in **a6**; 2x in **a5**). (Note that the copy number states of **U1/U2** are determined by the copy number of the flanking segments **-Bd1a** and **Cb1/Cb1a1/Cb1a2**.) Based on the copy number of the flanking segments, we infer the structure of **R** segments in both **a5** and **a6** to be #1.

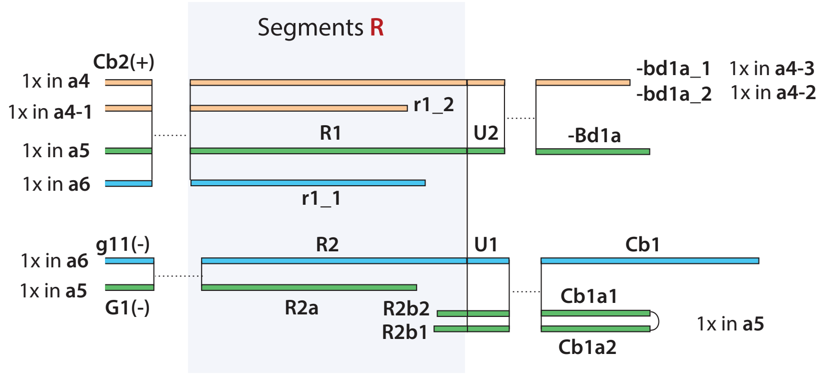


We infer the segments to have been generated by the following sequence of events (bottom figure). (i) End-joining between unreplicated dsDNA fragments, **R1/R2(-)** and **U1/U2(+)**; (ii) incomplete replication creates internal ssDNA ends in **R1/R2**; (iii) fusions of newly generated dsDNA ends: **R1(+)**, **R2(+)**, **U1(-)**, **U2(-)**. Note that the internal ssDNA ends remain unligated. (iv) After another round of replication, the internal ssDNA ends are converted into dsDNA ends, generating segments **R1a**, **R1b**, **R2a**, and **R2b**. **R1b** is lost; **R1a** (final name **r1_1**) and **R2a** undergo further fusions but **R2b** remains unligated. (v) **R2b** is duplicated to create **R2b1/R2b2**; **r1_2** is derived from **R1** or **R1a**. Based on this inference, the adjacent breakpoints of **R2a(-)** and **r1_1(-)** originate from sister DNA ends due to incomplete replication, whereas the adjacency between **R2a(-)** and **r1_2(-)** occurs by chance.

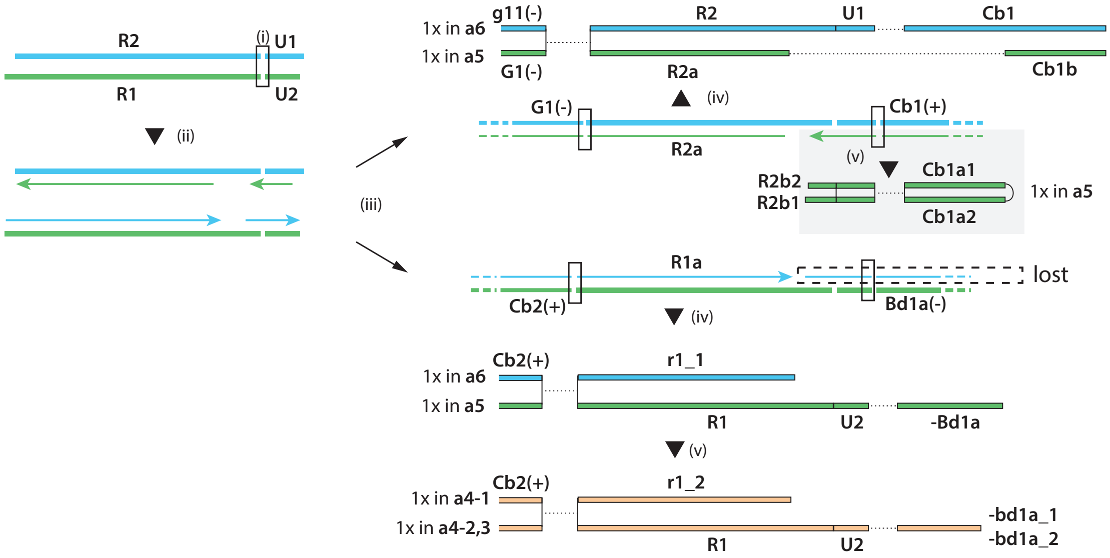

#### Copy-number variation in bridge clone a: 39-41Mb

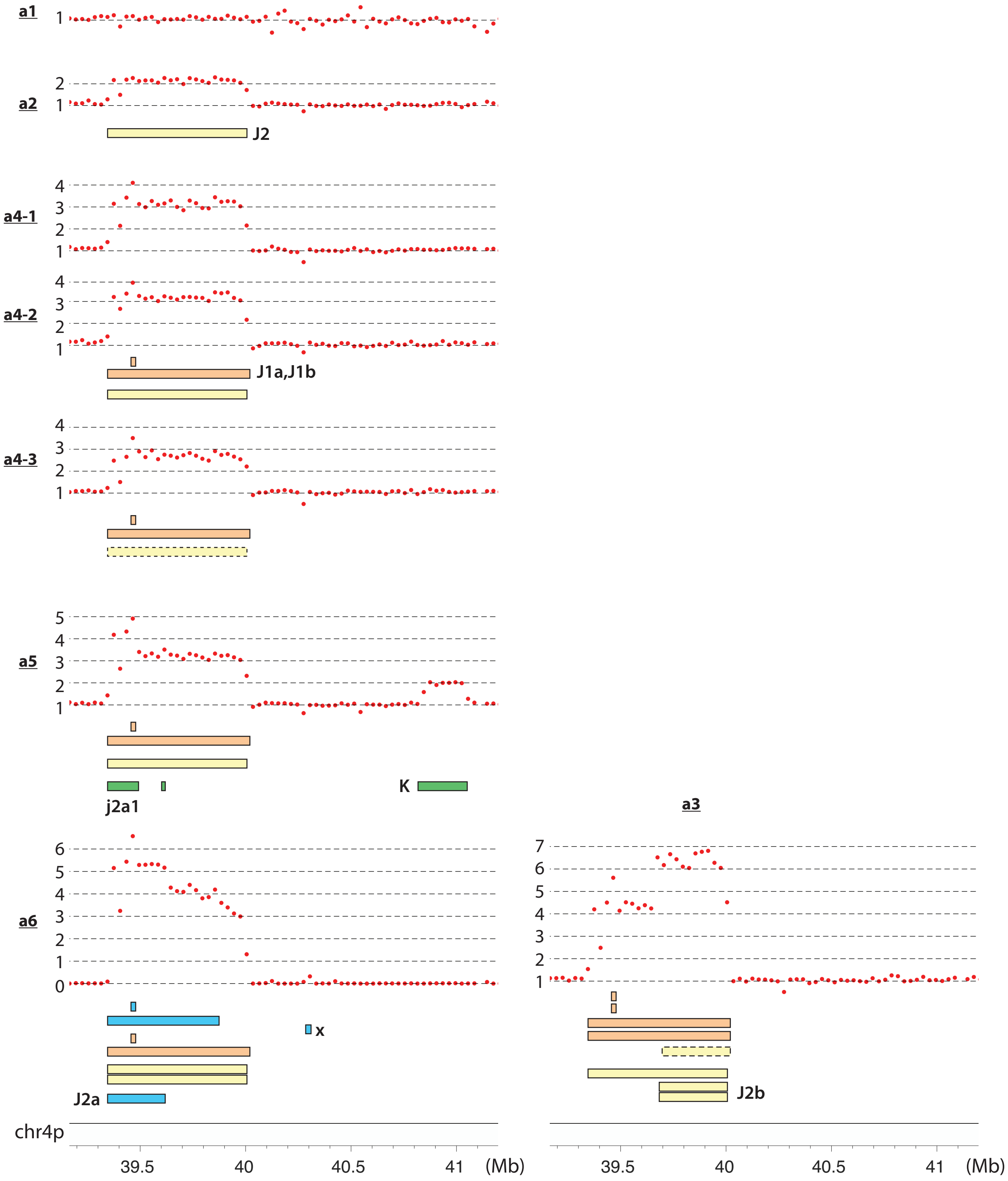


Copy-number gains at 40.83-31.05Mb in subclone **a5** and at 40.29-40.31Mb in subclone **a6** are attributed to segments **K** and **x**. Segment **x** is private to subclone **a6** but segment **K** is inferred to be ancestral as the breakpoints form junctions with segments **I1** and **M2**, both of which inferred to be ancestral. The **J** segments are discussed in the next page.

**Segmental structure of copy-number gains: 39-41Mb**

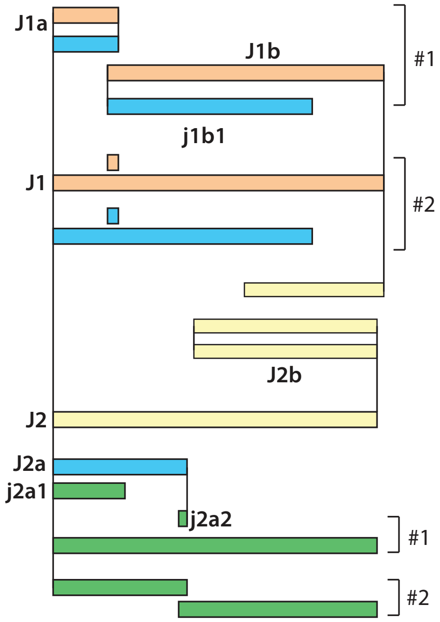


(Left figure) For the amplified **J** segments, the copy-number gains in subclone **a2** and **a4** suggest the presence of two sister segments **J1** and **J2** (39.35-40.01Mb), plus a 23kb triplication at 39.45-39.47Mb. We infer there is a contiguous **J2** segment based on the copy-number gain in subclone **a2**; the extra duplications in **a4** can be realized either with two pieces (**J1a/J1b**) with a 23kb partial overlap (#1), or with a single segment **J1** plus a 23kb segment (#2). We favor the two-piece configuration because of the opposite coordination between breakpoints that we attribute to sister-DNA exchange (cf. **Extended Data Figure 4**). However, the other configuration cannot be definitively excluded.

We further determine the presence of segment **J2a** (39.36-39.63Mb) in subclone **a6** and **j2a1** (39.36-39.48Mb) in subclone **a5**. Although the breakpoint of **j2a1(-)** at 39.482Mb is close to the breakpoint at 39.469Mb (**J1a(-)**), we infer **j2a1(-)** to be a secondary breakpoint because its partner **R2b1(+)** is secondary; by contrast, **J1a(-)** is an ancestral breakpoint. Therefore, we think the adjacency between **J1a(-)** and **j2a1(-)** occurs by chance.

Subclone **a5** further contains a small duplication at 39.622-39.630Mb; this short duplication can be realized either as a fragment of **J2a** (**j2a2**, #1), or two partially overlapping fragments (#2) generated by replication-bypass. Before discussing these two possible configurations, we first analyze the segmental copy number to determine long-range segmental structure of rearranged DNA.

Based on breakpoint junctions, we assemble the following compound segments in the rearranged DNA (right figure):

i (from **a1**, gray) **cb11:Ca:Db2:-F2**;
ii (from **a2**, yellow) **cb11:Ca:Db2:-F1:-T2:Pa:J2**(:G2:-O:S);
iii (from **a4**, orrange) **Cb2:Ca:Db1**(:Da:-Cc1);
iv (from **a4**) (S:Cc2:-Da:N1:-M1b:Q2:Pb:L1:) **-J1a:-Pa:T1:J1b**(:Cc1);
v (from **a6**, blue) **Cb1:Ca:Db2:-F1:-T2:Pa:J2a**(:E:H1);

The bold-faced segments are shown on the right and the flanking segments (in parentheses) are not shown.

We can further infer a plausible evolutionary history of the triplicated segments **cb11** and **J2a** from re-replication. The compound segments suggests five rearranged DNA fragments (i)-(v). From (i) and (ii), we infer the presence of an ancestral dsDNA fragment with the same composition as (ii). (ii) and (v) can result from a re-replication (e.g., due to late-firing origins within **F1**) that creates unpaired segments **cb11** and **J2a**. Finally, the pairing of adjacent breakpoints suggests two ancestral dsDNA fragments with staggered ends generate (iii), (iv), and the ancestor of (ii) and (v). Note the opposite strand coordination between **T1/T2** and **J1/J2** due to sister-chromatid exchange.

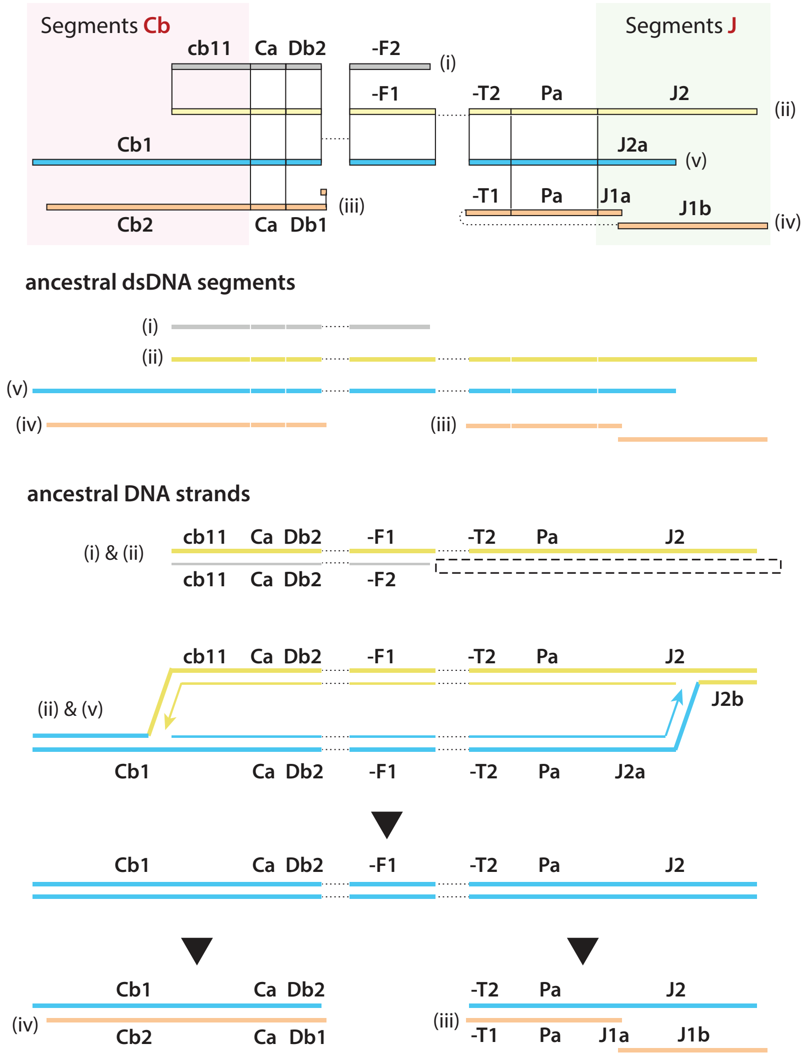


 

Based on the assembled chromosome fragments (ii-v), we can determine the copy number state of each compound segment in each subclone. This analysis can verify the continuity of the assembly and determine the locations of secondary breakpoints. For example, the presence of a single copy of **Cb1** and **Cb2** in subclone **a6** indicates there is a single copy of **J2a** (linked to **Cb1**) and **J1b** (linked to **Cb2**). But the presence of two copies of **S** that are linked to **J1a** indicates a breakpoint between **J1a** and **J1b**. This leads to the inference of **j1b1**. Note that this inference is independent of the configuration of **J1**.

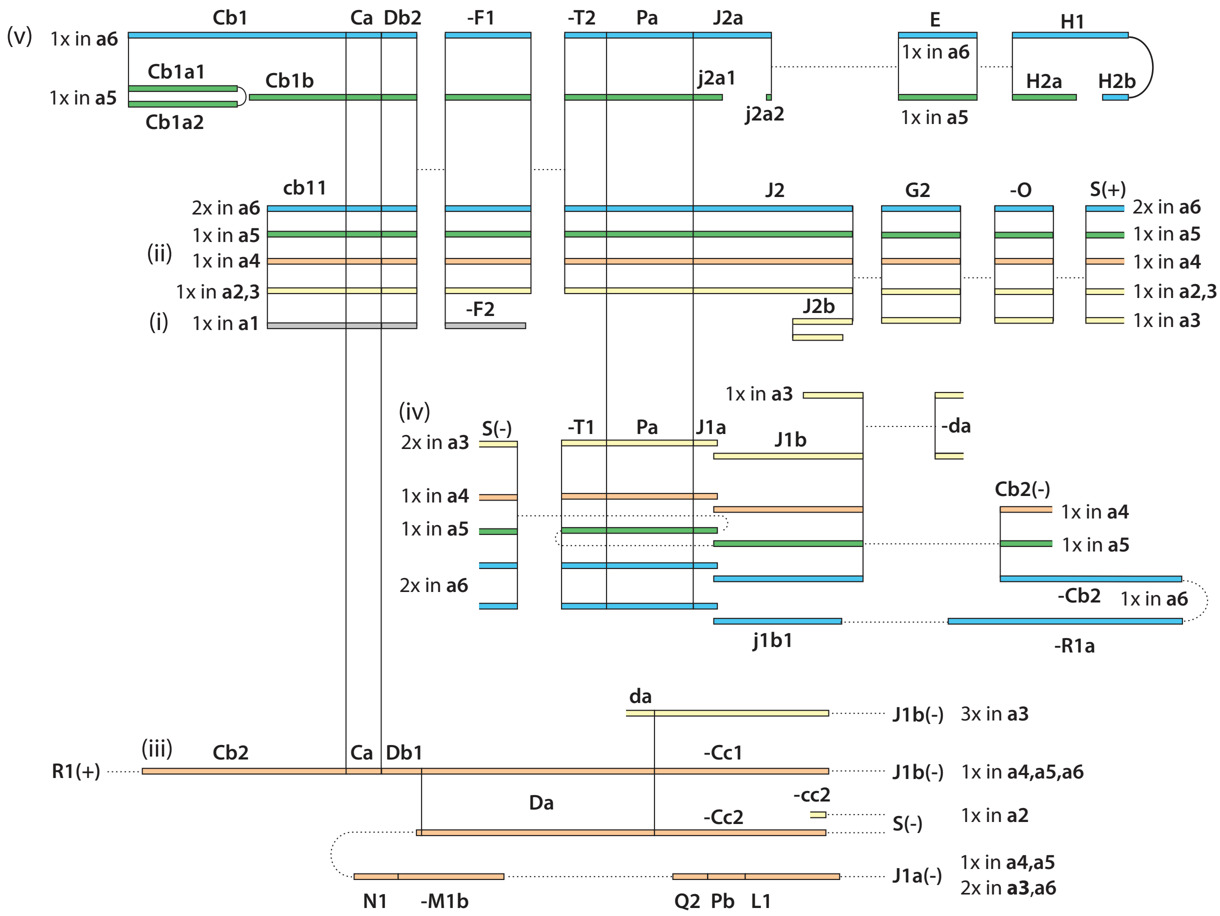


There remains some uncertainty about amplified segments in subclone **a3** and **a5**. In **a3**, there are three extra truncated copies of **J1/J2** and we can only identify one breakpoint at 36.66Mb(+) that is fused with telomeric repeats. A plausible origin of this breakpoint is from the breakage of the re-replication bubble. We therefore named this segment **J2b** (36.66-40.01Mb) as a *cis* segment of **J2a** (39.35-39.62Mb), although we cannot rule out the possibility that they are generated independently. In **a5**, the private breakpoints at 39.48Mb and 39.63Mb can reside on any of **J2a**, **J2**, or **J1b**, as shown on the right. We choose the first configuration in the schematic diagram for convenience. In either configuration, the breakpoint at 39.63Mb with newly added telomere repeats represents the end of the rearranged chromosome.

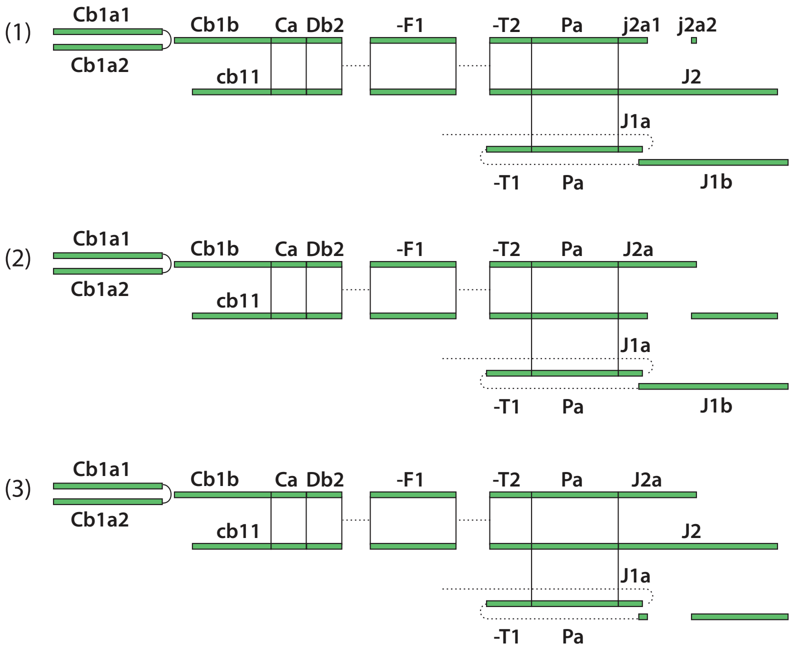

#### 4. Segmental structure of rearranged chromosomes in different subclones of bridge clone a

We can determine the order of segments in the rearranged chromosomes in subclone **a1**, **a2**, and **a4** by concatenating segments and junctions. The rearranged chromosomes are broken into multiple fragments (**1-5**) each consisting of multiple segments. Segments in each fragment are colored based on the subclone where they are present at the lowest copy-number state (usually single copy): **a1** (gray); **a2** (yellow); **a4** (orange). Open and filled numbers denote linkage between two fragments.

**Segmental structure of rearranged chr4 in a1; a2; a4**

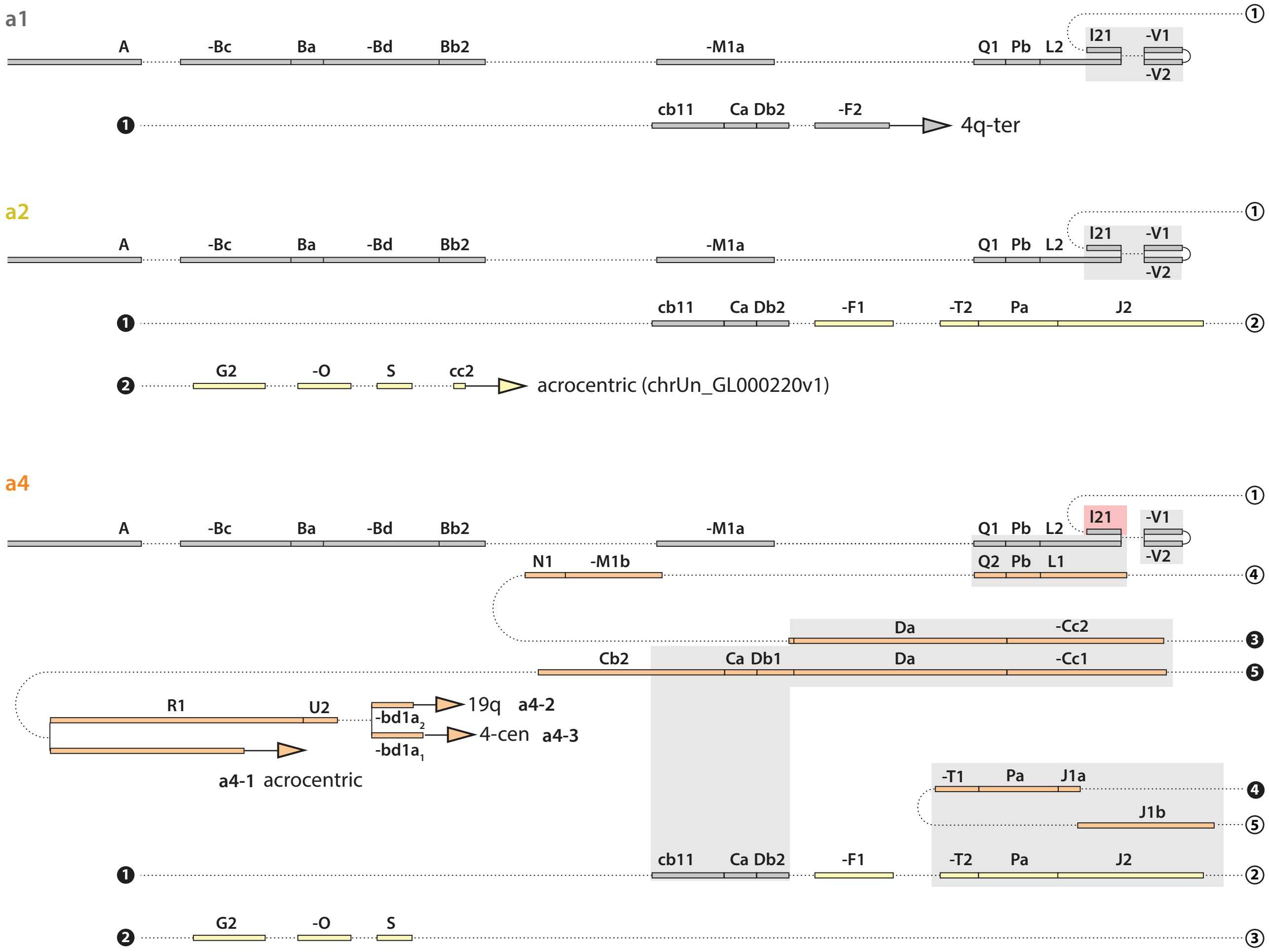


Duplicated segments are highlighted in gray rectangular areas. A triplicated segment (**I21**) is highlighted in a red rectangular area. The following duplications have adjacent parallel breakpoints and are inferred to originate from sister DNA segments generated by replication:**V1/V2**, **Q1/Q2**, **L1/L2**, **Cc1/Cc2**, **Db1/Db2**, **T1/T2**, and **J1/J2**. As sister duplications are generated concurrently, if one of them is inferred to be ancestral, the other also has to be ancestral. Therefore, the absence of one sister segment in a subclone (e.g., **Q2** is not present in **a1**) indicates a secondary loss. We further infer that the same process generates duplicated copies of **Ca**, **Da**, **Pa** and **Pb**. These segments are sandwiched between segments with adjacent parallel breakpoints, i.e., compound dsDNA fragments with staggered ends (e.g., **Q1:Pb:L2/Q2:Pb:L1**). The only exceptions are **cb11** and **I21**, both of which are partial duplicates of **Cb1** and **L2**.

The end of the rearranged chr4 in subclone **a1** is capped by a telomeric segment from 4q (gray triangle). The end of rearranged chr4 in subclone **a2** is capped by rDNA (`chrUn_GL000220v1`). The end of rearranged chr4 in **a4-1** is capped by a sequence that maps to acrocentric arms; the end of rearranged chr4 in **a4-2** is capped by a telomeric segment from 19q; the end of rearranged chr4 in **a4-3** is capped by a sequence that maps to the chr4 centromere. The approximate locations of the repeats are determined by alignment to the `CHM13` reference using `BLAT` from the UCSC genome browser.

Based on the structure of rearranged chromosomes in **a2** and **a4**, we can partially determine the structure of amplified DNA in subclone **a3** based on the copy number.

**Segmental structure of rearranged chr4 in a3**

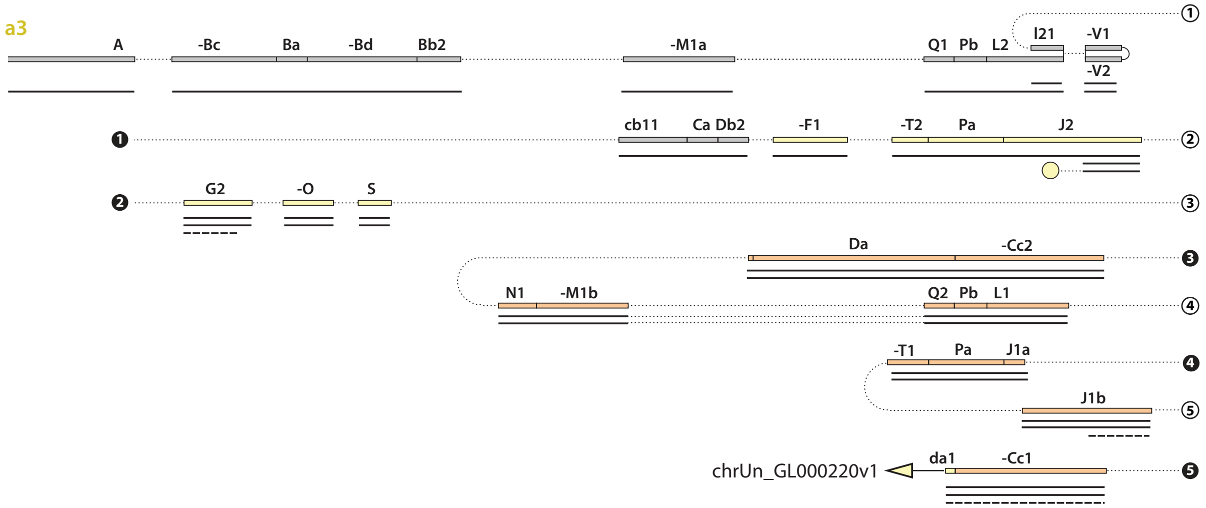


Here each solid line represents a single copy of the chromosome fragment and dashed lines represent fragments that are partially lost during subclonal expansion. Copy-number transitions indicate secondary breakpoints. The yellow circle in segment **J2** denotes fusion to telomeric repeats; the yellow triangle next to segment **da1** denotes fusions to rDNA repeats.

Subclone **a5** and **a6** contain additional duplications including both ancestral segments and secondary duplications of ancestral segments. We assemble fragments **6-8** in addition to fragments **1-5** that are preserved in subclone **a1,a2** and **a4**. Fragment **6** is only preserved in subclone **a5**; fragments **7** and **8** are assembled from junctions preserved in both **a5** and **a6** (bottom box): **G1:R2/R2a**, **U1:Cb1/Cb1a1/Cb1a2**, **J2a/j2a2:E:H1/H2a**.

**Segmental structure of rearranged chr4 in a5 and a6**

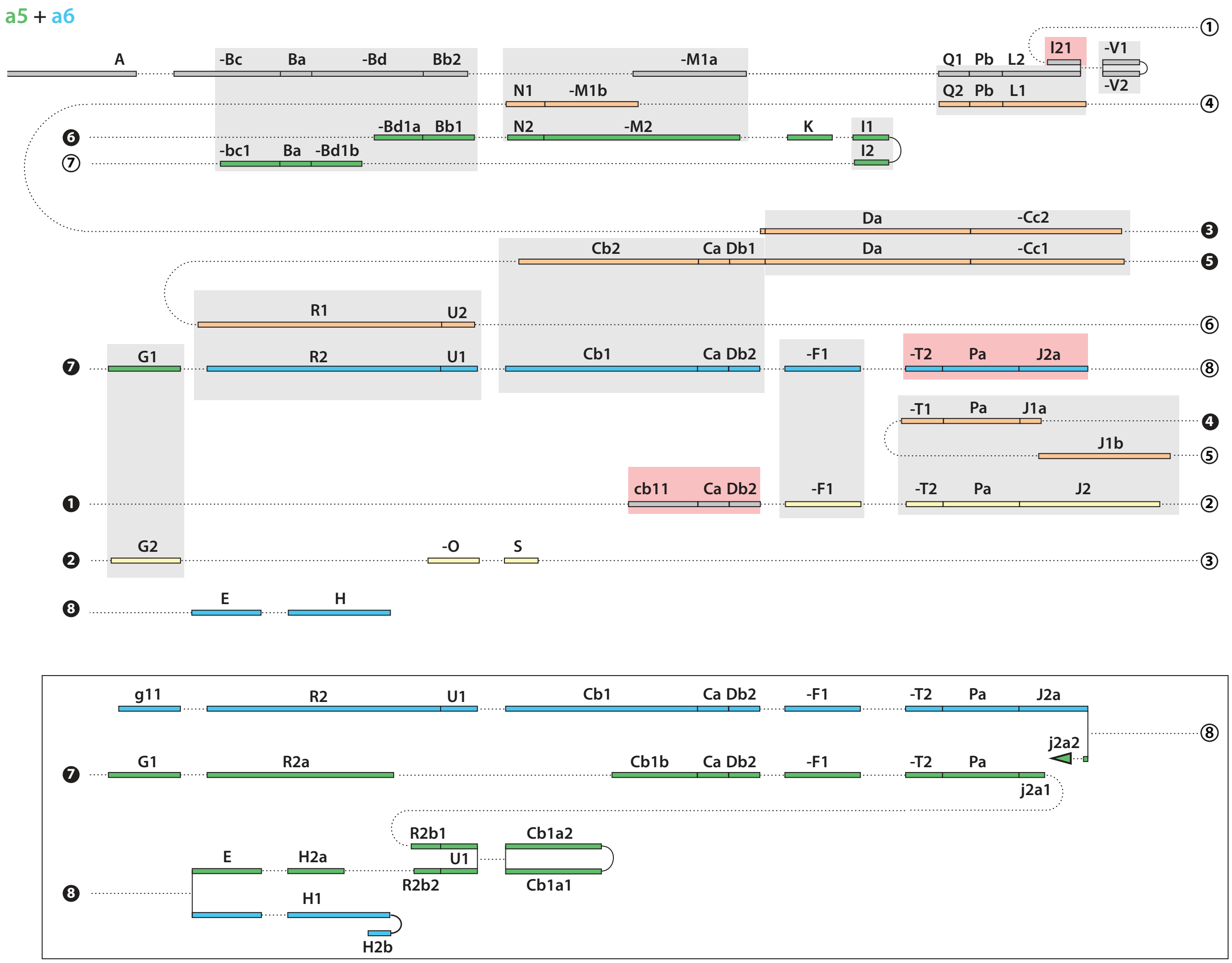


Duplicated segments are highlighted in gray rectangular areas. Triplicated segments are highlighted in red rectangular areas. The end of the rearranged chr4 in subclone **a5** is capped by telomeric repeats (green triangle next to **j2a2**). The structure of rearranged chr4 in subclone **a6** is not completely resolved.

From the fragments (**1-8**) of rearranged chromosomes determined from all the subclones, we can infer the structure of the *ancestral* DNA as shown here.

**Segmental structure of the ancestral chr4**

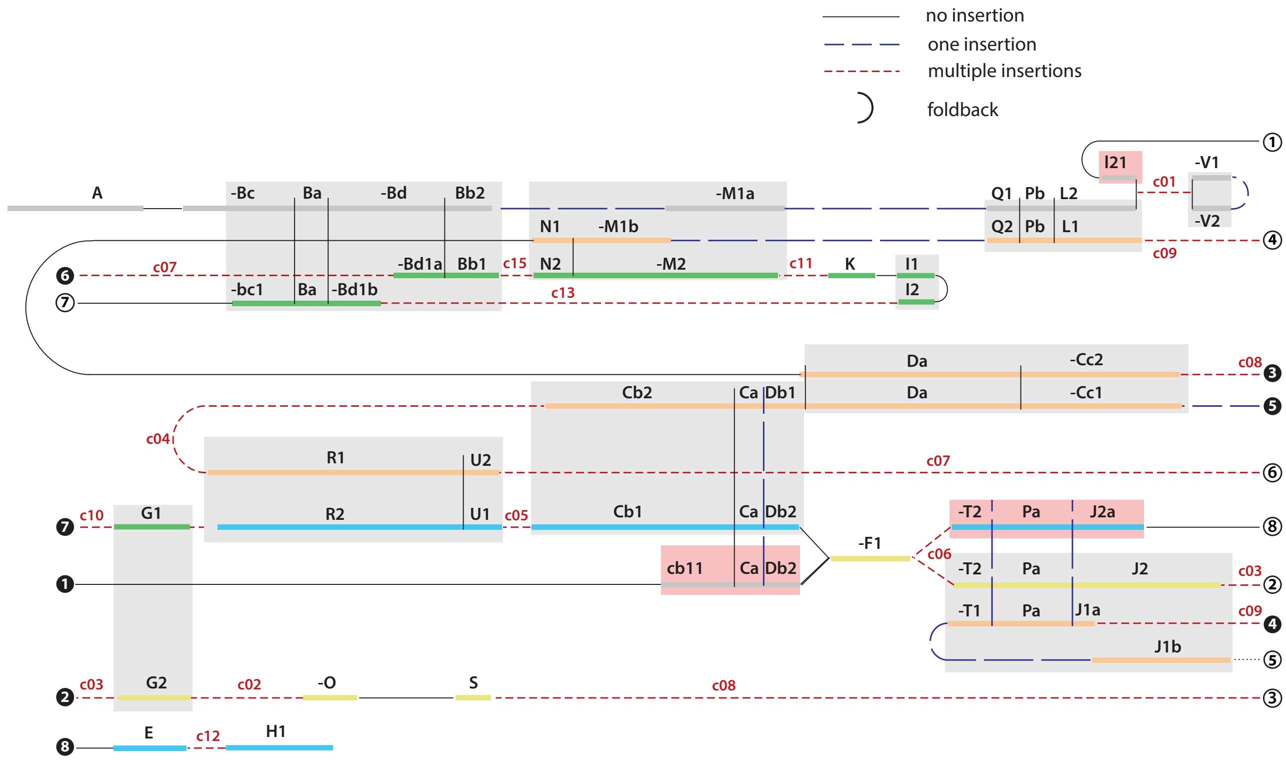


The ancestral DNA segments are colored by the same scheme as above. Features of the junctions between segments are represented as follows. Solid lines: no insertion or short insertions that cannot be mapped; long dashed lines: one insertion; red short dashed lines: complex junctions with more than one insertion. Complex junctions are annotated (**c01-c13**,**c15**). Complex junction **c14** is between **H1** and **H2b** and is likely generated in the next generation.

Based on the structure of the ancestral DNA, we can draw the following conclusions. (1) The following segments are not duplicated: **A**,**E**,**H**,**K**,**O**,**S**. (We infer segments **H2a/H2b** to be secondary duplications since the right breakpoint of **H2a** joins secondary breakpoints on **R2b2**.) (2) The following segments are triplicated (red rectangular areas): **cb11, I21, J2a**. (3) Except for **bc1** as a partial duplication of **Bc**, all the other duplications are sister DNA segments generated by replication (gray rectangular areas). (4) Foldback junctions between **V1** and **V2** and between **I1** and **I2** are formed between DNA ends that are not chromosome ends.

The rearranged chromosome in subclone **a6** is consistent with a duplication of the ancestral chromosome followed by chromothripsis. The DNA copy number suggests that fragments **2-5** are nearly completely duplicated. A comparison between **a5** and **a6** suggests an uneven segregation of duplicated fragments **7** and **8** (bottom box). This uneven segregation most likely occurred in the next generation after the formation of the ancestral rearranged chromosome.

**Segmental structure of rearranged chr4 in a6**

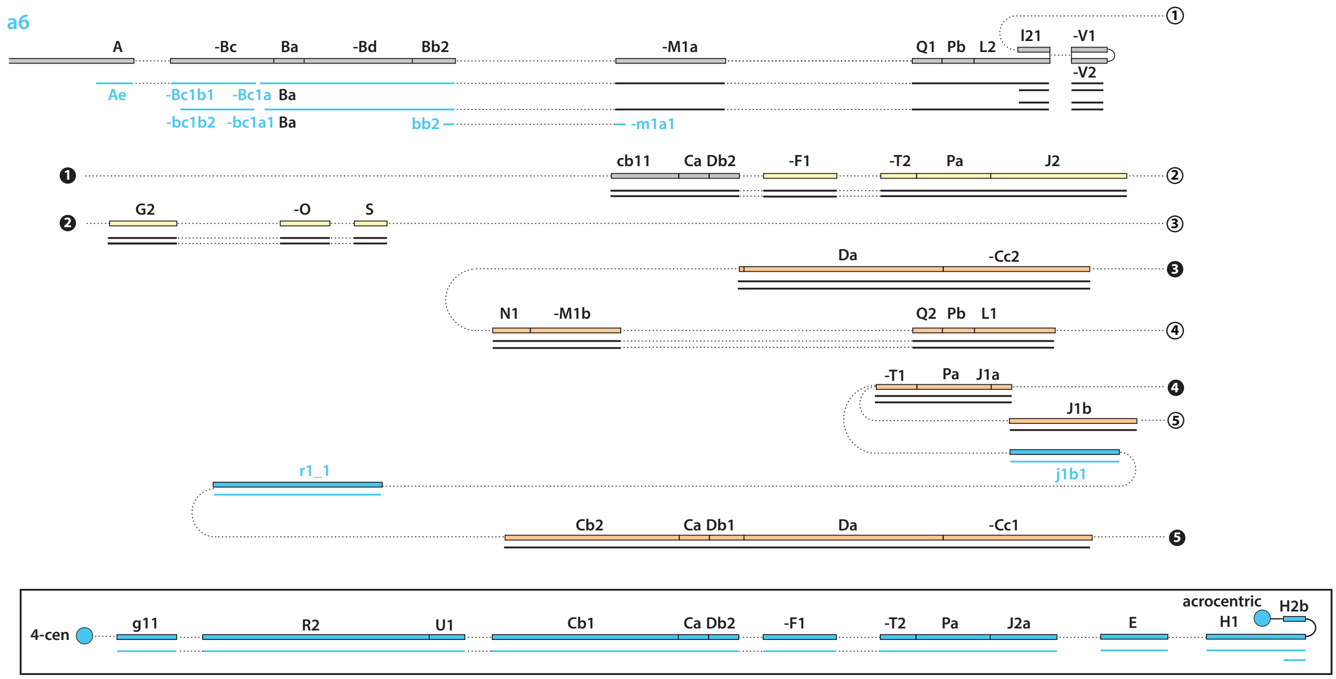

The **a6**-specific segments including **Aa-Ae** are generated in a secondary chromothripsis that also creates complex segmental gains in chr14q (right panels on the bottom figure). The concurrence of these events is supported by two observations. First, there is a junction between segment **Bc1b1** and a breakpoint on chr14:20049947(+). Second, the junction between **j1b1** and **r1_1** contains a short insertion that is mapped to a region on chr14 (20049701-938) right next to this breakpoint.

**Secondary chromothripsis in a6**

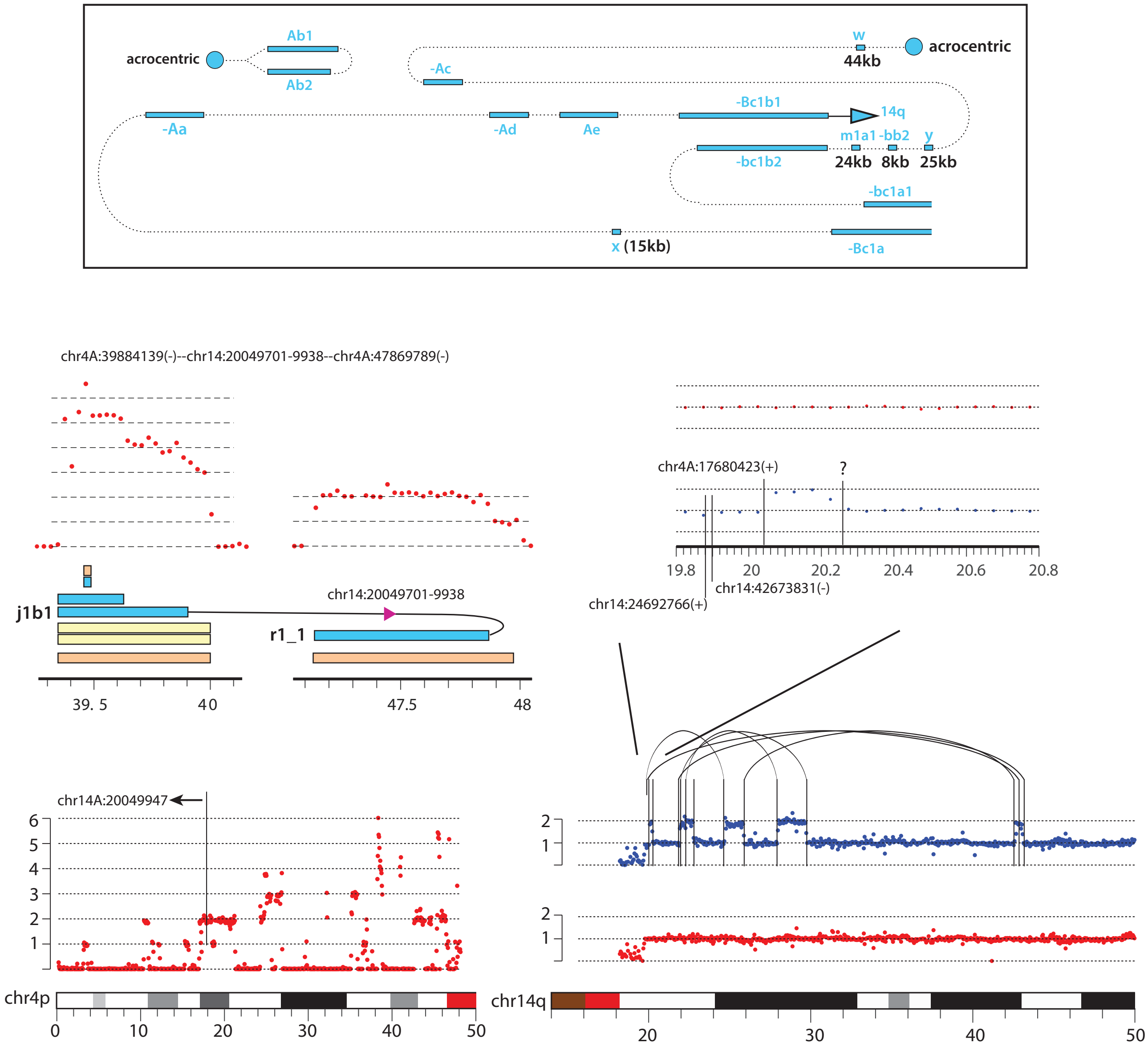

#### 5. Translocations of the ends of rearranged chromosome 4 in bridge clones and subclones

**Translocations between chr4 break ends and other chromosomes**

| Primary clone | subclone | Homolog A | Homolog B |
|---|---|---|---|
| PC1a (F11) | a6 | **10.54 Mb** (+) — acrocentric<br>**17.68 Mb** (+) — **chr14:20.05 Mb** (+)<br>**36.24 Mb** (+) — ? (4-cen)<br>**38.00 Mb** (+) — acrocentric<br>**38.76 Mb** (-) — acrocentric | intact |
| | a5 | **39.62 Mb** (+) — telomere | **44.64 Mb** (+) —acrocentric (13p,14p,21p) or chr9 |
| | a4-2 | **19.94 Mb** (+) — **chr19:41.87 Mb** (+) | |
| | a2 | **25.73 Mb** (-) — acrocentric | |
| | a4-1 | **47.84 Mb** (-)— acrocentric | **44.54 Mb** (+) — acrocentric (13p/21p) |
| | a4-3 | **19.88 Mb** (+) — **49.17 Mb** (+);<br>(subclonal) 20.80 Mb (+) — ? | |
| | a3 | **39.66 Mb** (+) — telomere (3x)<br>**27.51 Mb** (+) — acrocentric (3x)<br>39.95 Mb (-) — ? | **46.86 Mb** (+) — acrocentric |
| | a1-s2,7,8,11,12,16 | **33.10 Mb** (+) — **164.21 Mb** (+) | variable (unmapped) |
| | a1-s13,14 | | **5.82 Mb** (-)—**129.61 Mb** (+) |
| | a1-s18 | | **47.08 Mb** (-)—**185.14 Mb** (+) |
| PC2b (F3) | | **77,011** (+)—**138,703** (+) plus not mapped | variable/not mapped |
| PC1b (F2) | | **36.64 Mb** (+) — **149.25 Mb** (+) | **47.88 Mb (+)** foldback, end not mapped |
| PC2a (F9) | | **35.16 Mb** (-) — acrocentric | p-arm loss/centromeric |
| PC3a/3b(E5/E8) | | mostly lost (not mapped) | intact |
| PC4a (K2) | | **176.87 Mb** (-) — acrocentric | p-arm loss/centromeric |
| PC4b (K11) | | **185.07 Mb** (-) — **chr6:94.91 Mb** (+) | intact |
| PC5a (O16) | | **1.12 Mb** (+), **23.73 Mb** (+), **26.33 Mb** (+), , **31.59 Mb** (-) —acrocentric (14p);<br>**18.39 Mb** (+), **18.42 Mb** (-) — 4-cen;<br>**37.01 Mb** (+) — 7-cen/4-cen | **188.50 Mb** (-) — 4p-tel |
| PC6a (N11) | | **188.88 Mb** (-) foldback, end not mapped | intact |
| PC6b (N6) | | **188.70 Mb** (-) — 4p-tel. | intact |
| PC7a (B4) | | **181.74** Mb (-) — ? (4-cen)<br>**181.06 Mb** (+) — ? (chrUn_KI270746v1)<br>**183.86 Mb** (+) — acrocentric | p-arm loss |

This table summarizes breakpoints on chr4 that form junctions with other chromosomes. The primary clone IDs are consistent with names in Umbreit et al. (2020). Note the prevalence of junctions with repeat sequences from the acrocentric arms, centromeric repeats, or other unmappable repeats. The probable locations of the repeats are inferred by aligning the junction/split sequences to the `CHM13` reference by `BLAT`. Also note the rare presence of foldback junctions at the ends of chromosomes.

We are able to identify several translocations at the ends of rearranged chr4 (**SI Figure 7** on next page). **A**. De novo telomere addition is identified in two subclones: In **a5**, telomeric repeats are inferred to be added to the end of the rearranged chromosome; in **a3**, telomeric repeats are inferred to be added to a duplicated end, so may be interstitial. **B**. Other repeats identified at the ends of broken chromosome 4. **C**. *Top*: An interchromosomal translocation that caps the broken end of chr4 in subclone **a4-2**. A short sequence mapped to a locus (19942536-3070) near the chr4 end at 19940488(+) is inserted at the junction between chr4 and chr19. *Bottom*: Another example of interchromosomal translocation that caps the broken end of chr4 in the PC4b bridge clone. In both examples, the translocated segments from chr19 (∼17Mb) and chr6 (∼75Mb) are beyond the range of break-induced replication; these translocations are most likely generated by fusions between the broken chr4 ends and a replicated DNA fragment that could have been initiated by strand invasion. See Rearrangement outcomes of break-induced replication. **D**. Three examples of intrachromosomal rearrangements that cap the broken ends on 4p by a 4q-terminal segment. In the first example, the broken end on the 4A homolog joins a 4q terminal segment, with a breakpoint located within a pre-existing duplication; the junction sequence (in bold) contains an insertion that is partially mapped to the locus adjacent to the 4q breakpoint. Note the presence of 5-10bp short sequences (**1**,**2**,**1'**) that can be mapped to sequences near the breakpoints. These features could reflect template-switching events after strand invasion; however, the duplicated 4q segment (~30Mb) has to involve conventional DNA replication as the examples shown in **C**. In the second example, the broken chr4B homolog has a large inverted duplication on the 4p terminus joining a 4q segment on the same homolog. In the last example, the broken 4B homolog has a short inverted duplication joining a large 4q terminal segment on the 4A homolog. The last two examples are identified in subclones that have the same copy number of homolog A (**a1**) and inferred to be secondary events.

**SI Figure 7: Examples of translocations of chr4 break ends**

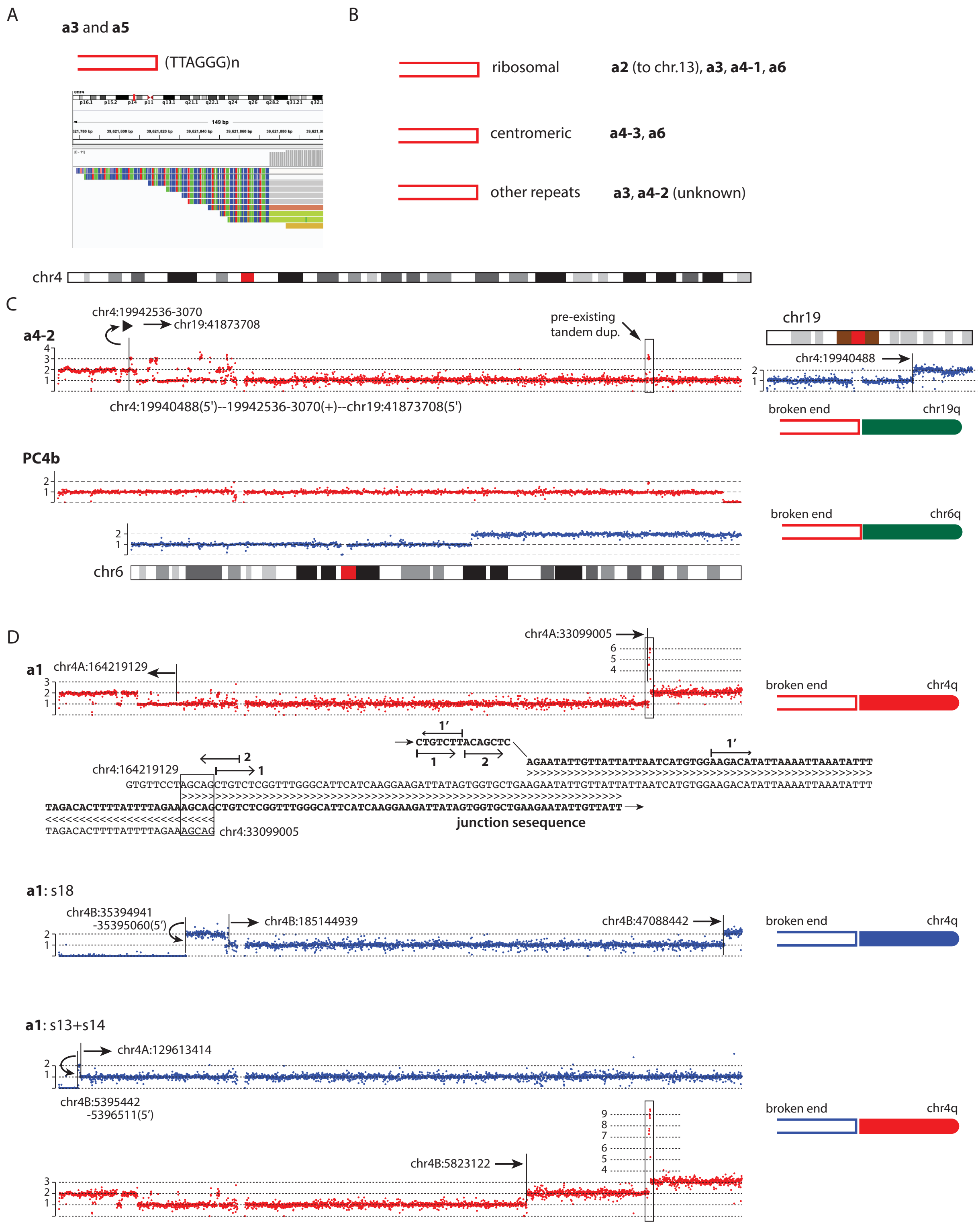

#### 6. Rearrangement outcomes of break-induced replication

Break-induced replication (BIR) is commonly assumed to be the mechanism of repair of unpaired DNA ends. For reviews of BIR, see Anand, Lovett, and Haber (2013), Verma and Greenberg (2016), and Kockler, Osia, Lee et al. (2021). Here we only discuss the rearrangement outcomes of BIR given the translocation outcomes of chromosome ends shown in the previous section.

**SI Figure 8: Breakpoint/rearrangement outcomes of break-induced replication**

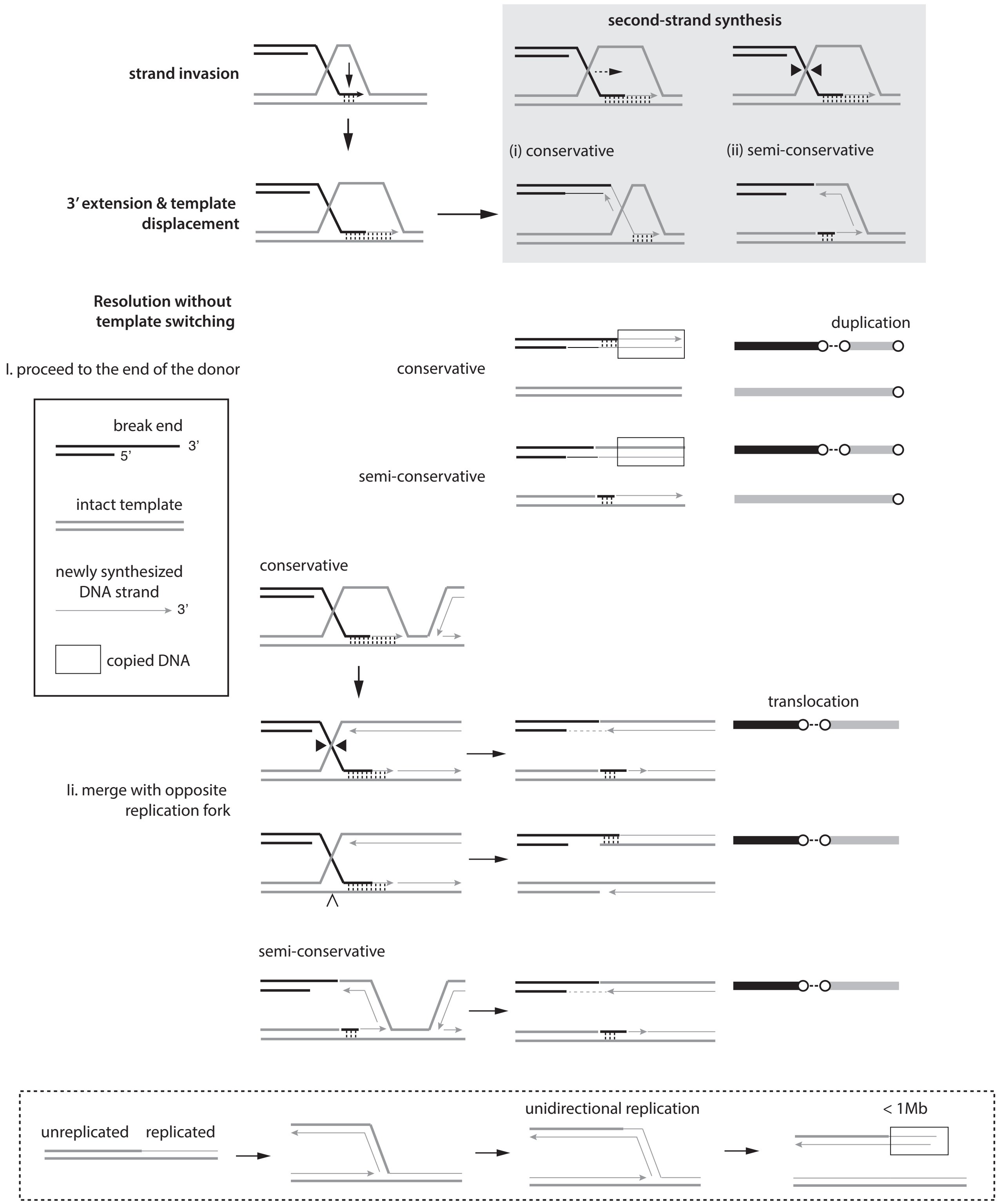


 

The first step of BIR is strand invasion. This step is mediated by homology-dependent pairing of a break end (black) and an intact donor template (gray). Strand invasion in BIR is similar to homologous recombination, both requiring significant homology between the break end and the donor. By contrast, microhomology-mediated BIR (MMBIR) is thought to require only limited homology (microhomology). We note that both homology-mediated and microhomology-mediated strand annealing can occur in other forms of DNA repair that join two DSB ends, including HR and SDSA (synthesis-dependent strand annealing), which requires extensive homology, and microhomology-mediated end-joining. Therefore, the junction homology feature is not a definitive signature of BIR or MMBIR.

After strand invasion, the 3'-end (arrowhead) is extended by DNA polymerases (Pol $\delta$ or its subunit *POLD3*). But how lagging-strand synthesis ("second strand synthesis") is carried out is largely unknown. Studies in yeast suggested BIR synthesis to be *conservative* (Saini, Ramakrishnan et al., 2013; Donnianni and Symington, 2013); however, it is unknown if the same applies to mammalian BIR or MMBIR. Moreover, it is unclear whether lagging-strand BIR requires some or all of the enzymes involved in normal lagging-strand synthesis, including Pol $\alpha$, Pol $\delta$, and ligase I. See Discussion in Donnianni, Zhou, Lujan et al. (2019). Both forms of second-strand synthesis (conservative and semiconservative) are considered here.

The original model of BIR suggests that the BIR fork can progress to the end of the donor chromosome. A recent study suggested the rate of BIR synthesis to be 0.5kb per minute (Liu, Yan, Osia et al. 2021). This is considerably lower than the rate of normal DNA synthesis or the extension rate of Pol $\delta$ ($\sim$ 4.3kb/min). Even at a rate of 1kb/min, it will take 1000 minutes, or 16.7 hours, to synthesize 1Mb of DNA. This cannot solely account for the large duplications (>10Mb) at the chromosome ends as shown in the previous section. Moreover, Liu et al. suggested that BIR progression is often interrupted by roadblocks such as the transcription unit. Based on these findings, we consider it *highly unlikely* that BIR synthesis (including MMBIR) can produce large segmental duplications (>1Mb) or reach the end of the donor chromosome.

If BIR synthesis is not terminated at the end of the donor chromosome, it can be resolved in three possible ways.

The first is when the BIR fork comes to the end of the donor that is a chromosome fragment. This scenario can generate a duplication of the donor template whose size is within the processivity of a BIR fork ($\lesssim$100kb). This process also duplicates the end of the donor template.

The second is when the BIR fork merges with an opposite replication fork. See Smith, Lam, and Symington (2009) and Costantino, Sotiriou, Rantala et al. (2014). This scenario is only possible when the original DNA end invades an unreplicated donor. It will generate a translocation between the original DNA end (black) and a sister fragment of the donor chromosome. Notably, this scenario will create a donor chromosome that consists of both unreplicated (left) and replicated (right) DNA; the firing of replication origins within the unreplicated DNA will create an unpaired DNA end on the donor chromosome and may lead to re-replication (bottom dashed box).

The third possibility is template switching. Template switching can occur if the invading strand is displaced from the donor, which will create an insertion outcome.

Based on the above discussion, the translocation outcome in **SI Figure 7D**, **a1:chr4A** is consistent with a BIR/MMBIR invasion, which creates short insertions ($\sim$10bp) at the site of invasion, followed by merging of the BIR fork with an opposite replication fork. The other outcomes in **SI Figure 7C** and **7D** may arise from a similar mechanism or by end-joining between the end of rearranged chr4 with DNA segments generated by secondary events.

**SI Figure 9: Segmental structure of rearrangement of chr11 in the X-29 clone from Maciejowski et al. (2015)**

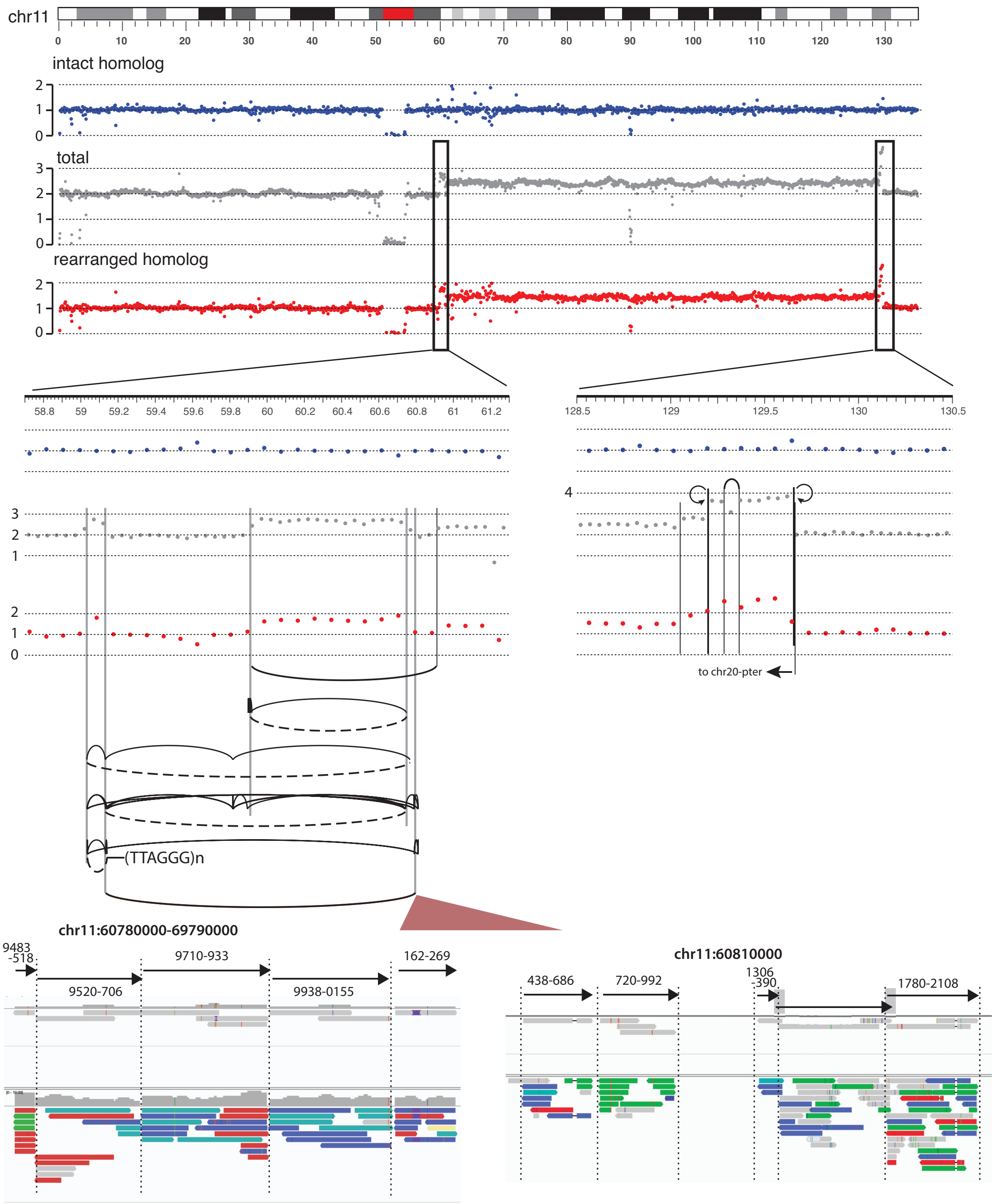


*Top*: Haplotype-specific and total DNA copy number of chr11 in the X-29 clone.
*Middle*: Zoomed-in view of copy-number transitions near 60Mb and 130Mb.
*Middle Bottom*: Rearrangement junctions at 60Mb and 130Mb, shown similarly as in **Extended Data Figure 5**.
*Bottom*: Two hotspots with tiling insertions near 60.78Mb, and 60.81Mb. The top IGV snapshots show contigs assembled from split and discordant reads shown in the bottom. Additional hotspots include 59.03Mb, 59.13-59.14Mb, 59.82Mb, 59.90-59.91Mb, 60.74Mb, and 60.79-60.81Mb. Insertions near 59.03Mb and 59.13Mb are shown on the next page.

Although it is impossible to determine the segmental structure of duplications based on bulk sequencing data alone, we can draw inferences about the structure of duplicated segments based on DNA copy-number transitions and rearrangement junctions. This analytical strategy may be applicable to standard bulk sequencing of cancer genomes.

For the copy-number calculation, we use the total coverage to derive more accurate quantification. This is feasible because all copy-number transitions occur to a single parental chromosome based on the haplotype-specific coverage. The copy-number gains are approximately 0.8x (59.03-59.13Mb and 59.9Mb-60.74Mb) and 0.4x (60.76-60.79Mb and 60.9Mb-130Mb) and the copy-number transitions are associated with the following breakpoints: paired parallel breakpoints at 59.03Mb(+), 59.13Mb(-), 59.91Mb(+), and 60.74Mb(-); single breakpoints at 60.77Mb(+), 60.80Mb(-), and 60.90Mb(+). For single breakpoints, their copy number is determined by the copy-number difference (~0.4). Based on the junctions (red dashed lines for junctions with insertions, solid line for simple junctions), we further determine that all the breakpoints have the same copy number ~0.4.

We can draw the following inferences about rearranged segments.

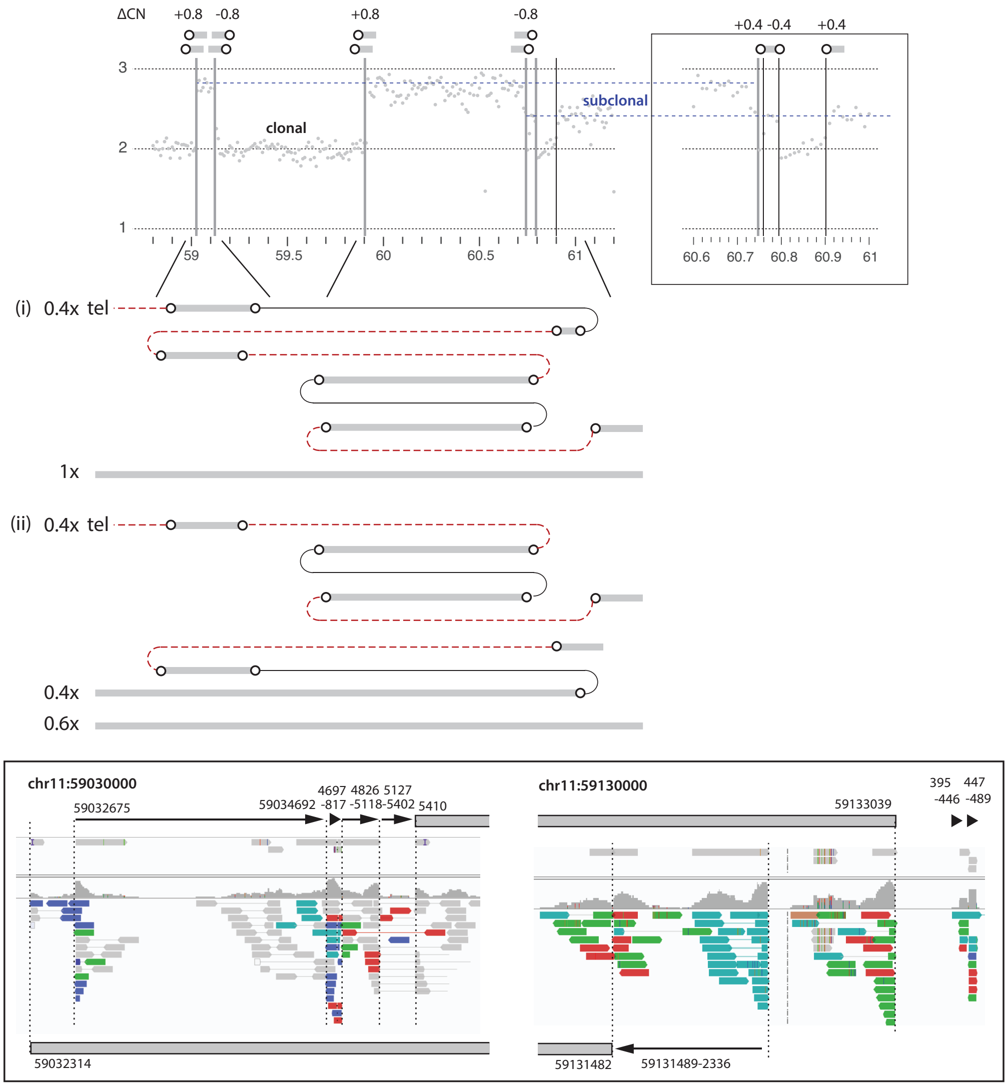


First, the gains of 59.03-59.13Mb and 59.91-60.74Mb are contained in two sister segments generated by replication. This inference is based on the presence of adjacent parallel breakpoints on both sides of the duplications. (Note that the same conclusion may not be drawn for duplications flanked by flush breakpoints and adjacent breakpoints. See Evolutionary analysis of *trans* segments and breakpoints.) The association of parallel breakpoints with staggered dsDNA ends is further supported by the observation of non-overlapping insertions mapped to regions near all four copy-number transitions. (Insertions mapped to 59.03Mb and 59.13Mb are shown below.)

Second, because the breakpoints at 59,917,316 and 60,749,797 are joined together, they cannot be in *cis* as that will create an ecDNA circle that is unlikely to have the same copy-number state as the other segments. We note that this inference suggests a pairing of the shorter end on the left (59,917,316) with the shorter end on the right (60,742,114), which violates strand coordination (see **Extended Data Figure 4**). This could reflect an ancestral dsDNA end with a significant 5'-overhang, or a sister-chromatid exchange within the two segments.

We show two possible configurations of rearranged segments that are consistent with the above inferences.

**SI Figure 10: Copy-number variation and segmental structure of rearrangement in Primary Clone 5a from Umbreit et al. (2020)**

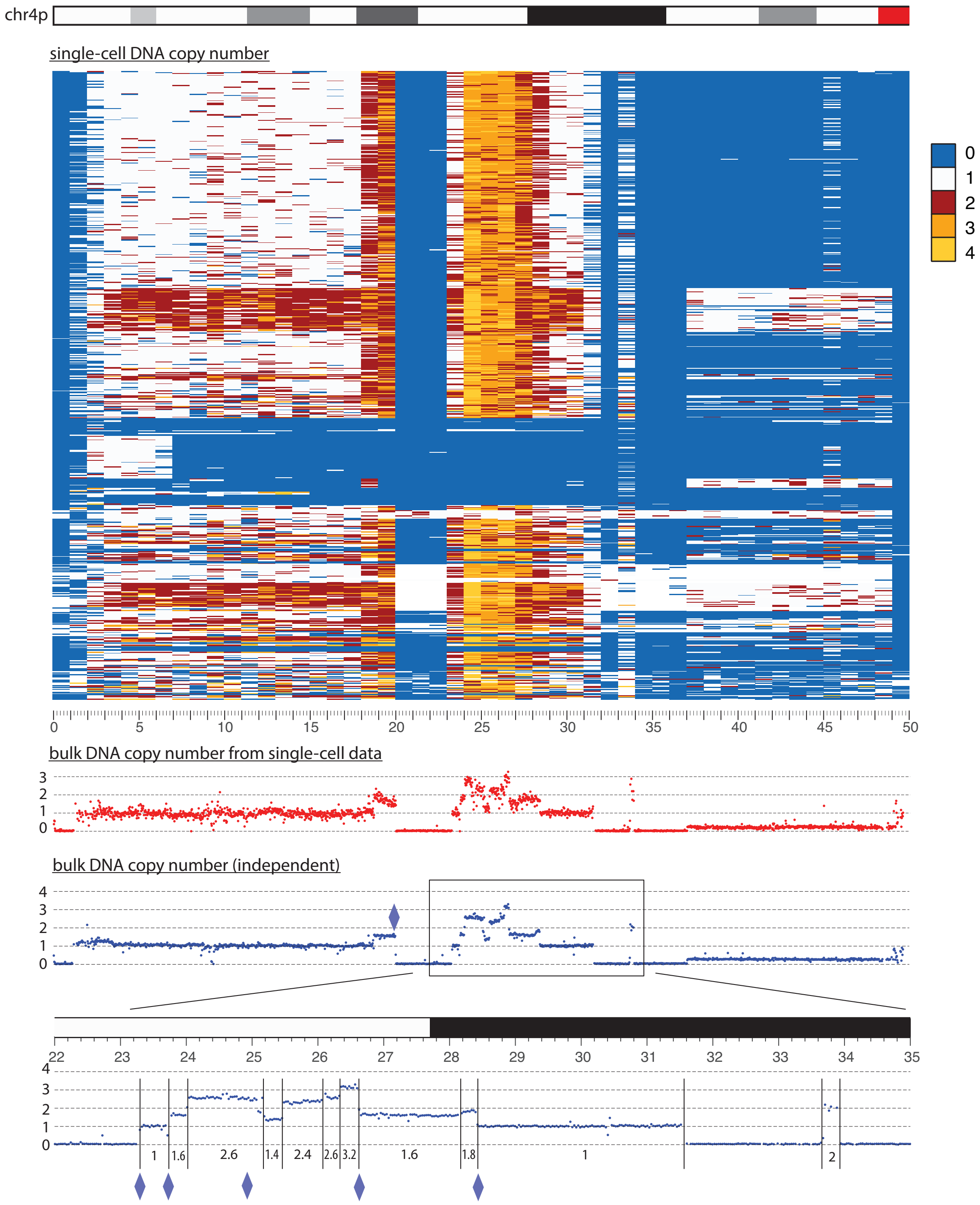


*Top*: Single-cell copy-number profiles of the altered haplotype.
*Middle (red)*: Bulk DNA copy number calculated from the single-cell data.
*Middle bottom (blue)*: Bulk DNA copy number calculated from bulk DNA sequencing.
*Bottom*: Zoomed-in view of copy-number transitions between 22 and 35Mb. Diamonds represent hotspots of mapped insertions. The hotspot at 25.71Mb has two reciprocal breakpoints (not shown).

We also want to draw inferences about the segmental structure of duplications based on DNA copy-number transitions and rearrangement junctions. First, we determine the copy number of breakpoints.

**Inference of breakpoint copy number based on segmental copy number transitions**

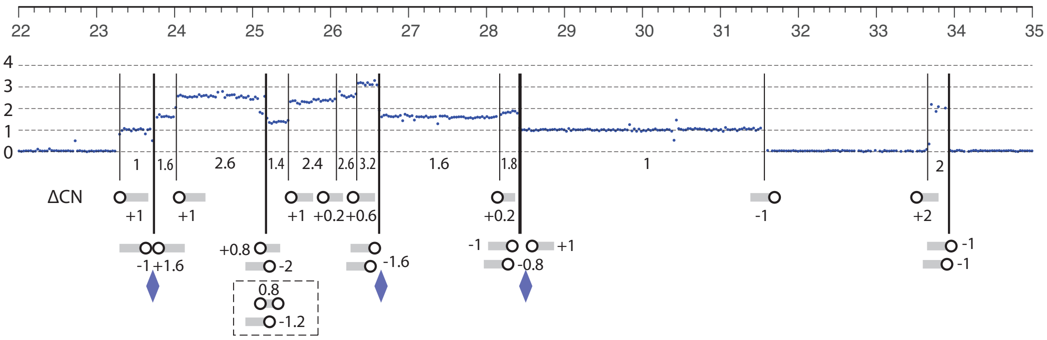


For unpaired breakpoints, their copy number states are determined from the copy-number difference:

- 23.29Mb(+1), 24.01Mb(+1), 25.46Mb(+1), 26.04Mb(+0.2), 26.33Mb(+0.6), 28.18Mb(+0.2), 31.59Mb(-1), 33.68Mb(+2).

We then use the junction information to determine the copy number of adjacent breakpoints. There are two adjacent gapped breakpoints at 25.014Mb(-) and 25.018Mb(+): based on the junction between 25.018Mb(+) and 23.29Mb(+1), we determine its copy number to be 25.018Mb(+1); because there is no net copy-number change at 25.01Mb, we further determine the breakpoint at 25.014Mb to be 25.014Mb(-1). From the junction between 25.014Mb(-1) and 23.72Mb(-), we determine the latter to be 23.72Mb(-1); finally, because the net copy-number change at 23.72Mb is +0.6, we determine the copy number of 23.73Mb to be 23.73Mb(+1.6). Together, we have:

- 23.72Mb(-1), 23.73Mb(+1.6), 25.014Mb(-1), 25.018Mb(+1).

We next consider the copy-number transition at 25.17Mb with two breakpoints 25.170(+) and 25.173(-). The breakpoint 25.173(-) forms a junction with the breakpoint 33.68Mb(+2), therefore its copy number is 25.173Mb(-2); based on the segmental copy-number difference, we determine 25.17Mb(+) to be 25.17Mb(+0.8). Note that this inference is independent of the *cis* or *trans* relationship between the two breakpoints at 25.17Mb. The junction between 25.17Mb(+0.8) and 28.417Mb(-) then determines the latter to be 28.417Mb(-0.8). Finally, the copy-number transition at 28.42Mb has a net copy-number change of -0.8 that is equal to the copy number of 28.417Mb(-0.8); therefore, the two remaining breakpoints 28.423Mb(-) and 28.424Mb(+) must have the same copy number. Because these two breakpoints form junctions with 33.93Mb(-) and 33.94Mb(-), the copy number of the latter two breakpoints must be equal; therefore, the latter two breakpoints must each have copy number -1 to balance the copy-number gain at 33.68Mb(+2). Together, we have

- 25.170Mb(+0.8), 25.173Mb(-2), 28.417Mb(-0.8), 28.423Mb(-1), 28.424Mb(+1), 33.93Mb(-1), 33.94Mb(-1).

The total copy number of two breakpoints at 26.61Mb(-) and 26.62Mb(-) is -1.6 but the individual copy number state cannot be determined. There is a short internal deletion of 28,419,969-28,420,319 whose copy number cannot be determined.

The breakpoint copy number can be used to determine the structure of rearranged segments. For example, the breakpoint at 23.72Mb(-1) has to be in *cis* with the breakpoint at 23.29Mb(+1) because the flanking sequence to the left of 23.29Mb is deleted. The identical copy number of these two breakpoints further suggests that they are retained only in this segment. Similar reasoning can be applied to determine the presence of one unique segment between 28.42 and 31.59Mb and two unique (sister) segments between 33.68 and 33.93Mb. Moreover, the breakpoint at 28.18Mb(+0.2) has to be in *cis* with either 28.417Mb(-0.8) or 28.423Mb(-1).

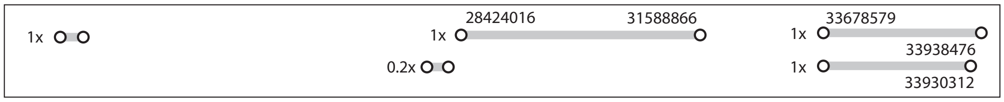


For the remaining segments, their breakpoints cannot be determined. Two possible configurations are shown on the next page. In both configurations, we have avoided the *trans* triplication configuration (right, top) and minimized partially overlapping segments (right, bottom).

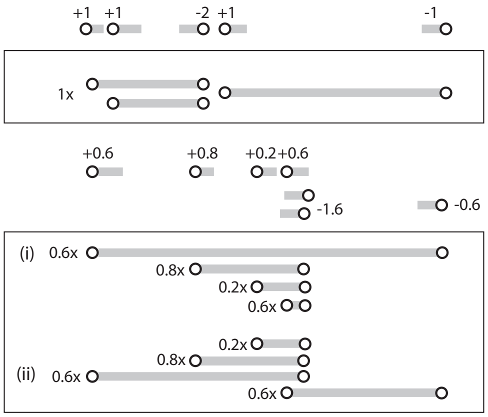
+1 +1 -2 +1 -1
1x
+0.6 +0.8 +0.2 +0.6
-1.6
-0.6
(i) 0.6x
0.8x
0.2x
0.6x
0.2x
0.8x
(ii) 0.6x
0.6x

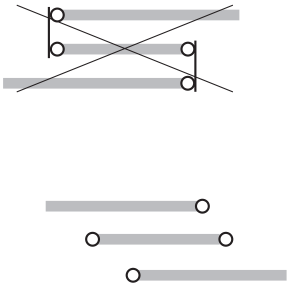

#### 8. Origin of insertions that share a breakpoint with large segments

**SI Figure 11: Kataegis within insertions and a proposed model for insertions that share a breakpoint with large segments.**

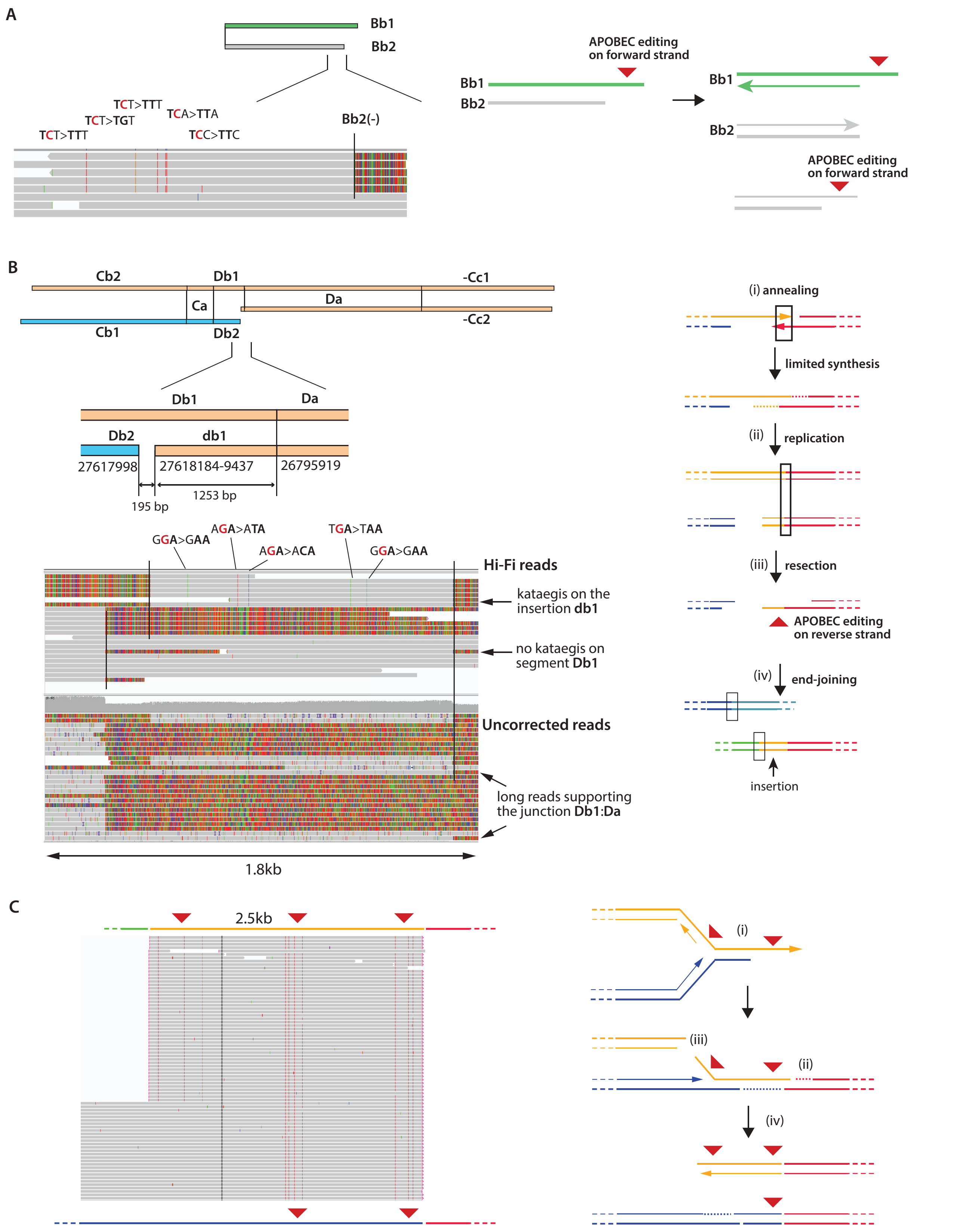

**A**. An example of kataegis at the shorter end of parallel breakpoints. Mutations indicating cytosine deamination on the forward DNA strand (C>T substitutions) are observed at the ends of both **Bb1** and **Bb2** segments. Since the ancestor of **Bb2** is the reverse strand DNA, deamination must have occurred after replication and resection.

**B**. *Left*: Structure of compound duplications consisting of segments **Cb1/Cb2**, **Ca**, **Db1/Db2**, **Da**, and **Cc1/Cc2**. The junction between **Db1** and **Da** is duplicated in two separate segments. Evidence for the presence of both **Db1:Da** junction and **db1:Da** junction is provided by long reads. Interestingly, there are five substitutions restricted to the insertion **db1** and having the signature of deamination on the reverse strand DNA.

*Right*: A proposed model for creating short insertions derived from sequences near dsDNA ends. (i) Two staggered DNA ends anneal with each other followed by gap filling synthesis; the top strand is fully ligated but the bottom strand has a gap due to hyperresection of the left DNA end (dark blue). (ii) DNA replication converts the gap on the bottom strand into dsDNA ends. (iii) Newly created ends are resected and undergo deamination. Note that the strand of deamination is opposite to the original DNA overhang. This is further supported by two substitutions (TGA>TCA and AGA>ACA) on the left end of **Da** that are restricted to the copy of **Da** linked to the short insertion **db1** but not the longer **Db1** segment. (iv) The newly created ends are ligated to other DNA ends (light blue and green), during which deaminated cytosines are converted into substitutions.

**C**. An example from the HCC1954 breast cancer genome where a 2.5kb (chr8:132,804,543-132,807,079) insertion has additional deamination than the large segment (bottom). The patterns of deamination with the same signature suggest there were two copies of the *same* ssDNA strand. A plausible model is shown on the right.

(i) Deamination occurs to ssDNA both near the dsDNA end and internally when the newly synthesized DNA (lagging strand) is resected. (This can also occur when the fork is reversed.)

(ii) Annealing and gap-filling synthesis create a junction and convert deaminated cytosines at the distal end to substitutions.

(iii) Cleavage of the top strand creates a 5'-ssDNA end.

(iv) A leftward replication fork from the right (red segment) creates the short insertion (orange) from the top strand that contains additional deamination; additional gap-filling synthesis and ligation create the bottom segment that only contains substitutions from the distal deamination. This model is similar to the model shown in **B** except that the bottom strand is ligated but the top strand is cleaved.

In both **B** and **C**, insertions are created from sequences in the overhang region of a staggered DSB end. In **B**, the left segment is unreplicated and has a resected 5'-end; in **C**, the left segment is replicated but resected end is not. In both examples, the first step is the annealing of the staggered DSB end with another DSB end on an unreplicated DNA segment. After annealing, the two strands of paired DSB ends are not fully ligated: As the replication fork from the partner segment (red) passes through the junction, an insertion junction is created from the unligated strand.

Insertions generated by the mechanism described above always share a duplicated (flush) breakpoint with a large segment. This feature is observed for the following insertions in bridge clone a: **bb2** (8kb, paired with **Bb2**), **cc2** (4kb, paired with **Cc2**) and **db1** (1kb, paired with **Db1**), and in the example shown in **Figure 6C** in the HCC1954 genome. Notably, the size of insertions generated by this mechanism is dependent on the size of the ssDNA overhang and the processivity of fill-in synthesis, which may reach 10-100kb. It is possible that segment **da1** (44kb, paired with **Da**) and **m1a1** (24kb, paired with **M1a**) are also generated by a similar mechanism.

#### 9. Origin of insertions mapped to regions with adjacent breakpoints

In addition to insertions that share a breakpoint with large segments, we can classify the remaining insertions identified in bridge clone **a** into five categories as shown on the right (**SI Figure 12**). In each category, the inserted sequences are either adjacent to each other (1st category) or adjacent to the breakpoints of large segments (2nd-5th category); both features suggest a direct connection between insertions and DNA breakage.

We envision several potential mechanisms that can generate insertions. The first is RNA or reverse transcribed cDNA, which was first described by Moore and Haber (1996) and Teng, Kim, and Gabriel (1996). The integration of these insertions at rearrangement junctions involves non-homologous end-joining. The second mechanism is by cleavage of ssDNA flaps, e.g., during microhomology-mediated end-joining repair. This mechanism could explain insertions that are adjacent to unpaired breakpoints or the longer end of staggered breakpoints. The third mechanism is by cleavage of ssDNA in R-loops. This mechanism could explain insertions originating in ssDNA gaps.

**SI Figure 13: Origin of insertions**

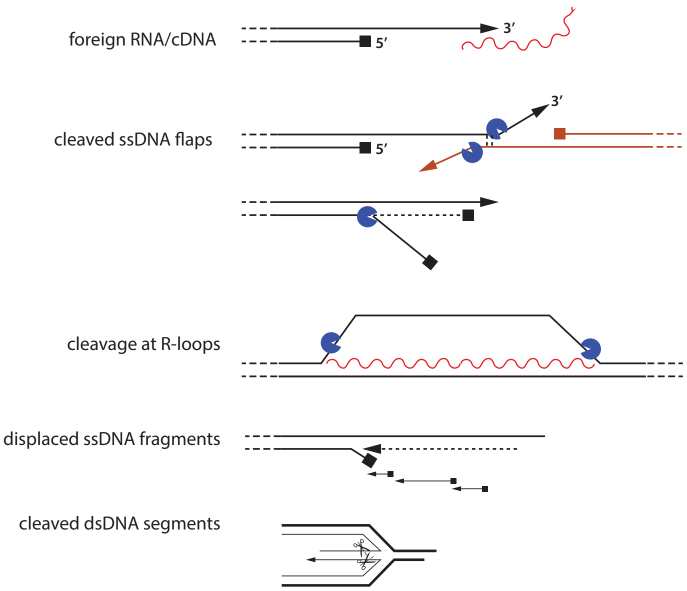


For insertions originating from the shorter end of staggered breakpoints, there are two possible mechanisms. The first is by displacement of pre-existing ssDNA fragments. The second is by cleavage of ssDNA or dsDNA in a reversed replication fork. The second possibility is disfavored because the resulting fragments can have gaps but not small overlap (<10bp) that is observed frequently (**Supplementary Table 2**). Such overlap most likely reflects some form of DNA synthesis with limited displacement activity. We therefore favor the first possibility that the fragments arise as discontinuous ssDNA fragments near the resected 5’-end of a staggered DSB end.

**SI Figure 12: Adjacency between insertions and segments**

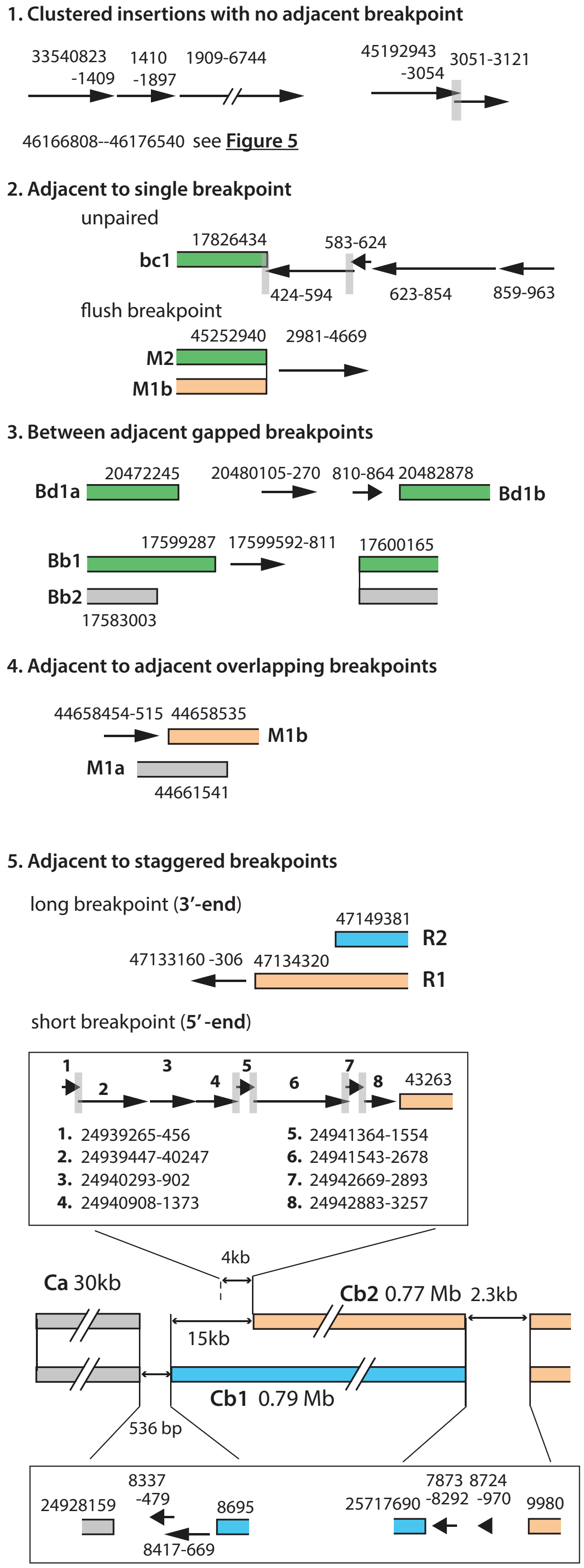

We next discuss several observations from the single-cell experiments that are informative about the mechanisms of insertions.

**SI Figure 14: Clustering of insertions in single cell T-1(a)**

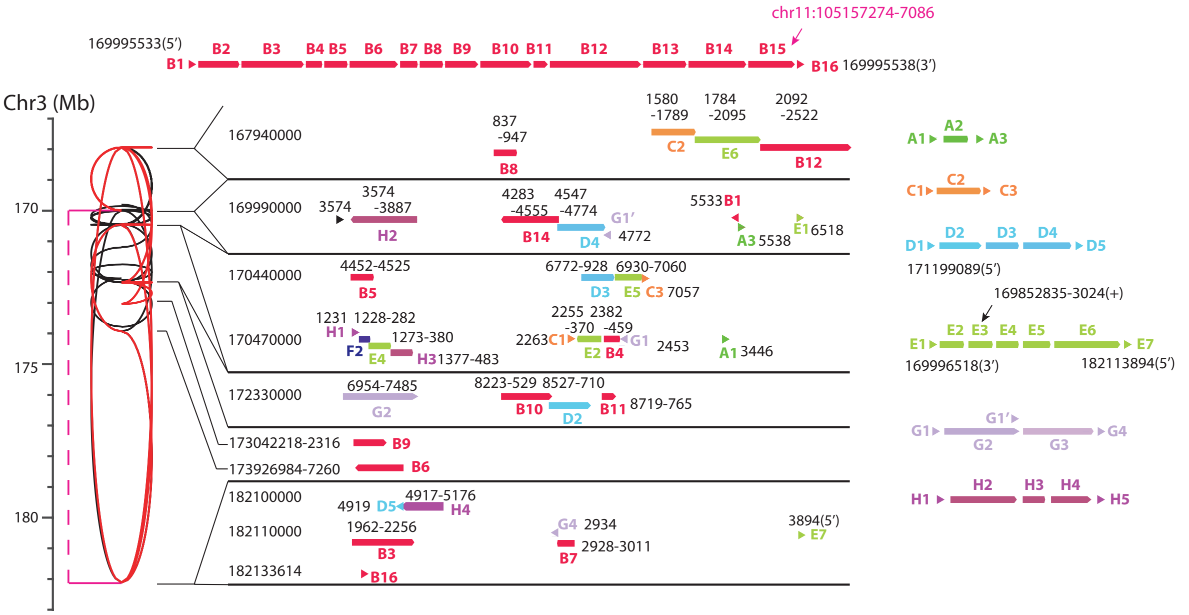


We identified chains of insertions in four single daughter cells with broken bridge chromosomes. For the daughter cell pair (**C-4**) shown in **Extended Data Figure 5**, bridge breakage occurred 3.5 hours after mitosis and cells were collected 24 hours after mitosis (by this timepoint the junctions were already formed). For cell **T-1(a)** that is shown above, bridge breakage occurred 12.3 hours after mitosis and cells were collected 22.6 hours after mitosis. For cell **C2(a)** shown on the right, bridge breakage occurred 3.5 hours after mitosis and the cell was collected 16 hours after mitosis. In all cases, we expect these daughter cells to be in G1 or early S phase when the bridge chromosomes were broken. Therefore, the initiation of DNA replication prior to bridge breakage is not required for the generation of junctions with tandem insertions.

Second, we never observed insertion junctions in cells with unbroken bridges and insertions are often mapped to regions adjacent to sites of chromosome breakage. In the **T-1(a)** daughter cell, insertions are mapped to several hotspot regions with adjacent breakpoints. In the **C-2(a)** daughter cell, we further identified insertions mapped to hotspots from non-bridge chromosomes. Together with observations in the bridge clones, these data suggest that insertions originate from sites of DNA breakage.

**SI Figure 15: Insertions in single cell C-2(a)**

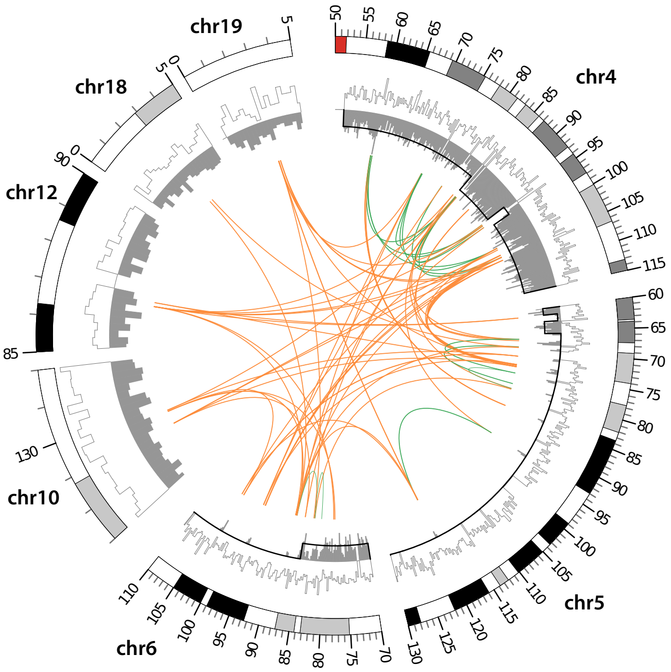


**Hotspot of insertions near chr12:86.696-86.701Mb**

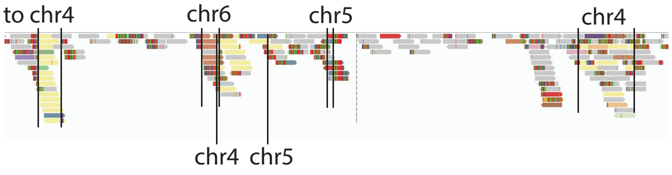

Finally, we never observed insertion junctions in cells where the bridge was mechanically broken and cell collection took place right away. We also did not detect insertion junctions in cells with micronuclei, although insertion junctions are prevalent in daughter cells that inherit damaged chromosome from micronuclei. Therefore, we infer that DNA breakage is required but insufficient to generate insertions.

The observed features of insertions (**SI Figure 16**) strongly disfavor the MMBIR model that postulates insertions arise from BIR-type synthesis ("copy-and-paste") when a DNA end sequentially invades and switches between multiple intact donors. Even if we assume the insertions arise from strand invasion into ssDNA donors (**SI Figure 17**), the tiling pattern of the inserted sequences (i.e., gapless with little/no overlap) cannot be explained by random strand invasion events. Moreover, if insertions are derived from DNA synthesis using ssDNA template, the newly synthesized DNA should show strand coordination: The insertions should originate from 5'->3' ssDNA fragments that are complementary to the donor template (the 3'-overhang), and should also be added sequentially to the 3'-end of the original DNA end. The presence of many instances of insertions with opposite orientation (strand alternation) in the same junction as shown in **Figure 5** excludes the MMBIR interpretation of insertions observed in our data.

**SI Figure 16: Features of short insertions observed in the experimental data**

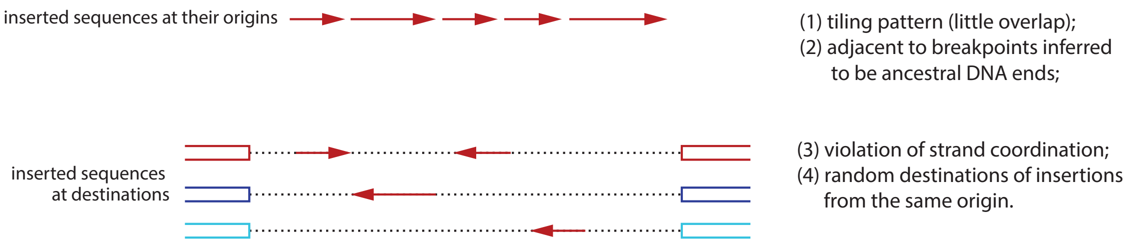


**SI Figure 17: Features of short insertions predicted by copy-and-paste or end-joining models**

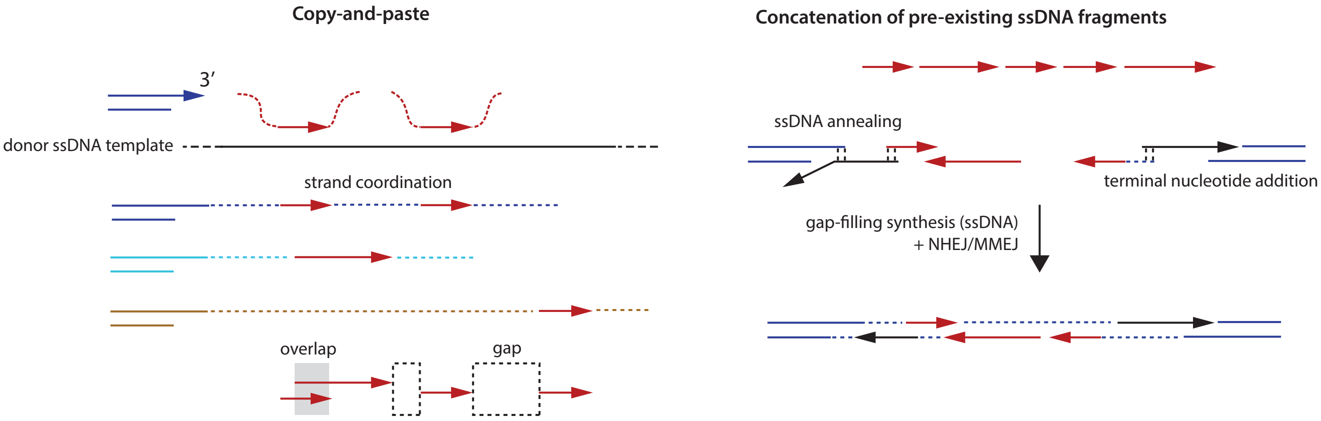


Both the tiling pattern of insertions at their origins and the strand alternation between insertions at the destination junctions can be explained if the insertion junctions are generated by concatenation of pre-existing ssDNA fragments (**SI Figure 17**, right). Direct annealing between ssDNA fragments based on microhomology will create insertions with strand coordination. ssDNA annealing can also prime DNA synthesis that coverts ssDNA fragments into dsDNA fragments, which can then be ligated at either orientation. Finally, gap-filling synthesis and ligation of ssDNA ends or dsDNA ends create a junction between distal dsDNA ends (blue) containing multiple insertions originating from ssDNA fragments from one or multiple origins.

Finally, we want to discuss potential mechanisms that can produce short insertions mapped to the 'overhang' region of parallel breakpoints, which we infer to be ssDNA fragments near the 5'-ends of resected DSBs. A plausible mechanism for the creating of these ssDNA fragments is by fill-in synthesis mediated by CST (CTC1, STN1, TEN1) and its associated DNA polymerase $\alpha$/primase. See the review by Mirman, Cai, and de Lange (2023).

CST/Pol $\alpha$/primase mediated fill-in synthesis occurs at both deprotected telomere ends and non-telomeric DSB ends. However, both the outcome of fill-in synthesis and its roles on DNA repair remain largely unknown. The first question is where fill-in synthesis starts. In vitro experiments suggest that CST preferentially binds ssDNA at the junction with dsDNA (Bhattacharjee,

Wang, Diao et al., 2017). If this were true in vivo, fill-in synthesis would preferentially start from regions near the 5'-ends of resected DSBs, which is consistent with the observation.

The second question is how long the fill-in product can extend and whether its extension involves a long-processivity polymerase such as Pol $\delta$. If the ssDNA fragments indeed originate from fill-in synthesis, our observation suggests that these fragments can reach 100-1000 base pairs that will require Pol $\delta$. Moreover, the presence of multiple discontinuous fragments with occasional small overlaps (less than 10bp) resembles unligated Okazaki fragments (Liu, Hu, Wang et al., 2017). This similarity is also reflected in their similar sizes (100-1000bp). These observations suggest that fill-in synthesis may generate discontinuous ssDNA fragments by a similar hand-off from Pol $\alpha$ to Pol $\delta$. As the processing of Okazaki fragments requires both ligation (by LIG-I or LIG-III) and flap removal (by the endonucleases FEN1 or DNA2), the persistence of these fragments may be explained by the deficiency of these enzymes (e.g., in an abnormal nuclear environment such as micronuclei or bridges). The observation that DNA2 deficiency promotes the generation of insertion junctions in budding yeast (Yu, Pham, Xia et al., 2018) is consistent with this explanation.

Discontinuous DNA synthesis near the 5'-end can fulfill a couple of roles during DNA repair. First, it counteracts 5'-resection. If the fill-in product reaches the 3'-end, it will create a blunt or near blunt end that is eligible for c-NHEJ or MMEJ. Second, it reduces ssDNA that is prone to deamination or cleavage (e.g., during MMEJ). Third, it provides a backup mechanism for lagging-strand synthesis during break-induced replication.

We propose a speculative model for the generation of short insertions as shown in **SI Figure 18**.

**SI Figure 18: A conjecture for the generation of short insertions originating from regions near resected DSB ends.**

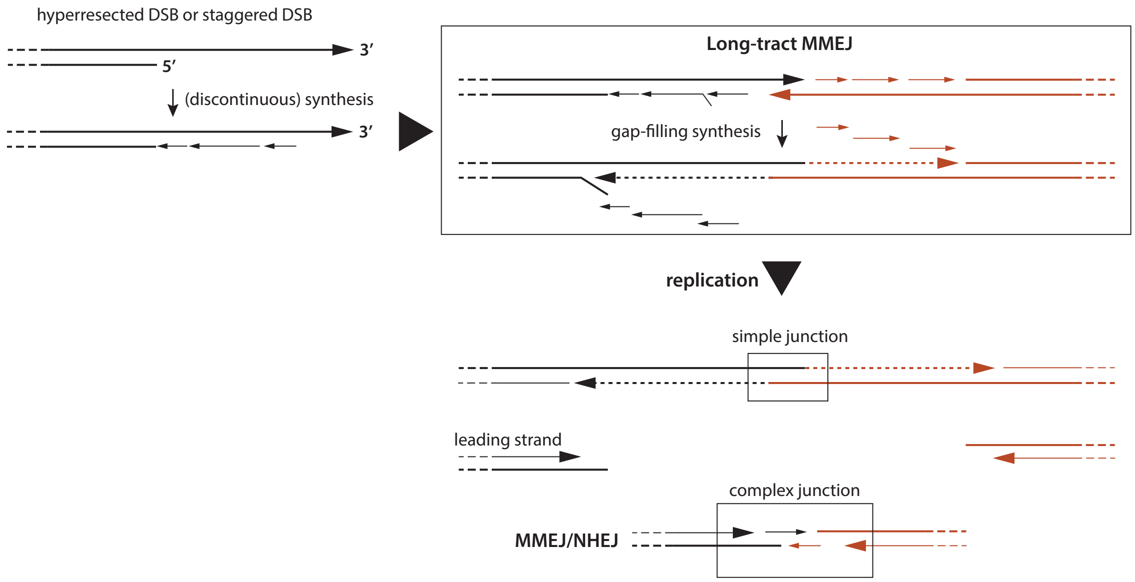

#### 10. Clusters of adjacent parallel breakpoints and foldback junctions

In **Extended Data Fig. 8B-E**, we show four examples of regions with clustered ("nested") foldback junctions. The three foldback junctions at 39.3Mb on chr8p (**Extended Data Fig. 8B**, recapitulated below) are on the telomeric side of the focal amplification that has three additional foldback junctions on the centromeric side.

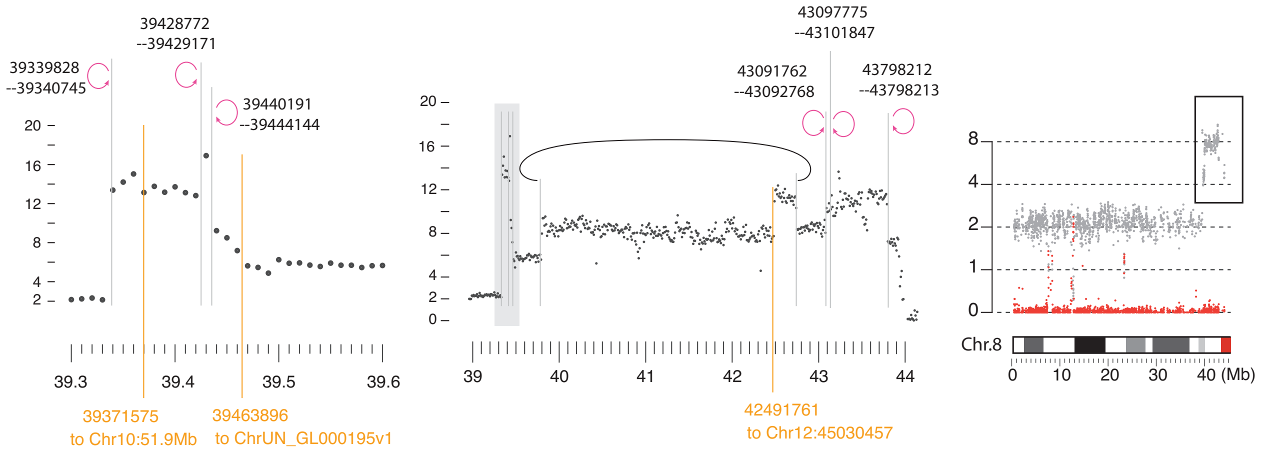


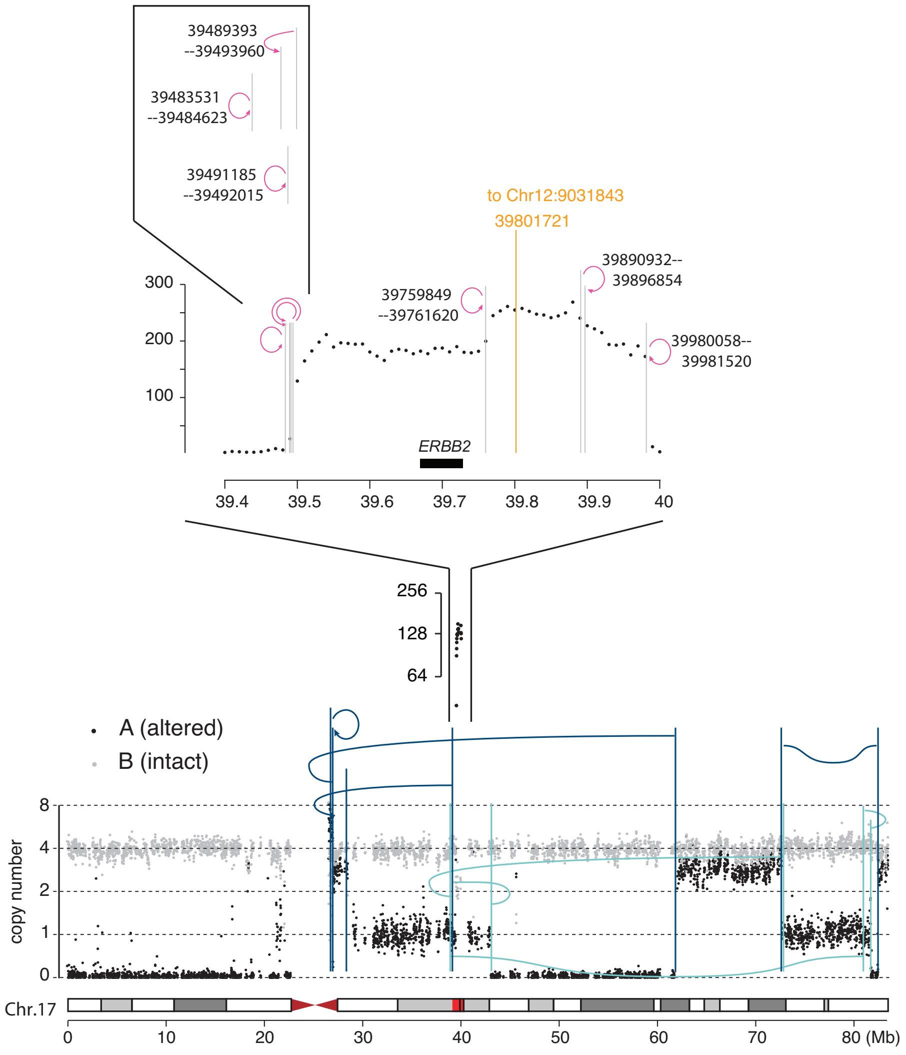


Clustered foldback junctions have two features: First, they are often located within 1Mb. The example from chr20 shows six foldback junctions within 3Mb, but three of them are within 0.5Mb; the example from chr12 shows five foldback junctions within 1.5Mb; in the most extraordinary example, there are three foldback junctions within 10kb near 39.5Mb of 17q. Second, adjacent foldbacks can either have identical (e.g., two foldbacks near 34Mb on 12p, three foldbacks near 39.5Mb on 17q) or opposite orientations (e.g., 8p, 20q).

If the foldback junctions are generated by sister-chromatid fusions over multiple breakage-fusion-bridge cycles, the breakpoints of different foldback junctions should correspond to sites of bridge breakage in different BFB cycles. To generate such clusters of breakpoints by random breakage events is highly unlikely. Assume bridge breakage occurs randomly between the two centromeres (spanning 10-20Mb), the probability to break at the same locus (<1Mb) three times is less than $(1/10)^2 = 0.01$. In the most extraordinary case where three foldback junctions are located within

10kb (39.483-39.491Mb). The probability to generate breaks at such proximity within the amplicon (0.5Mb) is less than $(0.01/0.5)^2 = 0.0004$. The regional concentration of foldback junctions also cannot be explained by positive selection since the *ERBB2* gene is in the middle of the amplicon. Moreover, there is no bona fide oncogene in the focally amplified regions on chr8p or 12p. (The region from chr20 contains *GNAS* that is suggested to be an oncogene in some cancers, but not in breast cancer.)

Given the mechanistic interpretation that foldback junctions arise from fusions between sister DNA ends, the concentration of foldback junctions implies a concentration of dsDNA ends. But a single dsDNA fragment can have at most two dsDNA break ends. How can a regional amplification of dsDNA ends occur? We suggest this can be accomplished by iterative B-R/F cycles (**Extended Data Figure 10**, recapitulated below).

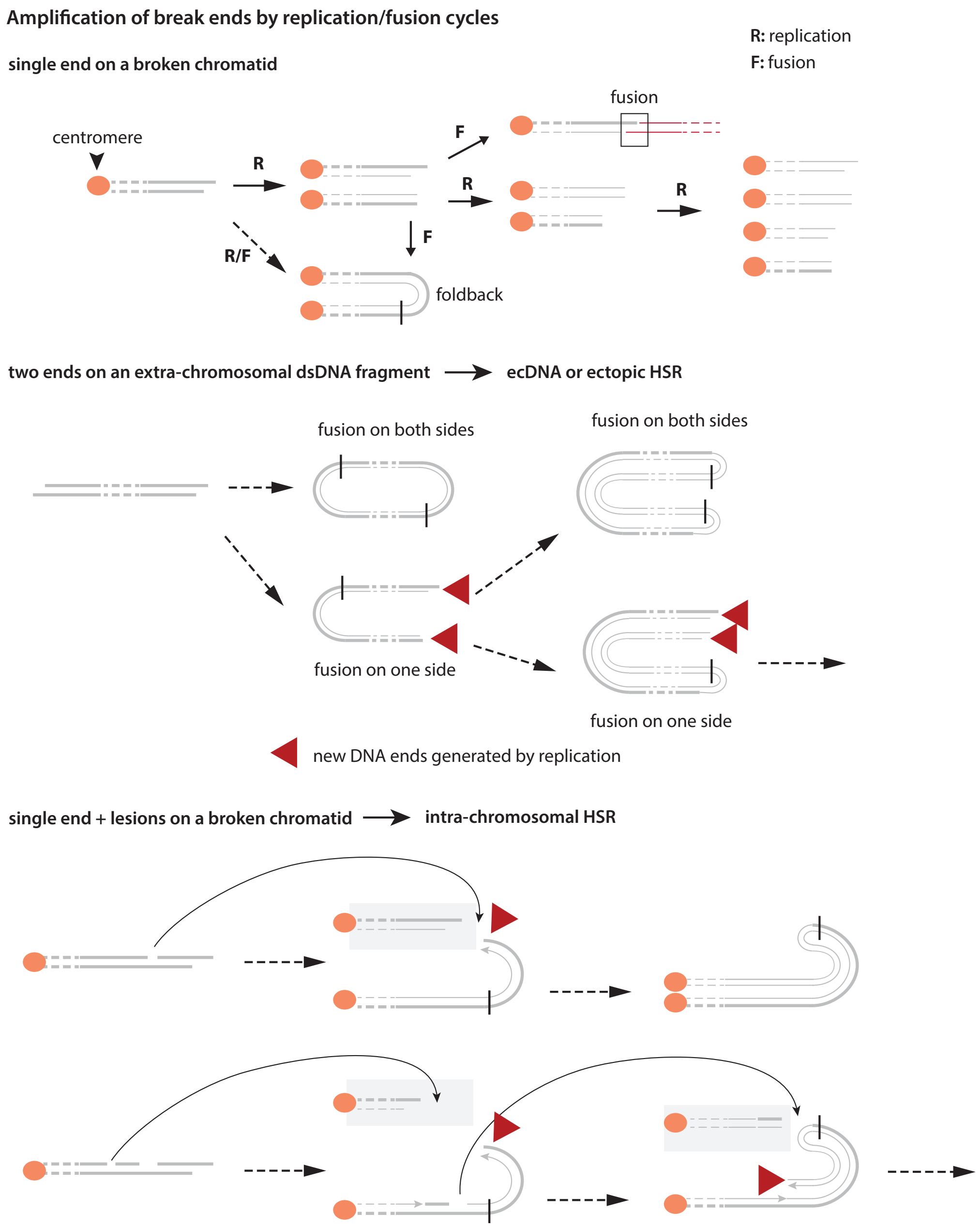


From a single dsDNA end on a broken chromatid, one round of replication creates two sister ends. If the two ends are fused together, they will form a foldback junction. In rare circumstances, the sister DNA ends can remain unligated and persist over another round of DNA replication; this can create four adjacent DNA ends. This provides a mechanism of DNA end amplification. However, the amplified DNA ends should normally segregate into different daughter cells. So the next question is how the amplified DNA ends and their junctions are preserved in a clonal population expanded from a single cell.

One plausible mechanism is when the replicated DNA fragments are linked by fusions on the opposite side (‘**extra-chromosomal**’). If both sides of the dsDNA fragment form foldback junctions at the same time, the outcome is a circular chromosome with foldbacks; this was previously recognized as “type II episomes” (Wahl, 1989) in contrast to “type I episomes” with the head-to-tail type junctions by a direct ligation of the two ends. However, if one side forms a foldback junction (left) but the other is replicated but not ligated (red arrows), the replicated DNA becomes a linear segment with inverted duplications; subsequent end fusions can generate further amplification and create a large, linear array of amplified DNA with only foldback junctions. The predicted outcome of this process is consistent with the structure of the amplification on 17q; this model is further supported by the observation that the amplified region on 17q does not overlap with other segments on 17q, indicating an ancestral chromothripsis event that produced the ancestor amplicon. For further analysis of 17q segments, see Brunette et al. (2024).

In the conventional model of double-minute (DM) amplification, DNA amplification occurs by *random* segregation of replicated acentric circular chromosomes (DM’s). In contrast to this model, DNA amplification by end-fusion will *autonomously* create a large linear amplicon consisting of DNA copies arranged in tandem but at *inverted* orientation. Eventually, the amplicon can either form a large circular chromosome or integrate into a linear chromosome to create an ectopic homogeneously staining region. For the *ERBB2* amplicon, the amplified DNA forms an isochromosome that is capped by telomeric segments from 12p. We suggest this process to be the mechanism of high-level focal amplifications with nested foldbacks.

When multiple DNA fragments are generated at the same by chromosome fragmentation (chromothripsis), these fragments can also fuse together into a compound fragment before amplification. Amplification by the same process as described above will create an array of amplified DNA with inverted orientation, with each amplicon consisting of multiple ancestral fragments that are non-overlapping. An example of such amplification is the *BCR-ABL* amplification in K-562 cells described in a previous study from us (Tourdot et al., 2021).

We note that both random double-minute segregation and sister DNA fusion generate amplified DNA that is **extra-chromosomal**. Both types of ecDNA amplification can occur after the generation of ecDNA fragments, e.g., from chromothripsis. The outcomes of these amplification processes include both circular DMs and ectopic HSRs.

We can envision a different type of iterative B-R/F cycles that generate **intra-chromosomal** amplification at the end of a broken chromatid (bottom of the figure on the previous page). In the first example, an internal ssDNA gap is converted into two dsDNA ends (long curved arrow), one of which becomes a new foldback junction after replication/fusion. This process can create two foldback junctions on opposite sides. In the second example, the presence of more DNA lesions leads to further amplification of DNA ends that can create further DNA amplification. Given the observations of regional hypermutation and DNA fragmentation near the site of breakage of chromosome bridges (Maciejowski et al., 2015, 2020; Umbreit et al., 2020), we expect the terminal region of a broken bridge chromosome to have frequent ssDNA lesions or gaps. These observations provide support for the proposed model of DNA amplification by replication/fusion cycles of DNA ends generated in a single episode of DNA damage. Moreover, the concentration of DNA lesions near the broken ends of the bridge chromosome directly explains the concentration of foldback junctions in the focally amplified region, eliminating the need to explain the recurrence of breakage implicated in multigenerational BFB cycles.

#### 11. Copy-number gain and amplification after micronucleation

In our prior studies, we have shown that chromosomes in micronuclei can both undergo partial replication (replicated DNA shown in red) and acquire DNA breakage (both ssDNA gaps and dsDNA breaks). The distribution of the damaged, partially replicated MN chromosome between two daughters can have two outcomes. When the broken fragments are distributed into both daughters, the fragments retained by each daughter should be largely mutually exclusive, but some replicated fragments may be retained by both daughters (**SI Figure 19**, uneven segregation into both daughters). In our previous paper (Zhang, Spektor, Cornils et al., 2015), we estimated that 3% of segments from the broken chr3 were retained by both daughters of the MN4 mother cell. Alternatively, the MN chromosome may be completely segregated into one daughter, with a complete loss in the other.

**SI Figure 19: Copy-number outcomes generated by the segregation of a fragmented chromosome from a micronucleus**

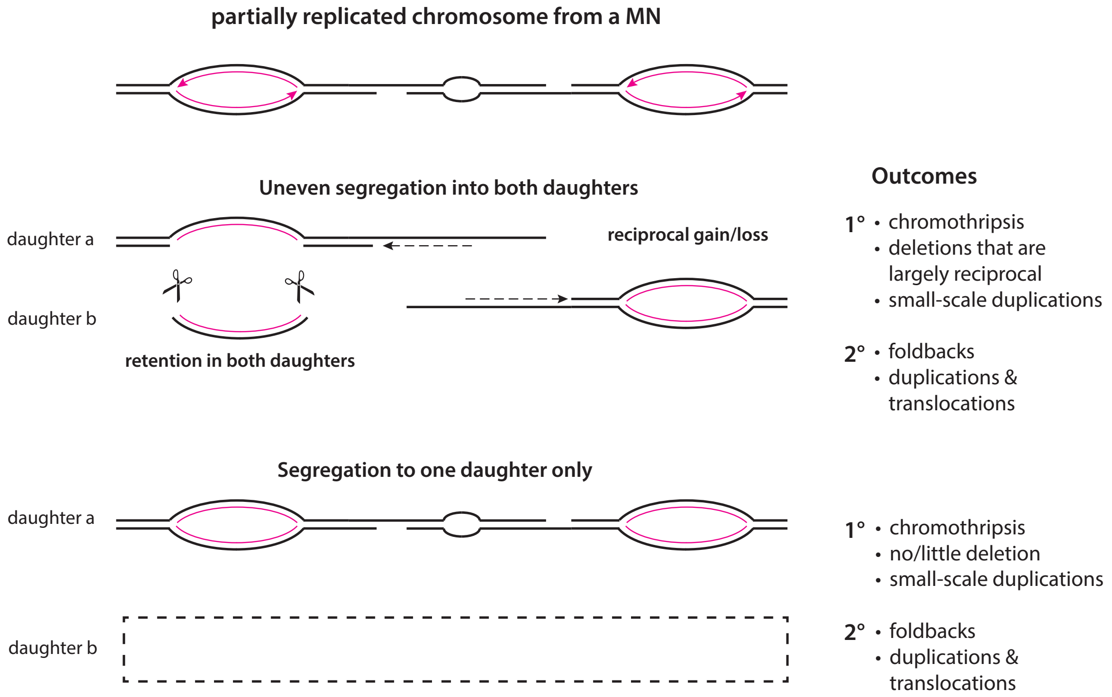


After the MN chromosome is reincorporated into the primary nucleus, a new round of replication can create dsDNA fragments from discontinuous ssDNA strands generated by partial replication (*left*). Fusions between these DNA ends can generate both long-range rearrangement junctions and foldbacks. As the MN chromosome can have up to four DNA strands at a given locus, after replication/fusion the MN chromosome can have up to four copy-number states: 0 (deletion from reciprocal distribution); 1 (ssDNA); 2 (dsDNA); 4 (replicated dsDNA). The segregation of the replicated MN chromosome can therefore have copy-number states from 0 to 4, not considering the rare instance of re-replication as shown in **Extended Data Figure 6**.

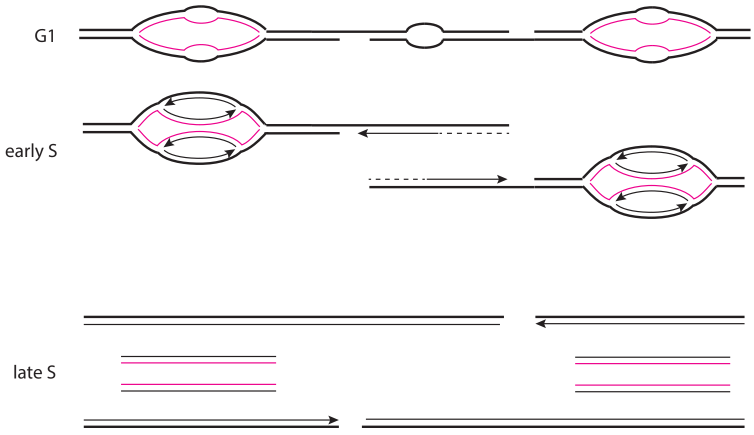

**SI Figure 20: DNA amplification from multiple generations of incomplete replication in MN**

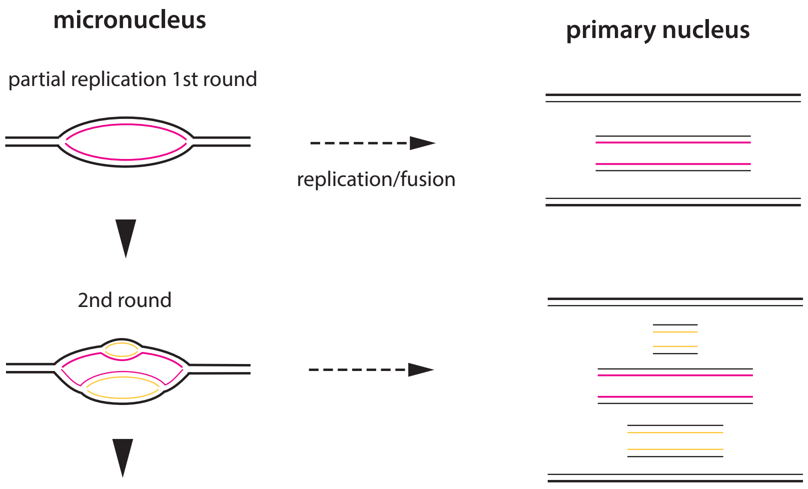


If the MN chromosome is partitioned into a newly formed micronucleus (persistent MN), the previously replicated region may undergo another round of partial replication (**SI Figure 20**). The eventual reincorporation of this chromatid into the primary nucleus will release all the partially replicated ss- and dsDNA fragments and lead to focal amplification. This mechanism is conceptually similar to the amplification of *Drosophila* amplicons in follicle cells (DAFC) reported previously (Alexander, Barrasa, Orr-Weaver, 2016). It provides a compelling explanation for the generation of complex amplifications on chr5, chr8, and chr21, where the concentration of breakpoints suggests a sequestration of these chromosomes, but the amplified copy-number states suggest multiple rounds of replication. A specific prediction of this model is that amplification should preferentially occur to sequences in *early*-replicating regions. This prediction can be tested in cancer genomes.
